# Neurodegeneration risk variants promote lysosomal TMEM106B fibril accumulation

**DOI:** 10.64898/2026.03.26.714634

**Authors:** John Michael Replogle, Jordan D. Marks, Martin G. Fernandez, Hebao Yuan, Dahyun Yu, Elizabeth Winters, Vidhya Maheswari Jawahar, Rishi Deshmukh, Renaldo Sutanto, Isabelle Kowal, Ashley Frankenfield, Rachel Shi, Yari Carlomagno, Karen Jansen-West, Tiffany W. Todd, Andrii Kopach, Iradukunda Sandra Ndayambaje, Yue A. Qi, Ananth Shantaraman, Ignacio Pozo-Cabanell, Udit Sheth, Mei Yue, Duc Duong, Shawn M. Ferguson, David A. Bennett, Markus Damme, Brad F. Boeve, Gregory S. Day, Benjamin Kellman, William C. Skarnes, Ronald C. Petersen, Keith A. Josephs, Neill R. Graff-Radford, Justin A. McDonough, Mercedes Prudencio, Sami J. Barmada, Yongjie Zhang, Ling Hao, Michael DeTure, Bailey Rawlinson, Erica Engelberg-Cook, Monica Castanedes Casey, Nathan Perez, Dennis W. Dickson, Aliza Wingo, Yue Liu, Nicholas T. Seyfried, Thomas S. Wingo, Shyamal Mosalaganti, Leonard Petrucelli, Michael E. Ward

## Abstract

Variants in *TMEM106B* and *GRN*, which encode lysosomal proteins, interact through unknown mechanisms to increase the risk of age-related cognitive decline and neurodegeneration. Here, we show that these variants converge on a single molecular intermediate: the cleaved intra-lysosomal fibril core of TMEM106B, a precursor to amyloid fibrils that accumulate in the aging brain. A protein-coding *TMEM106B* risk variant (p.T185) drives fibril core accumulation by impairing its degradation and *GRN* risk variants amplify this effect. Mice over-expressing the fibril core develop hallmarks of neurodegeneration, and cryo-electron tomography reveals intra-lysosomal fibrils in cultured neurons, mice, and diseased human brain. In *GRN*-mutation carriers, in whom fibril burden is greatest, fibrils extrude through ruptured lysosomal membranes. These findings identify intra-lysosomal TMEM106B fibrillization as a convergent neurodegeneration mechanism and potential therapeutic target.

---

Lysosomal function progressively declines with age in multicellular organisms (*1*–*4*). The lysosome degrades macromolecules, including protein aggregates too large or structurally complex for the proteasome (*5*). In most neurodegenerative diseases, pathogenic amyloid fibrils accumulate as intra- or extracellular deposits in the brain (*6*–*9*). When lysosomal function is compromised, the degradative capacity of the cell declines and fibril burden increases. This problem is compounded in post-mitotic neurons, which cannot dilute aggregates through cell division. Fibril build-up within dysfunctional lysosomes can then damage lysosomal membranes, creating a self-reinforcing cycle of protein aggregation and lysosomal failure (*10*).

Genetic studies have strongly linked compromised lysosomal function to neurodegeneration. Protective variants in the endolysosomal pathway associate with cognitive resilience in centenarians (*11*), whereas deleterious variants substantially increase the risk for age-related neurodegenerative diseases, including Alzheimer’s disease (AD), Parkinson’s disease, and frontotemporal lobar degeneration (FTLD) (*4, 12*–*14*). Two of the strongest genetic modifiers of both cognitive resilience and neurodegeneration risk are interrelated lysosomal genes: *GRN*, which encodes the intra-lysosomal protein progranulin, and *TMEM106B*, which encodes a lysosomal transmembrane glycoprotein (*15*–*20*). Heterozygous loss-of-function mutations in *GRN*, which reduce progranulin levels by half (*21, 22*), often cause FTLD with TAR DNA-binding protein 43 (TDP-43) inclusions (FTLD-TDP). Yet the penetrance, age of onset, and clinical progression of FTLD-TDP in *GRN* mutation carriers depend on co-inheritance of a risk haplotype in *TMEM106B* (*23*–*25*). This intimate relationship between *GRN* and *TMEM106B* is one of the most compelling examples of genetic interactions in neurodegenerative diseases. However, despite robust genetic evidence implicating a functional link between *GRN* and *TMEM106B*, the molecular mechanism by which their variants converge to drive neurodegeneration is unknown.

Recent studies identified TMEM106B amyloid fibrils in the aging brain and in multiple neurodegenerative disorders, including FTLD and AD (*26*–*29*). Lysosomal proteases cleave full-length transmembrane TMEM106B, releasing a luminal C-terminal fragment (*30*) that is further processed into the fibril core (TMEM106B^FC^), the precursor of amyloid fibrils (*26*–*30*). The sole protein-coding variant in the *TMEM106B* risk haplotype – a serine to threonine conversion at residue 185 – falls within the fibril core domain. Fibril burden increases stepwise with risk haplotype dosage in FTLD brain (*31*–*33*), suggesting that the disease-related variant may directly modify fibril abundance. However, the mechanisms driving fibril accumulation and whether fibrils cause neurodegeneration remains unknown. Here, we explore how the *TMEM106B* protein-coding variant and concomitant progranulin loss converge to regulate TMEM106B fibril burden and the role of these fibrils in neurodegeneration.

## A risk-associated *TMEM106B* coding variant drives accumulation of TMEM106B^FC^

A recent protein quantitative trait locus mapping (pQTL) analysis of more than 1,000 postmortem human brains identified rs3173615 as the lead *cis-* variant associated with altered TMEM106B protein levels (fig. S1A and Table S1) (*34, 35*). This variant resides within exon 6 of the *TMEM106B* gene and encodes a p.T185S amino acid change. In European-ancestry populations, both alleles are common, but the p.T185 variant is most prevalent and associates with increased FTLD-TDP risk relative to the S185 variant (*15*).

The rs3173615 coding change resides in the intraluminal fibril core domain of TMEM106B (TMEM106B^FC^), which can be cleaved from the full-length protein (Fig. 1A). Once cleaved, TMEM106B^FC^ exists as a discrete protein species (fig. S1B) whose biological properties may differ substantially from those of the full-length protein. The prior pQTL analysis (*34, 35*) relied on standard MS-based proteomics, which measures total protein and does not distinguish full-length TMEM106B from cleaved TMEM106B^FC^. In contrast, peptide-level analysis of MS-based proteomics data can reveal genetic associations specific to individual protein domains (*36*). We therefore tested for associations between rs3173615 and peptides derived from either the TMEM106B N-terminus or fibril core domain in postmortem brain tissue from the Religious Orders Study and Memory and Aging Project (ROSMAP) cohort (Tables S2 and S3) (*35, 37, 38*). The fibril core domain, but not the N-terminus, associated with rs3173615 at both individual peptide and domain levels, with fibril core levels increasing additively with each copy of the risk-associated C allele (CC > CG >GG) (Fig. 1B, fig. S1C, and Table S3). Collectively, these results suggest that rs3173615 is specifically associated with the levels of TMEM106B^FC^ and not full-length TMEM106B protein in the aged human brain.

**Figure 1.**
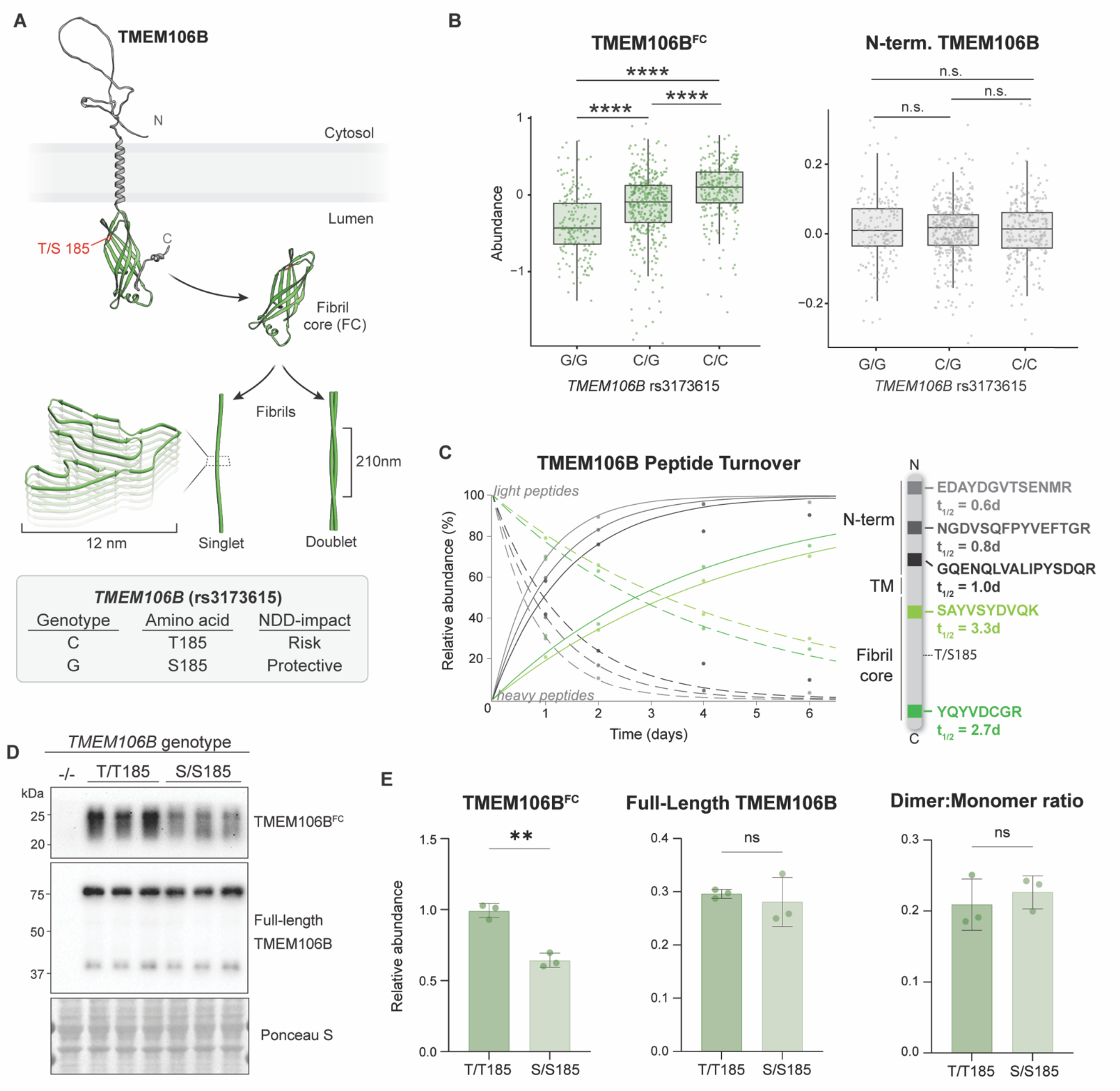
The risk-associated *TMEM106B* T185 coding variant selectively increases levels of TMEM106B fibril core. **(A)** Schematic illustrating the lysosomal localization of TMEM106B and its coding variant associated with neurodegenerative disease risk. TMEM106B is a lysosomal transmembrane protein whose luminal C-terminus is cleaved into its FC domain, a precursor of amyloid fibrils with mono- or doublet-fibril conformations. **(B)** *Cis* pQTL analyses of *TMEM106B* rs3173615 genotype versus TMEM106B N-terminal (gray, right) or C-terminal (green, left, ‘TMEM106B^FC^’) domain levels in the ROSMAP cohort. Statistics by one-way ANOVA with Tukey post-hoc. N = 146 G/G, 359 C/G, 218 C/C individuals. \**P* ≤ 0.05, \*\**P* ≤ 0.01, \*\*\**P* ≤ 0.001, \*\*\*\**P* ≤ 0.0001, ns: nonsignificant. **(C)** Plot of TMEM106B N-terminal (gray hues) and C-terminal (green hues) peptide half-lives. Time in days is plotted on the X-axis and relative abundance of light (dashed lines) and heavy (solid lines) peptides (as a % of baseline peptide abundance) is on the Y-axis. Intersections of same-color curves denote t_1/2_ of the represented peptide. Right: Schematic illustrating the location of plotted peptides with respect to domains of the full-length TMEM106B protein. Data from neuronprofile.com (Frankenfield *et al*. (2026)). **(D)** Western blot of endogenous full-length TMEM106B and cleaved C-terminal TMEM106B^FC^ in iNeurons differentiated from genetically engineered isogenic iPSC lines expressing TMEM106B (G/G; S/S185) or TMEM106B (C/C; T/T185). *TMEM106B* KO iNeurons were used as a negative control (leftmost lane). N= 3. (**E)** Quantification of panel D. Left: TMEM106B^FC^ normalized to Ponceau S. Student’s t-test. Middle: TMEM106B full-length (monomer+dimer sum) normalized to Ponceau S. Student’s t-test. Right: TMEM106B full-length dimer/monomer. Student’s t-test. Error bars = SD. \**P* ≤ 0.05, \*\**P* ≤ 0.01, \*\*\**P* ≤ 0.001, \*\*\*\**P* ≤ 0.0001, ns: nonsignificant.

To determine whether full-length TMEM106B and TMEM106B^FC^ have distinct degradation properties, we evaluated the half-lives of different peptides from TMEM106B using a recently published protein turnover atlas in human induced pluripotent stem cell (iPSC)-derived neurons (iNeurons) (neuronprofile.com) (*39, 40*). Endogenous N-terminal peptides have half-lives of less than one day, while C-terminal fibril-core peptides persisted for approximately three days (Fig. 1C). We then replicated this difference in iNeurons overexpressing full-length TMEM106B via dynamic stable isotope labeling by amino acids in cell culture (dSILAC) proteomics (fig. S1D). These two domains, therefore, exhibit distinct degradation kinetics that may allow for phenotypes or associations unique to each domain.

In European-ancestry populations, which comprise the majority of the ROSMAP cohort (*37*), rs3173615 (p.T185S) is in strong linkage disequilibrium with multiple noncoding variants and a 3’ untranslated region (UTR) Alu element insertion across the TMEM106B risk haplotype (*41, 42*). Genetic association studies therefore cannot identify which variant within this haplotype drive accumulation (*38, 44*–*46*). We hypothesized that because p.T185S lies within the fibril core, it might directly modulate TMEM106B^FC^ abundance. We edited endogenous *TMEM106B* in isogenic iPSC lines using CRISPR/Cas9 to encode either homozygous T185 or S185, making the amino acid at position 185 the sole genetic difference between lines (“T/T185” and “S/S185”, respectively; fig. S1E). We differentiated these iPSCs into iNeurons and measured TMEM106B^FC^ and full-length TMEM106B protein levels using immunoblotting (Fig. 1D). This analysis revealed that iNeurons expressing the T/T185 variant had substantially higher levels of TMEM106B^FC^ than iNeurons expressing the S/S185 variant (Fig. 1E). In contrast, the coding variant had no effect on the levels of full-length TMEM106B (Fig. 1E), nor its dimer to monomer ratio (Fig. 1E). Thus, the risk-associated T185 variant selectively increases TMEM106B^FC^ abundance without altering full-length TMEM106B levels, consistent with the human brain *cis*-peptide QTL results.

### Cellular and mouse models of TMEM106B^FC^ accumulation develop intra-lysosomal fibrils

The abundance of insoluble TMEM106B fibrils is associated with the TMEM106B risk haplotype, patient survival, and TDP-43 pathological burden in FTLD-TDP (*31*), but the mechanistic role of these fibrils in disease is unclear. To assess whether TMEM106B fibrils are causal mediators of neurodegeneration, we aimed to develop an experimental system that permits their direct evaluation. Developing this model required overcoming certain technical hurdles: while endogenous TMEM106B^FC^ is present in iNeurons (Fig. 1), its abundance is low compared to the full-length protein (fig. S2A), and spontaneous formation of TMEM106B fibrils in cultured neurons has not been reported. Overexpression of full-length TMEM106B can increase TMEM106B^FC^ levels (*31*); however, it also non-physiologically alters multiple aspects of lysosomal biology (*43*–*45*), complicating efforts to disentangle the effects of fibril burden from that of artificial overexpression. More importantly, simple TMEM106B overexpression paradigms fail to mimic disease, since risk variants cause a selective accumulation of TMEM106B^FC^ rather than the full-length protein (Fig. 1B and D) (*31*). We therefore developed a new model to specifically overexpress lysosome-targeted TMEM106B^FC^. We leveraged phospholipase D3 (PLD3), a transmembrane lysosomal protein whose cleavage upon lysosomal delivery yields a stable C-terminal domain in the lumen and a rapidly degraded N-terminal fragment (*46*). We hypothesized that a chimera between the N-terminus of PLD3 and the C-terminus of TMEM106B (PLD3-TMEM106B^FC^) would efficiently deliver TMEM106B^FC^ into lysosomes (Fig. 2A). We found that expression of PLD3-TMEM106B^FC^ in iNeurons showed the expected cleavage (Fig. 2B) and lysosomal localization of TMEM106B^FC^ (fig. S2B). Matching the processing and modifications described for the endogenous fibril core (*30*), PLD3-TMEM106B^FC^ underwent glycosylation and additional trimming beyond the end of the fibril core domain (past residue 255) (fig. S2C and S2D). This yielded a final fragment similar in size to the fibril core produced by the overexpression of full-length TMEM106B (Fig. 2B). Thus, the PLD3-TMEM106B^FC^ chimera allowed us to interrogate cells with a high burden of TMEM106B^FC^ with similar localization and processing to endogenous TMEM106B^FC^.

**Figure 2.**
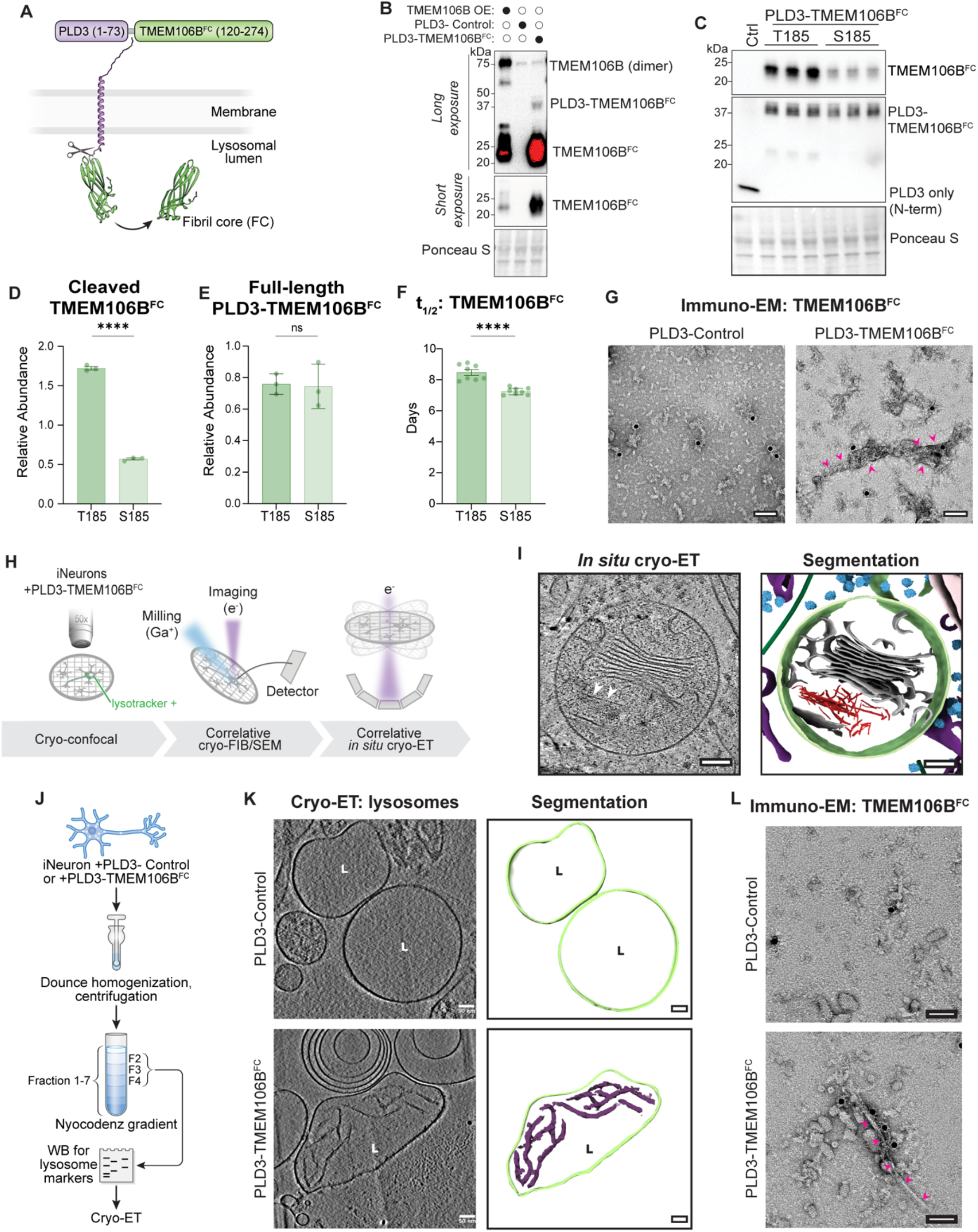
PLD3-TMEM106B expression drives accumulation of intra-luminal TMEM106B fibrils in neurons. **(A)** Schematic of PLD3-TMEM106B^FC^ chimera. Amino acids 1-73 of PLD3 were fused to amino acids 120-274 of TMEM106B via a flexible GS peptide linker to force cleavage of TMEM106B C-term and its release into the lysosomal lumen. Protein domain structures generated with AlphaFold 3. **(B)** Western blot using a C-terminal anti-TMEM106B antibody (AP26.2.3) from iNeurons overexpressing full-length TMEM106B (T185), PLD3-control (defined as: PLD3(1-73)–GSGSG without TMEM106B^FC^), or PLD3-TMEM106B^FC^ (indicated by solid black circle above lane). Ponceau S as loading control. **(C)** Western blot using a C-terminal TMEM106B (top) and an N-terminal PLD3 (middle) antibody from iNeurons expressing T185 or S185 versions of the PLD3-TMEM106B^FC^ construct, with a PLD3-control sample in the leftmost lane for comparison. **(D)** Quantification of TMEM106B fibril core from C, normalized to Ponceau S as a loading control. Student’s t-test. N= 3. Error bars = SD. **(E)** Quantification of PLD3-TMEM106B^FC^ chimera from C, normalized to Ponceau S as a loading control. Student’s t-test. N= 3. Error bars = SD. **(F)** dSILAC-derived half-life (t_1/2_) of median TMEM106B^FC^ peptide from iNeurons expressing PLD3-TMEM106B^FC^ (T185) or PLD3-TMEM106B^FC^ (S185). Student’s t-test. N= 8. Error bars = SD. **(G)** Representative negative stain electron microscopy of anti-TMEM106B^FC^ (AP26.2.3) immuno-gold labeled whole-cell sarkosyl-insoluble lysate from iNeurons expressing either PLD3-control (left) or PLD3-TMEM106B^FC^ (right). Black dots show gold particles. Arrowheads point to fibril. Scale bar: 50 nm. **(H)** Schematic of cryo-correlative light and electron microscopy (cryo-CLEM) and cryo-electron tomography (cryo-ET) workflow to identify lysosomal content in PLD3-TMEM106B^FC^ expressing iNeurons *in situ*. **(I)** Left: Slice through a reconstructed deconvolved tomogram, with white arrowheads pointing out intra-lysosomal fibrils. Right: Corresponding segmentation of tomogram in the left panel, highlighting pseudocolored cellular features – ER (purple), ribosomes (light blue), nuclear envelope (light pink), membrane stacks of unknown origin (gray), microtubules (dark green), lysosome (light green), and fibrils (red). Scale bar: 100 nm. **(J)** Schematic of lysosome enrichment and Western blot analysis from iNeurons via density gradient centrifugation. **(K)** Representative slices through the reconstructed deconvolved cryo-electron tomograms of lysosomes (‘L’) isolated from PLD3-control (top) or PLD3-TMEM106B^FC^ iNeurons (bottom). Fibrils are observed exclusively in PLD3-TMEM106B^FC^ lysosomes. Right: Corresponding segmentation of tomogram highlighting lysosome (light green) and fibrils (purple). Scale bar: 50 nm. **(L)** Representative negative stain electron microscopy images of anti-TMEM106B^FC^ immuno-gold labeled sarkosyl-insoluble lysosomal content isolated from iNeurons expressing either PLD3-control (top) or PLD3-TMEM106B^FC^ (bottom). Antibody against the C-terminal TMEM106B (AP26.2.3, black dots are gold particles). Arrowheads point to fibril. Scale bar: 100 nm.\**P* ≤ 0.05, \*\**P* ≤ 0.01, \*\*\**P* ≤ 0.001, \*\*\*\**P* ≤ 0.0001, ns: nonsignificant.

We next examined how the TMEM106B T185S variant affects the levels of TMEM106B^FC^ in our chimeric model. Consistent with endogenous TMEM106B, PLD3-TMEM106B^FC^ (T185) drove substantially higher abundance of the fibril-core, without altering expression of full-length PLD3-TMEM106B^FC^ (Fig. 2C-E). The levels of *PLD3-TMEM106B*^*FC*^ mRNA were also unaltered by expression of the T185 vs. S185 variant (fig. S2E), arguing that the coding variant alters TMEM106B^FC^ abundance at the protein level. We hypothesized two possible mechanisms by which the variant could mediate this effect: either by increasing TMEM106B^FC^ processing or slowing its degradation. If the PLD3-TMEM106B (T185) variant was more readily processed, we would expect to see elevated TMEM106B^FC^ levels alongside a concomitant decrease in levels of the full-length chimera. This was not the case (Fig. 2C-E). We therefore directly assessed the rate of degradation of TMEM106B^FC^ T185 or S185 in iNeurons expressing PLD3-TMEM106B^FC^ using SILAC pulse-chase proteomics. The T185 variant had a longer half-life than the S185 variant (Fig. 2F), while an N-terminal peptide from PLD3 showed no change in half-life (fig. S2F). This finding indicates that the T185 variant of TMEM106B^FC^ has a slower degradation rate, which underlies its increased accumulation in neurons relative to the S185 variant.

We next asked whether neurons expressing PLD3-TMEM106B^FC^ formed TMEM106B fibrils. Like many amyloid fibrils, TMEM106B fibrils are insoluble in sarkosyl-containing buffers (*26*–*29*). We assayed for TMEM106B^FC^ fibrils in sarkosyl-insoluble lysates from our PLD3-TMEM106B^FC^ iNeuron model via immunoelectron microscopy (immuno-EM), which revealed fibrils decorated with gold particles conjugated to an anti-TMEM106B^FC^ antibody (Fig. 2G). No such fibrils were present in iNeurons expressing the N-terminus of PLD3 alone (PLD3-control).

Structural insights into TMEM106B fibrils have relied on biochemical extraction of sarkosyl-insoluble material from human brain tissue (*26*–*29*), but this approach obscures the native subcellular location of fibrils. To directly resolve the localization of these fibrils at molecular resolution, we performed cryogenic-correlative-light and electron microscopy (cryo-CLEM) combined with cryo-electron tomography (cryo-ET) on iNeurons expressing PLD3-TMEM106B^FC^ (T185) cultured on electron microscopy grids. (Fig. 2H and fig. S3A). We observed fibrils directly within lysotracker-positive vesicles in iNeurons (Fig. 2I), indicating that these fibrils reside within the lysosomal lumen.

To directly visualize lysosomal contents at scale, we isolated lysosomes from iNeurons expressing PLD3-TMEM106B^FC^ or PLD3-control by Nycodenz density gradient centrifugation, confirmed lysosomal identity and integrity by immunoblotting (fig. S3B), and analyzed the lysosomal fractions by cryo-ET (Fig. 2J) (*47*). Filaments were present in ∼20% of lysosomes (*N*= 10 out of 50) from PLD3-TMEM106B^FC^-expressing neurons and in none of the lysosomes assessed (*N* = 55) from control neurons (Fig. 2K and fig. S3C and D). These filaments fell into two morphologically distinct classes: thin (<10 nm) and thick (>10 nm) (fig. S3D). To confirm that the filaments were composed of TMEM106B, we extracted sarkosyl-insoluble material from the lysosomal fractions and performed immuno-EM. The resulting fibrils were immunoreactive with an antibody against TMEM106B^FC^ (Fig. 2L). Together, these findings establish that TMEM106B^FC^ fibrils accumulate within the lysosomal lumen in neurons, consistent with the intra-lysosomal localization of the cleaved TMEM106B^FC^ domain prior to fibril assembly.

### Mice with TMEM106B fibrils develop neurodegeneration-associated phenotypes

Having established that PLD3-TMEM106B^FC^ generates intra-lysosomal fibrils in cultured iNeurons, we asked whether PLD3-TMEM106B^FC^ could drive fibrillization – and neurodegeneration – *in vivo*. We delivered adeno-associated virus (AAV) encoding PLD3-TMEM106B^FC^ (T185) or PLD3-control to the mouse brain by intracerebroventricular injection at postnatal day 0 and aged the animals for 13 months before analysis (Fig. 3A). Anti-TMEM106B^FC^ immunostaining revealed widespread transduction across cortex, hippocampus, pons, fiber tracts, and cerebellum in PLD3-TMEM106B^FC^ mice, with no signal in controls (Fig. 3B and fig. S4A). The staining pattern, which includes punctate and elongated morphologies, closely matched that seen in postmortem FTLD-TDP brain (Fig. 3B). Western blotting confirmed the production of cleaved TMEM106B^FC^ fragments and the presence of sarkosyl-insoluble TMEM106B^FC^ (Fig. 3C), recapitulating the solubility profile of TMEM106B^FC^ aggregates in human disease.

**Figure 3.**
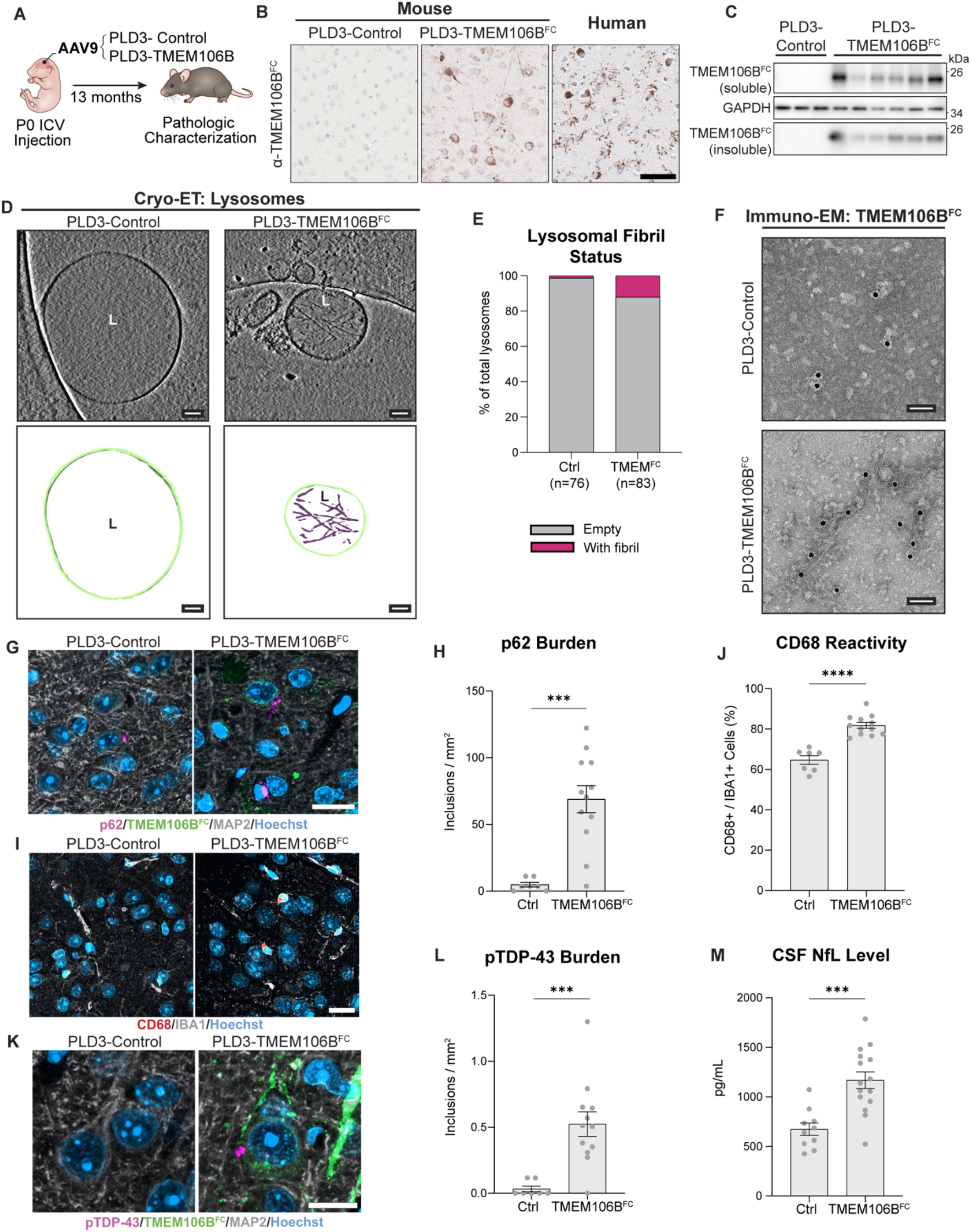
Intra-lysosomal TMEM106B fibrils from PLD3-TMEM106B^FC^ drive neuropathologic phenotypes in mice. **(A)** Schematic of an *in vivo* experiment to study the effects of TMEM106B fibril accumulation in mice. Mice were injected with AAV9 encoding PLD3-alone control or PLD3-TMEM106B^FC^ (T185) and aged for 13 months. **(B)** Representative immunohistochemical labeling of the TMEM106B fibril core in control or PLD3-TMEM106B^FC^ mouse cortex, and in human FTLD-TDP frontal cortex. Scale bar: 70 µm. **(C)** Western blots from mouse hemibrain expressing PLD3-TMEM106B^FC^ or PLD3-alone control. **(D)** Top: Representative slices through the reconstructed deconvolved tomograms of lysosomes (‘L’) isolated from PLD3-control (left) or PLD3-TMEM106B^FC^ mice (right). Below: Segmentation of corresponding tomograms on top with lysosome and intra-lysosomal fibrils pseudo colored in light green and purple, respectively. Scale bar: 50 nm. **(E)** Quantification of lysosomes identified from cryo-tomograms of gradient fractions, categorized as empty (gray) or filament-containing (pink) in each experimental group. **(F)** Representative negative stain electron microscopy images of anti-TMEM106B^FC^ immunogold-labeled, sarkosyl-insoluble material extracted from lysosomes isolated from PLD3-control (top) and PLD3-TMEM106B^FC^ (bottom) mice. Scale bar: 50 nm. **(G)** Representative co-immunofluorescent staining of p62, TMEM106B^FC^, MAP2 and nuclei (Hoechst) in cortex of WT mice expressing PLD3-control or PLD3-TMEM106B^FC^ (T185). Scale bar: 20 µm. **(H)** Quantification of cortical p62 burden in mice expressing PLD3-control or PLD3-TMEM106B^FC^ (T185). Mann-Whitney test. WT N= 7 PLD3-control, 12 PLD3-TMEM106B^FC^. **(I)** Representative co-immunofluorescent staining of CD68, IBA1, and nuclei (Hoechst) in cortex of mice expressing PLD3-control or PLD3-TMEM106B^FC^ (T185). Scale bar: 15 µm. **(J)** Quantification of cortical CD68 burden in WT and GRN-/-mice expressing PLD3-control or PLD3-TMEM106B^FC^ (T185). Student’s t-test. WT N= 7 PLD3-control, 12 PLD3-TMEM106B^FC^. **(K)** Representative co-immunofluorescent staining of pTDP-43 (Ser409/410), TMEM106B^FC^, MAP2 and nuclei (Hoechst) in cortex of mice expressing with PLD3-control or PLD3-TMEM106B^FC^ (T185). Scale bar: 10 µm. **(L)** Quantification of pTDP-43 inclusion density in mice expressing PLD3-control or PLD3-TMEM106B^FC^. Mann-Whitney test. WT N= 7 PLD3-control, 12 PLD3-TMEM106B^FC^. **(M)** Quantification of CSF NfL in mice expressing PLD3-control or PLD3-TMEM106B^FC^ (T185). Student’s t-test. WT N= 7 PLD3-control, 12 PLD3-TMEM106B^FC^. \**P* ≤ 0.05, \*\**P* ≤ 0.01, \*\*\**P* ≤ 0.001, \*\*\*\**P* ≤ 0.0001, ns: nonsignificant. All statistically analyzed data graphed as mean + SEM.

We next characterized the cellular and subcellular distribution of TMEM106B^FC^ aggregates in the mouse brain by co-staining tissue sections with cell-type markers and lysosomal proteins. TMEM106B^FC^ was expressed predominantly in neurons, with occasional astrocytic signal and little microglial labeling (fig. S4B). This distribution mirrors that of TMEM106B^FC^ in postmortem human brain, where neuronal and astrocytic staining predominate and microglial signal is sparse (fig. S4C), consistent with prior studies in human cohorts (*31, 48, 49*). TMEM106B^FC^ co-localized with cathepsin D in the mouse brain (fig. S4B), as well as postmortem human brain (fig. S4C), confirming intra-lysosomal localization of the fibril core. Overexpression of PLD3-TMEM106B^FC^ in mice therefore reproduces the cell type specificity, subcellular localization, and biochemical insolubility of TMEM106B^FC^ pathology observed in postmortem human brain.

To determine whether intra-lysosomal TMEM106B filaments form *in vivo*, we isolated lysosomes from PLD3-TMEM106B^FC^ and PLD3-control mouse brains by density gradient centrifugation (fig. S4D) and analyzed the fractions by cryo-ET and immuno-EM. Fibrils were present in 12% of lysosomes analyzed from PLD3-TMEM106B^FC^ mice, compared with 1% in controls (Fig. 3D, E, and fig. S4E).

Immuno-EM confirmed that these fibrils were immunoreactive for TMEM106B^FC^ in PLD3-TMEM106B^FC^ mice but not in controls (Fig. 3F and fig. S4F). TMEM106B^FC^ overexpression in the mouse brain, therefore, drives intra-lysosomal fibril formation *in vivo*, establishing this AAV model as a platform for interrogating the pathological consequences of TMEM106B^FC^ fibrils. Moreover, TMEM106B^FC^ overexpression in mice led to elevated p62 deposition in the cortex compared with controls (Fig. 3G and H). As p62 accumulates at damaged lysosomes and marks impaired autophagic clearance (*50*), this observation suggests that intra-lysosomal fibrils deleteriously alter lysosomal biology.

Neuroinflammation, characterized by microglial and astrocyte activation, accompanies neuronal dysfunction across neurodegenerative diseases (*51*). CD68 and IBA1-positive cells were elevated in mice overexpressing TMEM106B^FC^ relative to controls, indicating microglial activation (Fig. 3I, J, and fig. S4G). No changes were seen in GFAP astrocyte immunoreactivity (fig. S4H). Because AAV9 does not transduce microglia, this microglial response likely reflects a non-cell-autonomous reaction to pathology in neighboring neurons or astrocytes.

Phosphorylated TDP-43 (pTDP-43) inclusions are the defining neuropathological feature of FTLD-TDP, amyotrophic lateral sclerosis (ALS), limbic-predominant age-related TDP-43 encephalopathy (LATE) and occur in over half of AD cases (*52, 53*). TMEM106B fibril burden correlates with pTDP-43 load in FTLD-TDP (*31*), but whether TMEM106B^FC^ fibrillization drives pTDP-43 pathology has remained untested. We observed cortical pTDP-43 inclusions in 11 of 12 PLD3-TMEM106B^FC^ mice, whereas such inclusions were rare in controls (Fig. 3K, L and fig. S4I); immuno-EM confirmed pTDP-43 aggregates in sarkosyl-insoluble lysosomal fractions from PLD3-TMEM106B^FC^ mice (fig. S4J). These findings indicate that TMEM106B^FC^ accumulation promotes TDP-43 aggregation, a finding consistent with the strong association between *TMEM106B* variants and TDP-43 proteinopathies (*15, 54*).

Elevated neurofilament light chain (NfL) in the cerebrospinal fluid (CSF) is an established biomarker of neurodegeneration across neurological diseases, including AD and FTLD-TDP (*55, 56*). We found elevated CSF NfL levels in PLD3-TMEM106B^FC^ mice relative to controls (Fig. 3M), indicating that the TMEM106B^FC^ fibrillization and other neuropathological changes observed in these animals are accompanied by active neurodegeneration. Together, these results establish that overexpression of PLD3-TMEM106B^FC^ in mice recapitulates the formation of intra-lysosomal TMEM106B fibrils *in vivo*. Moreover, fibrillization was sufficient to drive cardinal neuropathologic features of FTLD-TDP and neurodegeneration.

### Progranulin levels regulate TMEM106B^FC^ accumulation

Beyond *TMEM106B* coding variants, are there additional factors that regulate the abundance of TMEM106B fibrils? In familial FTLD, *GRN* and *TMEM106B* genetically interact to modify disease penetrance and age-of-onset (*23, 24, 57*–*59*). *GRN* encodes progranulin, an intra-lysosomal protein, and rare pathogenic *GRN* coding variants reduce progranulin express by half (*60*). Common non-coding variants at the *GRN* locus modestly reduce progranulin expression and are associated with cognitive decline in the elderly and an increased risk of several age-related neurodegenerative diseases (*61*–*63*). The biological mechanism underlying the *TMEM106B* and *GRN* genetic interaction is unknown. We hypothesized that *GRN* and *TMEM106B* converge on the regulation of TMEM106B^FC^ levels (Fig. 4A).

**Figure 4:**
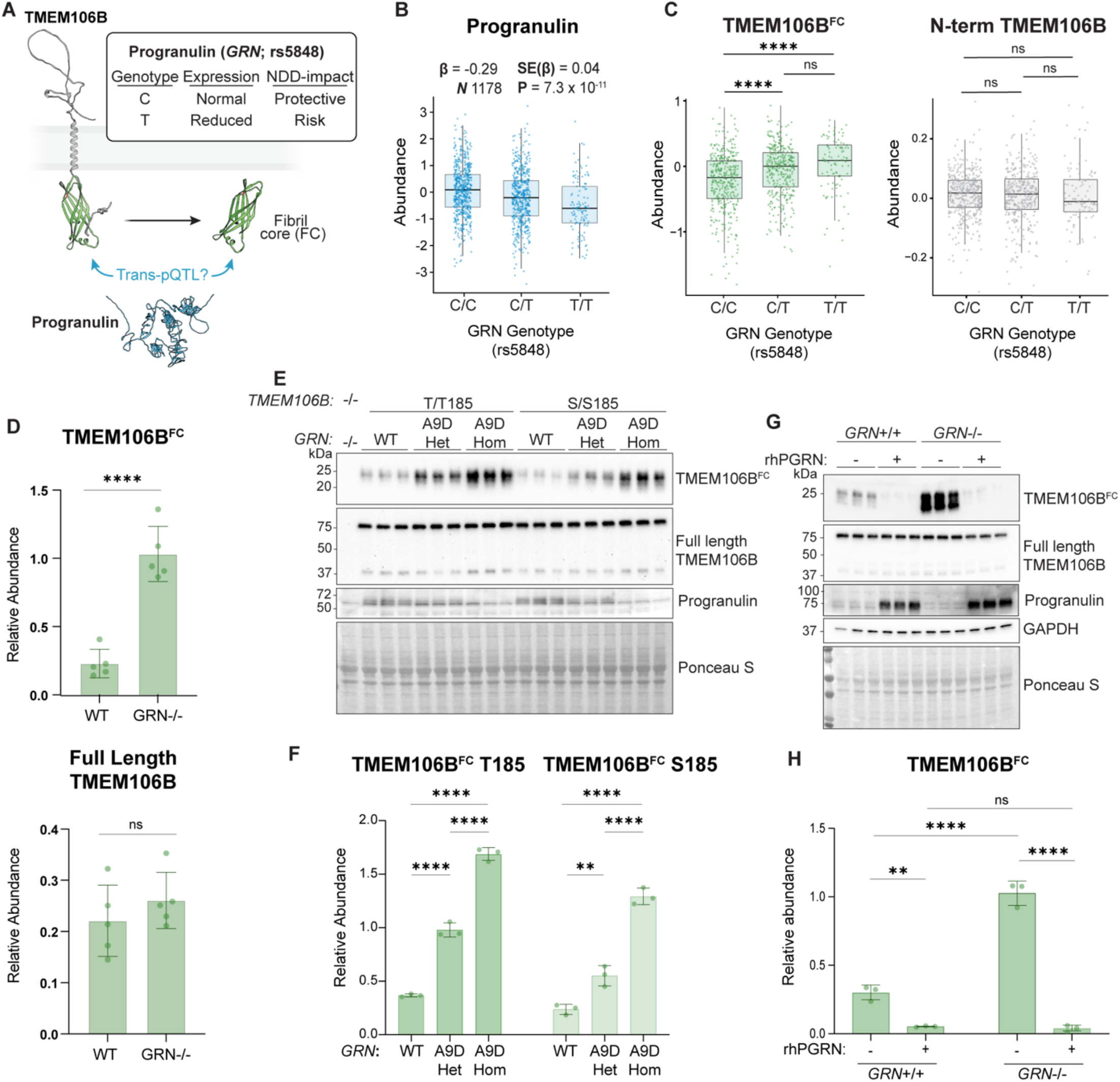
Genetic Variants in *TMEM106B* and *GRN* influence TMEM106B^FC^ levels. **(A)** Schematic illustrating lysosomal localization of TMEM106B and progranulin. Genetic interactions between *TMEM106B* and *GRN* variants drive neurodegeneration when co-inherited. An AD-risk variant in *GRN* is known to reduce progranulin levels and is associated with AD. We hypothesize that progranulin loss alters TMEM106B^FC^ accumulation. **(B)** *Cis* pQTL analyses of *GRN* rs5848 genotype versus PGRN levels in ROSMAP cohort. One-way ANOVA. N and SE indicated above graph. **(C)** *Trans* pQTL analyses of *GRN* rs5848 genotype versus TMEM106B N-terminal (right) and C-terminal (left) domain levels in ROSMAP cohort. Statistics by one-way ANOVA with Tukey post-hoc. N = 333 C/C, 296 C/G, and 90 T/T individuals. **(D)** Top: Quantification of TMEM106B^FC^ from Western blot in fig. S6A normalized to Ponceau S as loading control. Student’s t-test. N = 5, error bars = SD. Bottom: Quantification of full-length TMEM106B (monomer+dimer sum) from Western blot in fig. S6A normalized to Ponceau S as loading control. Student’s t-test. N = 5, error bars = SD. **(E)** Western blots of TMEM106B^FC^ and full-length TMEM106B in T185- and S185-expressing iNeurons engineered to express WT *GRN* or the *GRN*-A9D mutation (heterozygous, Het; and homozygous, Hom). N= 3. Error bars = SD. **(F)** Quantification of TMEM106B^FC^ level in panel E, normalized to Ponceau S as a loading control. One-way ANOVA within T185 or S185 groups, followed by Tukey’s multiple comparisons test. N= 3. Error bars = SD. **(G)** Western blots of endogenous TMEM106B^FC^ and full-length TMEM106B in *GRN* WT or *GRN* KO iNeurons with or without added rhPGRN in the media. N= 3. Error bars = SD. **(H)** Quantification of TMEM106B^FC^ in panel G, normalized to Ponceau S as a loading control. Two-way ANOVA with Šídák’s multiple comparisons test. N= 3. Error bars = SD. \**P* ≤ 0.05, \*\**P* ≤ 0.01, \*\*\**P* ≤ 0.001, \*\*\*\**P* ≤ 0.0001, ns: nonsignificant.

Recently, a pQTL analysis of 1,369 postmortem human brain samples identified rs5848, a noncoding variant in the *GRN* gene, as both a *cis-*pQTL for progranulin and a *trans-*pQTL for TMEM106B (Fig. 4B, fig. S5A, and Table S1) (*35*). Specifically, the risk-associated rs5848 T allele was associated with reduced progranulin, as is consistent with prior observations (*64*–*66*), and elevated TMEM106B (Fig. 4B and fig. S5A). In ROSMAP brain tissue, rs5848 associated with the C-terminal fibril core domain but not the N-terminal domain, both at the individual peptide and domain level (Fig. 4C, fig. S5B, and Tables S3 and S4). Relative to CC carriers, rs5848 CT and TT carriers had higher levels of the fibril core domain (Fig. 4C). The risk-associated rs5848 T allele thus associates with both lower progranulin and higher TMEM106B^FC^ levels, suggesting that progranulin and TMEM106B^FC^ levels are inversely, and mechanistically, related.

We turned to cellular models to directly test whether progranulin loss modulates TMEM106B^FC^ accumulation. We examined TMEM106B^FC^ levels in iNeurons derived from isogenic wild-type and *GRN* knockout (KO) iPSCs (fig. S6A). Consistent with our pQTL analysis, progranulin loss drove an increase in TMEM106B^FC^ abundance without altering full length TMEM106B levels or its dimer-to-monomer ratio (Fig. 4D and fig. S6A).

To determine whether disease-causing *GRN* mutations recapitulate the effect of progranulin loss on TMEM106B^FC^ levels in neurons, and to establish whether *GRN* variants interact with the *TMEM106B* coding variant, we engineered iPSCs to carry wild-type, heterozygous, or homozygous FTLD-allele dosage in both *TMEM106B* genetic backgrounds (Fig. 4E and F). The combination of homozygous *GRN*-A9D and the *TMEM106B* T185 risk variant produced the highest TMEM106B^FC^ levels, while full-length TMEM106B was unaffected across genotypes (Fig. 4E, F, and fig. S6D). Knocking out *GRN* in an independent isogenic T185/S185 iPSC line confirmed that the combination of progranulin loss and the *TMEM106B*-T185 variant causes the highest abundance of TMEM106B^FC^ (fig. S6E and F). Together, these data place *GRN* and *TMEM106B* in a unified post-translational axis that additively promotes neuronal TMEM106B^FC^ accumulation.

Cells can internalize progranulin from the extracellular space, whereupon it traffics to lysosomes, and exogenous progranulin rescues pathological phenotypes of *GRN*-KO cells and mice (*67, 68*). Modified recombinant progranulin with enhanced brain delivery is currently under evaluation in clinical trials for FTLD. We therefore asked whether exogenous progranulin is sufficient to reverse TMEM106B^FC^ accumulation. Recombinant human progranulin (rhPGRN) dramatically reduced TMEM106B^FC^ levels in *GRN*-KO iNeurons, and in iNeurons overexpressing PLD3-TMEM106B^FC^, without affecting full-length TMEM106B expression (Fig. 4G, H, and fig. S6G and H). Notably, rhPGRN almost completely abolished TMEM106B^FC^ in wild-type iNeurons, suggesting supra-physiological effects on TMEM106B^FC^ degradation rates (Fig. 4G and H). Thus, progranulin acts upstream of TMEM106B^FC^ abundance: progranulin loss elevates the fibril-forming fragment, while exogenous progranulin nearly eliminates it.

### TMEM106B fibrils accumulate within lysosomes in FTLD-TDP brain

To determine whether progranulin loss drives insoluble TMEM106B fibril accumulation in human disease, we stratified previously analyzed (*31*) postmortem FTLD-TDP cortex samples by *GRN* status and quantified full-length TMEM106B and sarkosyl-insoluble TMEM106B^FC^ by immunoblot analysis (fig. S7A). Twenty three percent of cases carried pathogenic *GRN* mutations, termed FTLD-*GRN*. Full-length TMEM106B expression and monomer-to-dimer ratio were equivalent between sporadic FTLD (sFTLD) and FTLD-*GRN* cases (Fig. 5A and fig. S7A and B). FTLD-*GRN* cases had substantially higher insoluble TMEM106B^FC^ than sFTLD cases (Fig. 5A), and the highest levels occurred in *GRN* mutation carriers with homozygous *TMEM106B* C/C (T/T185) risk genotypes (fig. S7C), mirroring the additive genetic effect seen in iNeurons (Fig. 4E). Progranulin levels and *TMEM106B* risk variant dosage therefore jointly determine insoluble TMEM106B fibril burden in human FTLD brain.

**Figure 5.**
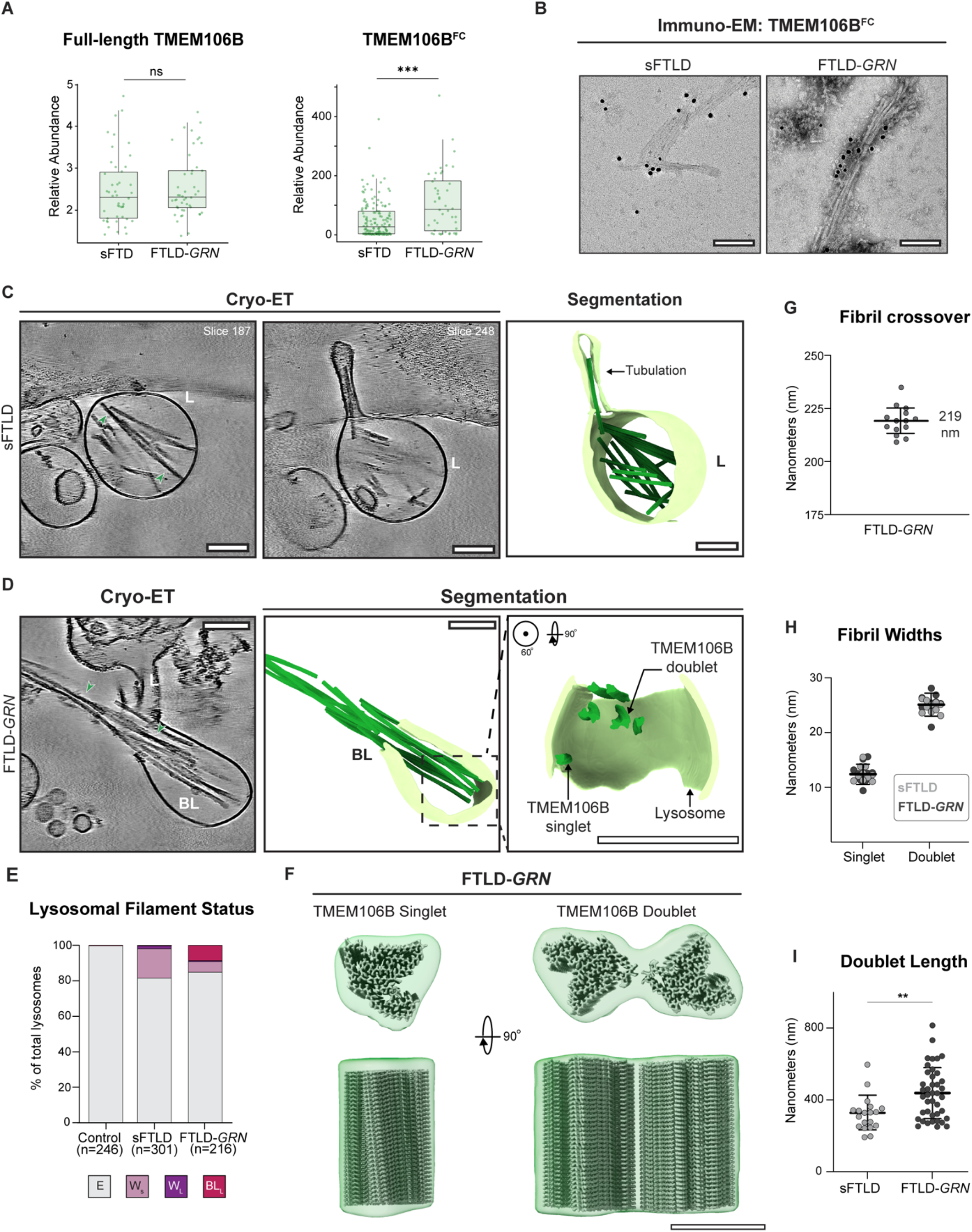
Patient brain-derived lysosomes contain TMEM106B fibrils. **(A)** (left) Analysis of full-length TMEM106B from sFTD and FTLD-*GRN* RIPA-soluble frontal cortex, normalized to GAPDH (representative WB in fig. S7a). N=46 cases per group. (right) Analysis of insoluble TMEM106B^FC^ in sFTD vs. FTLD-*GRN* using immunoblotting data from Marks et al. (2024). N= 149 sFTD, 44 FTLD-*GRN*. Two-tailed Mann-Whitney U Test. Error Bars = SEM. **(B)** Representative negative stain electron microscopy images of anti-TMEM106B^FC^ immunogold-labeled sarkosyl-insoluble material extracted from lysosomes isolated from sFTLD (left) and FTLD-neuronal TMEM106B^FC^ abundance (*manuscript in preparation*). The molecular basis by which T185 slows TMEM106B^FC^ degradation remains unknown; candidates include structural changes that alter fibrillization potential, *GRN* (right) patient brain tissue. Scale bar: 50 nm. **(C)** Left and Middle: slices through reconstructed and denoised cryo-tomograms of fraction F4 from sFTLD-case, revealing TMEM106B filaments of varying morphology within single-membraned lysosomes (L). Arrowheads depict the crossover of TMEM106B doublets. Right: Corresponding segmentation and rendering of the tomograms on the top showing the lysosomal membrane (light green) and TMEM106B fibrils (dark green). Scale bar: 100 nm. **(D)** Left: slice through reconstructed and denoised cryo-tomograms of fraction F4 from an FTLD*-GRN* case, revealing TMEM106B filaments protruding from a broken lysosome (BL). Green arrowheads depict the crossover of TMEM106B doublets. Middle and Right: Corresponding segmentation and rendering of the tomograms on the left showing the lysosomal membrane (light green) and TMEM106B fibrils (dark green). Scale bar: 100 nm. **(E)** Quantification from cryo-ET data of patient brain-derived lysosomes based on the presence and the width of the fibrils ensconced: empty lysosomes (gray), lysosomes with filaments of width <10 nm (light pink), lysosomes with filaments of width >10 nm (purple), and broken lysosomes with filaments of width >10 nm (red). **(F)** Maps (green) of the TMEM106B singlet and doublet obtained from subtomogram averaging. Previously determined structures of singlet (PDB: 7SAQ), and doublet (PDB: 7SAR) represented as density maps (gray) fit as rigid bodies in the lysosomal TMEM106B singlet and doublet density maps. Scale bar: 12 nm. **(G)** Quantification of crossover distances of TMEM106B doublets within FTLD-*GRN* patient brain-derived lysosomes. The mean crossover distance matches the previously reported value of 210 nm. N = 10, std. deviation = 5.49 nm. **(H)** Quantification of the width of the TMEM106B singlet and doublet filaments in the patient brain-derived lysosome fractions. N = 10, std. deviation = 1.82 nm (Singlet) and 1.91 nm (Doublet). **(I)** Quantification of the length of the TMEM106B doublet filaments in the sFTLD and FTLD-*GRN* patient brain-derived lysosome fractions. N = 19 (sFTLD) and 41 (FTLD-*GRN*), std. deviation = 9

Our cell and mouse model data established that TMEM106B fibrils reside within lysosomes. We asked whether the same is true in human disease. Using an optimized density-gradient centrifugation method, we enriched lysosomes from the frontal cortex of one control, one sporadic FTLD-TDP, and one FTLD-*GRN* patient (fig. S8A and Table S10). Immuno-EM of sarkosyl-insoluble material from these fractions identified TMEM106B-immunopositive fibrils in both FTLD cases (Fig. 5B). Distinct fibril populations from these fractions were immunopositive for anti-pTDP-43 antibodies (fig. S8B and C), consistent with lysosome fractions from mice overexpressing PLD3-TMEM106B^FC^ (fig. S4J) and our recent immuno-EM and cryo-ET observations from other FTLD-TDP cases (*47*). Therefore, both TMEM106B and TDP-43 fibrils co-sediment with lysosomal fractions from human brain, but whether they reside within the lysosomal lumen or associate with its exterior cannot be determined from these experiments.

Cryo-ET of lysosomal fractions confirmed that TMEM106B fibrils were present within lysosomes of both FTLD cases but extremely rare in the control brain (Fig. 5C, E, and fig. S8D). In sFTLD, fibrils were exclusively contained within the lumen and frequently accompanied by lysosomal enlargement (fig. S9A). In several instances, fibrils extended into elongated tubular protrusions of the lysosomal membrane, suggesting that fibril growth from within the lysosome may physically deform the membrane (Fig. 5C, fig. S9B, and movie S1). Similar elongated fibrils were present in FTLD-*GRN* lysosomes (Fig. 5D). Fibril-containing lysosomes were rare in the control case (1 of 246 lysosomes, Fig. 5E), but immuno-EM, which detects concentrated sarkosyl-insoluble fibril material, confirmed that TMEM106B fibrils were present at low levels in this specimen (fig. S8B). This observation is consistent with their reported occurrence in the brains of cognitively normal elderly individuals (*28, 29*).

To confirm that the intra-lysosomal fibrils observed in cryo-ET were indeed TMEM106B, we computationally extracted the fibrils from tomograms and performed subtomogram averaging of these fibrils from lysosomes *in situ* (Fig. 5F and fig. S10A-D). Our analysis revealed an exact fit of previously determined TMEM106B singlet and doublet structures in the obtained density maps (Fig. 5F). We further characterized the doublet TMEM106B polymorph, which is unique among known amyloid fibrils in its crossover distance, filament diameter, and absence of a fuzzy surface coat (Table S5). The crossover distance averaged 219 ± 10 nm, closely matching the ∼210 nm reported from cryo-EM structures of TMEM106B fibrils (Fig. 5G and fig. S10E). Singlet and doublet diameters were ∼12 nm and ∼25 nm, respectively, in agreement with reported values of ∼12.5 nm and ∼26 nm (Fig. 5G) *(26)*. Finally, intra-lysosomal fibrils lacked an identifiable fuzzy surface coat. These characteristics confirm that the intra-lysosomal fibrils are composed of TMEM106B.

Although both cases contained intra-lysosomal fibrils, those in FTLD-*GRN* were substantially longer than in sFTLD (median 436.0 ± 142.7 nm vs. 321.2 ± 96.9 nm; Fig. 5I), consistent with the impaired TMEM106B fibril core degradation observed in *GRN*-A9D neurons (Fig. 4E). Exceptionally long fibrils extended from within lysosomes through the membrane and into the surrounding space (Fig. 5D fig. S8E, and movie S2). Lysosomal membrane rupture accompanied all such extrusion events. The ends of some protruding fibrils pressed against the inner leaflet of the lysosomal membrane, suggesting the fibrils generate sufficient force to breach the membrane as they elongate (fig. S9B). Although the proportion of lysosomes containing fibrils was similar between sFTLD and FTLD-*GRN*, membrane rupture was mainly observed in FTLD-*GRN* (Fig. 5E). Together, these data establish that TMEM106B fibrils accumulate within lysosomes in human FTLD brain, consistent with the lysosome as their potential site of origin, and that fibril elongation may rupture the lysosomal membrane, a phenomenon more common in the context of the exaggerated fibril burden of FTLD-*GRN*.

## Discussion

Genetic studies have linked *TMEM106B* and *GRN* to age-related cognitive decline and multiple neurodegenerative diseases, but the underlying mechanisms have remained elusive. Both proteins localize to lysosomes, implicating lysosomal dysfunction as a shared disease mechanism (*45, 69, 70*). Supporting this, *GRN* and *TMEM106B* genotypes associate with transcriptomic signatures of lysosomal dysfunction in the human brain (*25*). Here, we provide a mechanistic basis for these genetic observations: *GRN* and *TMEM106B* risk variants drive accumulation of TMEM106B^FC^ within lysosomes, leading to the accumulation of neurotoxic TMEM106B amyloid fibrils within lysosomes.

Prior studies have linked age and the *TMEM106B* risk haplotype to fibril burden (*31, 32, 71*), but the causal variant within this haplotype remains unknown. The haplotype contains several noncoding *TMEM106B* variants, some of which have been suggested to contribute to neurodegeneration (*33, 41, 42, 72, 73*), as well as the coding variant T185 examined here. In human studies, contributions from these other linked variants cannot be excluded. However, genetic engineering of the coding variant alone is sufficient to alter TMEM106B^FC^ abundance in iNeurons without affecting full-length protein levels, identifying a post-translational mechanism that operates independently of the noncoding variants.

The T185 variant prolongs the half-life of TMEM106B^FC^ relative to the protective S185 variant, thereby raising intra-lysosomal TMEM106B^FC^ levels. This effect is compounded byprogranulin loss, providing a clear mechanistic explanation of the genetic link between *TMEM106B* and *GRN* variants in disease. In a parallel study, the Gitler and Abu-Remaileh labs also observed that *TMEM106B* and *GRN* genotypes affect differences in glycosylation at a neighboring asparagine, altered trimming of the C-terminal tail, or differences in TMEM106B^FC^ protein-protein interactions. While the precise biological function of progranulin remains unclear, the mechanism by which progranulin loss elevates fibril burden is more straightforward to hypothesize: Progranulin deficiency impairs lysosomal protease activity and raises lysosomal pH (*74*), thereby indirectly reducing TMEM106B^FC^ degradation.

Both common and rare genetic variants reduce progranulin levels in the brain, causing neurodegeneration and cognitive decline. Restoring progranulin is therefore a promising therapeutic strategy for these conditions, and clinical trials evaluating progranulin restoration in FTD-*GRN* are ongoing (*75*–*81*). We found that neurons exposed to supra-physiologic levels of recombinant progranulin had nearly undetectable levels of TMEM106B^FC^, even when TMEM106B^FC^ was overexpressed (Fig. 4G and fig. S6H). These observations predict that raising progranulin levels in patients will substantially reduce TMEM106B fibril burden through reducing the TMEM106B^FC^ fibril precursor. Because we show that *GRN* and *TMEM106B* variants are associated with elevated TMEM106B^FC^ levels in both AD and FTD, boosting lysosomal progranulin could have therapeutic relevance beyond FTLD-*GRN*.

Although both progranulin and TMEM106B reside in lysosomes, the intracellular location of TMEM106B fibrils was previously unknown. We show that TMEM106B fibrils accumulate within lysosomes (Fig. 5). While other amyloid fibrils have been observed inside lysosomes, including TDP-43, tau, and synuclein (*47, 82, 83*), their monomeric precursors are cytoplasmic, not lysosomal. For TMEM106B, however, both the native precursor protein and the fibrils exist within lysosomes. Thus, unlike other amyloid fibrils, TMEM106B fibrils have continuous access to substrate within the same closed compartment, potentially enabling self-sustaining fibril amplification. This scenario resembles recent findings with LLOMe, a lysosome-damaging agent that concentrates within lysosomes. Above a critical threshold, LLOMe forms intra-lysosomal amyloid fibrils that physically rupture the membrane (*84, 85*).

Based on our findings, we propose a model wherein healthy lysosomes degrade TMEM106B^FC^ at rates sufficient to limit fibril nucleation and extension. When lysosomal function declines during aging or disease, monomer production outpaces degradation, driving fibril growth. Genetically driven progranulin loss and the TMEM106B T185 coding variant act together in a two-hit process to promote toxic fibrillization: T185 slows TMEM106B^FC^ degradation kinetics, and progranulin loss compounds this by indirectly impairing lysosomal proteolysis.

In sFTLD, we observed a subset of lysosomes with deformed or tubular membranes surrounding elongated TMEM106B fibrils (fig. S9). Such lysosomes could represent a pre-rupture state, although lysosomes containing fibrils from the sFTLD patient tolerated large protrusions (∼1.5 µm) without rupturing (fig. S9). In contrast, we observed ruptured lysosomes with vase-like morphologies partially enveloping elongated fibrils in the FTLD-*GRN* case (Fig. 5D). Lysosomal rupture may be more common in FTLD-*GRN* for multiple reasons. First, TMEM106B fibrils in the FTLD-*GRN* case were longer than in the sFTLD case (Fig. 5I). This is consistent with our observations in cultured neurons, where progranulin loss increases TMEM106B^FC^ levels. Such increases in the levels of fibril precursors could potentiate fibril elongation, which could exert additional strain on the lysosome from multiple TMEM106B fibrils extending in parallel. Second, progranulin deficiency impairs lysosomal protein sorting in a neuron-specific manner, potentially directly compromising lysosomal membrane integrity. For example, *GRN* loss depletes TMEM63A (*86*), a lysosome-resident mechanosensitive cation channel that relieves hydrostatic pressure and membrane tension (*87*). Loss of TMEM63A would prevent lysosomes from accommodating the mechanical stress of fibril elongation, predisposing them to rupture.

The elevated p62 foci observed in mice expressing PLD3-TMEM106B^FC^ (Fig. 3H) may reflect the lysosomal damage exerted by TMEM106B fibril elongation. Severely damaged lysosomes recruit p62 and are targeted for lysophagy, as shown with LLOMe-induced amyloid filaments (*50*). Alternatively, ruptured lysosomes may release TMEM106B fibrils into the cytoplasm, where ubiquitination would recruit p62 for autophagosomal targeting (*88*).

Other amyloid fibrils, such as α-synuclein, spread through lysosomal escape (*89*). Lysosomal rupture could enable TMEM106B fibril propagation in a feed-forward cycle: Even minor lysosomal damage from intra-lysosomal fibrils could impair degradative capacity, resulting in fibril propagation and escape. Fibrils could spread to new lysosomes via autophagy or lysophagy, or they could propagate to neighboring cells through lysosomal exocytosis. Notably, we observed rare microglia staining positive for TMEM106B^FC^ in mice expressing PLD3-TMEM106B^FC^ (fig. S4B). Because our AAV does not transduce microglia, this TMEM106B^FC^ likely originated from neurons or astrocytes, suggesting potential intracellular fibril spread in the mouse brain.

Could TMEM106B fibril-induced lysosomal damage contribute to the pathological accumulation of other protein aggregates in neurodegenerative diseases? TDP-43, tau, and α-synuclein fibrils have all been detected within lysosomes (*47, 82, 83, 90*). Lysosomes damaged by TMEM106B fibrils may have impaired degradative capacity, predisposing cells to secondary aggregate accumulation. In cultured neurons, TDP-43 seeding potential correlates with TMEM106B fibril burden in brain extracts from FTLD-TDP patients (*91*). Mice expressing PLD3-TMEM106B^FC^ also accumulated pTDP-43 foci, and TDP-43 fibrils were present in lysosomal fractions. Therefore, TMEM106B fibril-induced lysosomal dysfunction likely potentiates TDP-43 co-pathology. If the same holds true for tau and α-synuclein, this mechanism could explain why *TMEM106B* variants increase risk across neurodegenerative diseases with distinct proteinopathies.

Our work demonstrates that the production of TMEM106B fibrils is sufficient to induce neuropathology and neurodegeneration *in vivo*. Mice with intra-lysosomal TMEM106B fibrils developed several molecular markers of neurodegeneration, including microgliosis, pTDP-43 accumulation, and elevated CSF NfL (Fig. 3). CD68-positive microglial activation could reflect direct sensing of fibrils released by neurons or astrocytes, or indirect activation arising from neuronal damage. Elevated CSF NfL levels suggest neuronal damage in these animals, and future analyses could reveal whether this damage leads to behavioral or cognitive deficits with age.

Variants in *TMEM106B* and *GRN* that affect TMEM106B^FC^ levels are common, even in healthy individuals. Amongst healthy controls in our ROSMAP dataset, 83% and 56% of people carried at least one risk allele of *TMEM106B* rs3173615 and *GRN* rs5848, respectively (Tables S2 and S4). Gradual intra-lysosomal TMEM106B fibril accumulation, shaped by progranulin levels and *TMEM106B* variants, may drive neuronal dysfunction and cognitive impairment in otherwise healthy people. Consistent with this, protective *TMEM106B* and *GRN* alleles are enriched in cognitively healthy centenarians, and the protective *TMEM106B* allele associates with cortical neuron abundance in cognitively healthy elderly individuals, suggesting that reduced fibril accumulation preserves cognitive function during aging (*11, 19*). Enhancing lysosomal degradative capacity to more efficiently clear TMEM106B^FC^ might therefore slow age-related lysosomal damage that insidiously contributes to cognitive decline.

In summary, we show that risk variants in *GRN* and *TMEM106B* convergently increase TMEM106B^FC^ levels through impaired degradation, and that the resulting accumulation of intra-lysosomal TMEM106B fibrils associates with lysosomal damage. This feed-forward mechanism likely operates wherever lysosomal function is compromised, across neurodegenerative diseases and aging, making TMEM106B^FC^ degradation a compelling therapeutic target for preserving cognitive health.

## Supporting information

D_1000306295_val-report-full_P1

D_1000306332_val-report-full_P1

## Acknowledgements

We would like to thank Frances Diehl, Joseph Replogle, and members of the Ward lab for helpful comments and discussions on this manuscript. We would like to thank Durga Atili for his help with iNeuron *in situ* cryo-ET experiments and Jorge Alaiz Noya for his help with mouse experiments. This work was supported, in part, through the Intramural Research Program of the National Institutes for Neurological Diseases at the NIH, the Kissick Family Foundation, the Rainwater Charitable Foundation, and the Chan Zuckerberg Initiative (MEW). JMR was supported by an Association for Frontotemporal Dementia (AFTD) Holloway Fellowship.

## Author Contributions

Conceptualization: JMR, SF, LP, MEW, Experiments: JMR, JDM, MGF, HY, RD, IK, AF, RS, AK, ISN, EW, MY, IPC, DY, US, RS, VMJ, YC, KJW, MCC, NP, SM, Data Analysis and Interpretation: JMR, JDM, MGF, AF, BK, EW, VMJ, DY, US, TWT, YZ, SM, MEW, AS, DD, AW, YL, Key Resources: JMR, YAQ, LH, MD, DWD, YZ, MP, BFB, GSD, KAJ, RCP, NG-R, MD, BR EE-C, MCC, NP, SJB, YZ, SM, LP, MEW, NTS, TSW, WCS, JAM, Writing-original draft: JMR, JDM, Writing-revisions and editing: SM, MEW, LP, TWT, all other authors

## Data, Code, and Materials Availability

All data described in this study are included in the manuscript or supplementary materials. The structures of the TMEM106B fibrils obtained by subtomogram averaging have been deposited in the Electron Microscopy Data Bank (EMDB) under accession codes: EMD-76248 (TMEM106B Singlet) and EMD-76230 (TMEM106B Doublet). Materials, reagents, and resources generated as part of this study are available from the corresponding authors upon reasonable request and will be shared with qualified researchers according to guidelines set for by the institutions involved. Postmortem human specimens were obtained from the Mayo Clinic Brain Bank.

## Supplementary Materials

**Figure S1.**
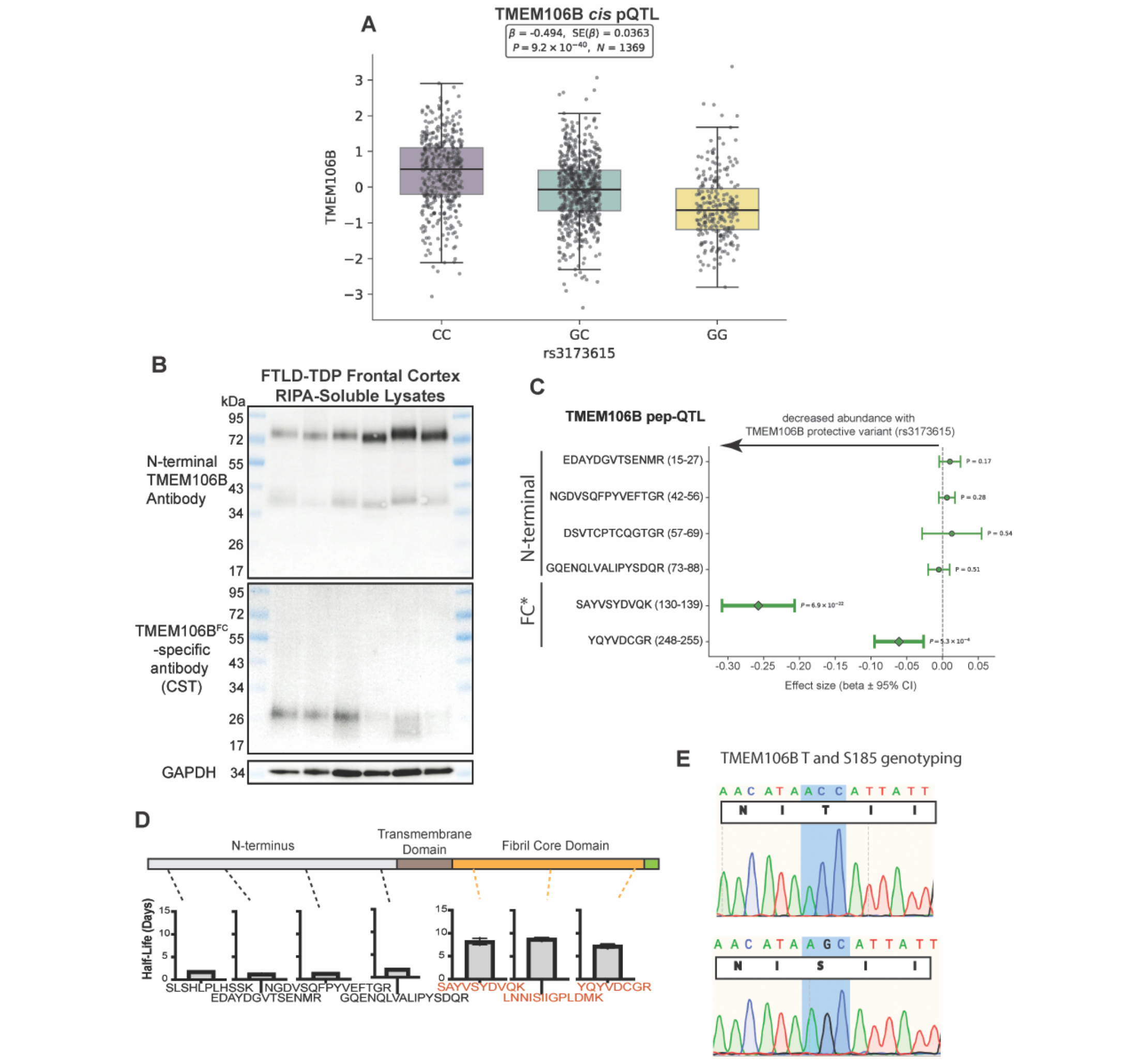
The C-terminal fragment from TMEM106B is increased in abundance with the risk allele. **(A)** *Cis* pQTL analyses of *TMEM106B* rs3173615 genotype versus TMEM106B protein levels in the ROSMAP cohort. N and SE indicated above graph. **(B)** Western blot of RIPA-soluble lysates from FTLD-TDP frontal cortex using an N-terminal TMEM106B antibody (top) or anti-TMEM106B fibril core antibody (CST #87145). GAPDH (bottom) as loading control. **(C)** *Cis*-peptide-QTL analyses for each individual TMEM106B peptide well-represented in the ROSMAP data. For each peptide, association was measured between peptide abundance values and *TMEM106B* rs3173615 genotype. P-value indicated next to each bar; Effect size indicated on the x-axis. Negative effect size indicates decreased abundance with the minor allele (G for rs3173615). FC: Fibril Core *YQYVDCGR peptide was included as a ‘fibril core’ peptide due to evidence of C-terminal boundary from Held et al. **(D)** dSILAC measured half-lives of TMEM106B peptides from iNeurons overexpressing TMEM106B (S185). N=3. Position of Peptide in Protein indicated by dotted lines to protein schematic. Nested T-Test on N-terminal vs C-terminal peptides produces p < 0.0001 (not shown on graphs). Error bars = SD. **(E)** TMEM106B sanger sequencing data from isogenic KOLF2.1J cell lines edited to express either a homozygous TMEM106B TT185 (C/C genotype) or TMEM106B SS185 (G/G genotype) variant. \**P* ≤ 0.05, \*\**P* ≤ 0.01, \*\*\**P* ≤ 0.001, \*\*\*\**P* ≤ 0.0001, ns: nonsignificant.

**Figure S2.**
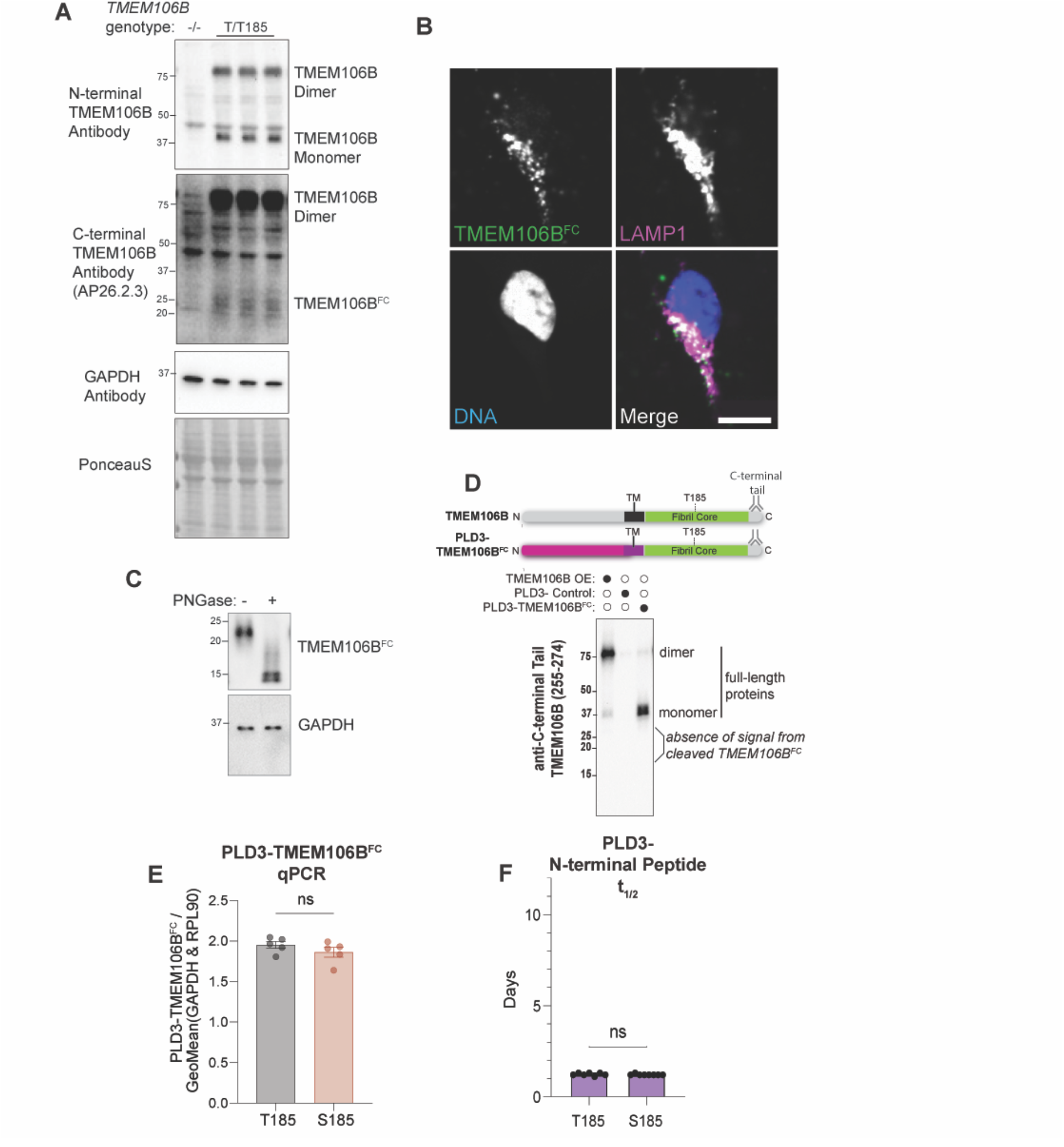
The PLD3-TMEM106B chimera mimics the biology of TMEM106B^FC^ from the endogenous TMEM106B protein. **(A)** Western blot of lysates from isogenic iNeurons with either TMEM106B-/-(left lane) or expressing the TMEM106B T/T185 using a C-terminal TMEM106B antibody (AP26.2.3) or N-terminal TMEM106B antibody (CST #93334). GAPDH and Ponceau S shown as loading controls. TMEM106B full-length dimer and cleaved TMEM106B C-terminal fragment indicated on right. N= 1 for TMEM106B-/-, 3 for T/T185. **(B)** Representative immunofluorescence image of an iNeuron expressing the PLD3-TMEM106B^FC^ chimera, stained for TMEM106B^FC^ (green) and co-stained with LAMP1 (magenta) and Hoechst33342 for DNA (blue). Scale bar: 10 µm. **(C)** Western blot of lysates from iNeurons expressing PLD3-TMEM106B^FC^ (T185) treated with either vehicle or PNGase to remove glycans. Stained with antibodies for the TMEM106B Fibril Core (AP26.2.3), OR GAPDH. **(D)** Western blot of lysates from iNeurons expressing TMEM106B (T185) (left lane), PLD3-control (middle lane), or PLD3-TMEM106B (right lane) stained with an antibody targeting the C-terminal tail (beyond residue 255) of TMEM106B. Re-probed blot from Figure 2B, which depicts loading control for this same blot. **(E)** qPCR of iNeurons expressing PLD3-TMEM106B^FC^ (T185) or PLD3-TMEM106B^FC^ (S185). N = 5. Student’s t-test. Error bars = SD. **(F)** dSILAC-derived half-life (t_1/2_) of N-terminal PLD3-peptide, LMYQELK, from iNeurons expressing PLD3-TMEM106B^FC^ T185 or S185 variant. Student’s t-test. N=7 (T185) or 8 (S185). Error bars = SD. \**P* ≤ 0.05, \*\**P* ≤ 0.01, \*\*\**P* ≤ 0.001, \*\*\*\**P* ≤ 0.0001, ns: nonsignificant.

**Figure S3:**
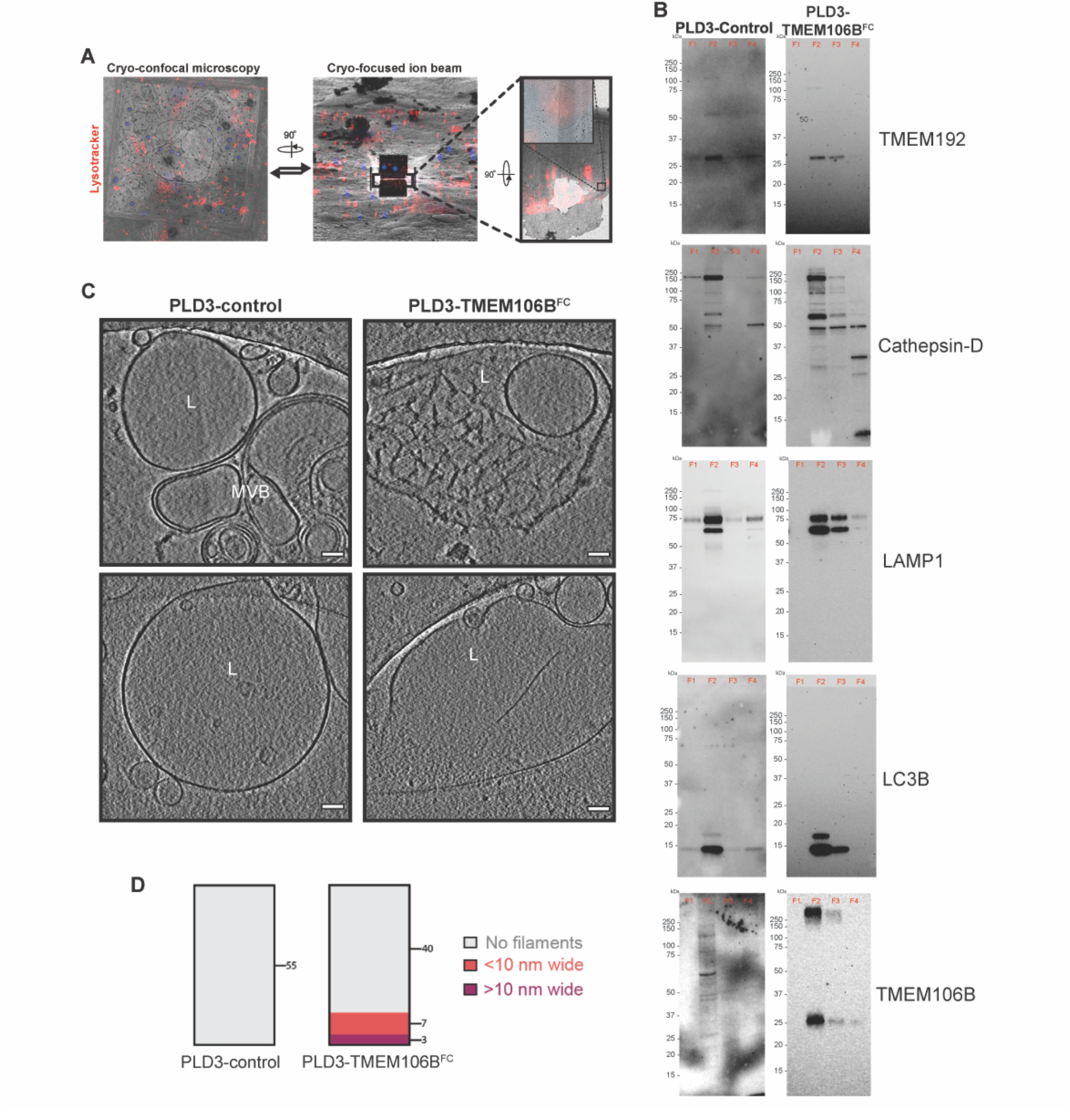
PLD3-TMEM106B^FC^ model generates intralysosomal filaments. **(A)** iPSCs are differentiated into iNeurons on micropatterned electron microscopy grids (see ‘Materials and Methods’ section for details). The grids are manually screened under a widefield microscope for quality and subsequently plunge-frozen in liquid ethane. FluoSpheres (diameter 2 µm) are added immediately before plunge-freezing. Left: A maximum Intensity projection image of the z-stack obtained from cryo-confocal fluorescence microscopy of a grid square of a plunge-frozen grid. Image is pseudo-colored as follows: 2 µm FluoSpheres (blue), and LysoTracker (red). Scale bar: 20 µm. Middle: Overlay of the ion beam image with the cryo-fluorescence microscopy image in (A), highlighting precise 3D correlation and a 200 nm thick lamella of a neuron post cryo-FIB milling. Scale bar: 20 µm. Right: Overlay of the cryo-fluorescence image with signal for LysoTracker (red) on a medium magnification cryo-transmission electron microscopy image (6500×). The black rectangle highlights the region selected for tilt-series acquisition and shows a zoom-in of the area (inset). Note: The slice presented is rotated 90° relative to the lamella view (middle). **(B)** Uncropped western blots of different gradient fractions probed for lysosomal proteins. Fraction 2 was identified as the lysosomal fraction for further interrogation (see Fig. 2). **(C)** Additional slices through cryo-tomograms of lysosomes from fraction 2 of gradient fractionation of iNeurons expressing PLD3-alone control or PLD3-TMEM106B. L: Lysosome; MVB: Multivesicular Body **(D)** Quantification of the percent of lysosomes inspected from fraction 2 that contain fibrillar structures, indicating whether the fibrils appear to be thick (>10 nm in width, purple) or thin (<10 nm in width, red). The number of lysosomes examined is indicated on the graph.

**Figure S4:**
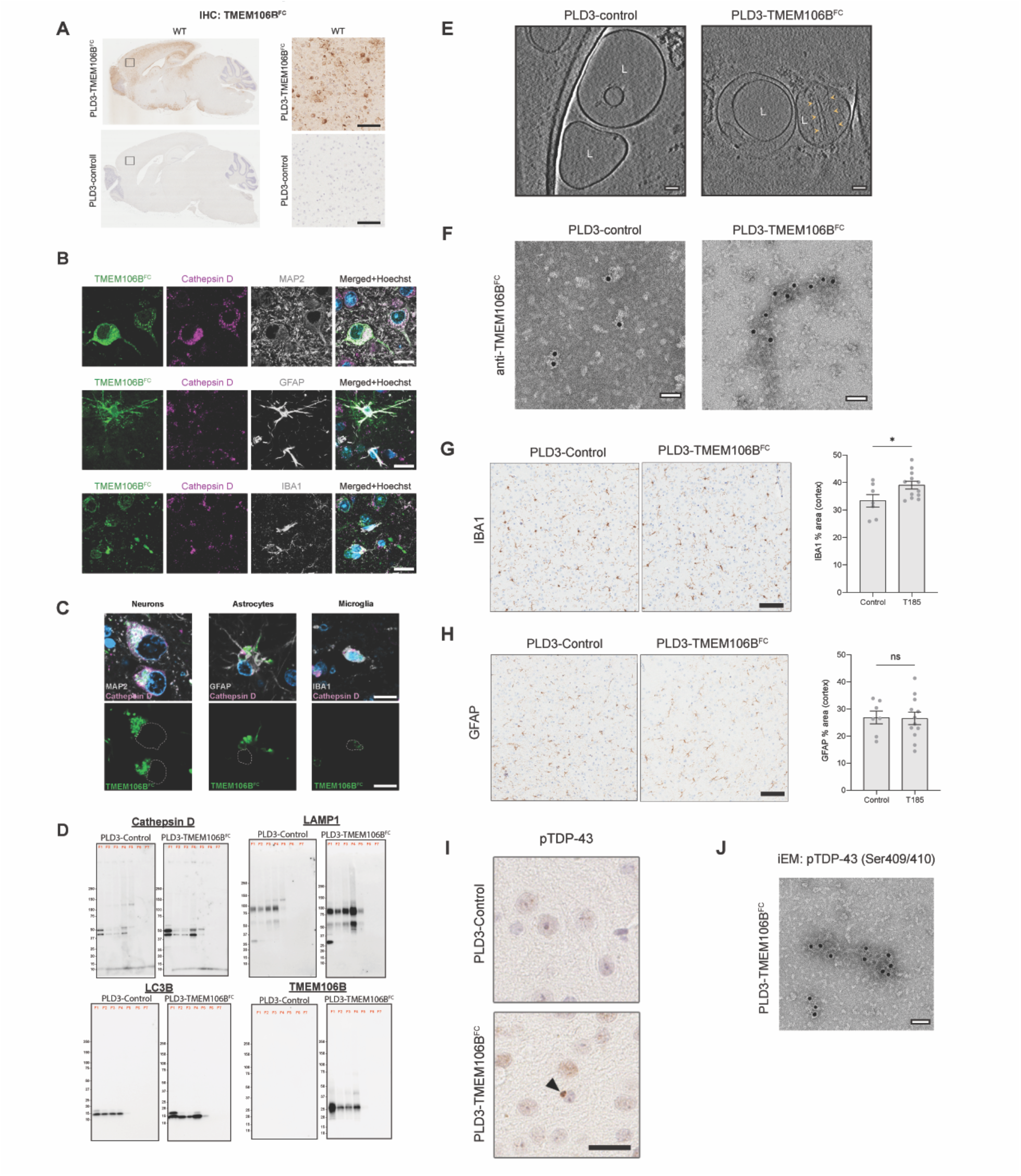
PLD3-TMEM106B^FC^ develop phenotypes of neurodegeneration. **(A)** Representative TMEM106B^FC^ immunohistochemistry (IHC) demonstrating expression in numerous brain regions vs control. Small black rectangles on left panels are magnified in right panels. Scale bar: 100 µm. **(B)** Immunofluorescence of mouse brain from mice expressing PLD3-TMEM106B^FC^. TMEM106B^FC^ stained along with CTSD to mark lysosomes and cell type markers for neurons (MAP2), astrocytes (GFAP), and microglia (IBA1). Scale bar: 10 µm. (**C)** Top row: co-immunofluorescent imaging in post-mortem FTLD brain depicting colocalization of TMEM106B fibril core (green) and cathepsin D (magenta) within neurons (MAP2, gray) and astrocytes (GFAP, gray), though very little TMEM106B-FC is observed in microglia (IBA1, gray). Bottom row: TMEM106B fibril core signal as detected using a cleaved TMEM106B fibril core-specific antibody (CST #87145). Nucleus tracings represented by white dashed lines. Scale bars: 10 µm. **(D)** Western blots of density gradient centrifugation fractions, probed for lysosomal markers to identify the lysosome-enriched fraction from mouse brains expressing PLD3-alone control or PLD3-TMEM106B^FC^. **(E)** Slices through reconstructed and denoised cryo-tomograms of fraction F4 from the brain of PLD3-alone control or PLD3-TMEM106B^FC^ mice, revealing fibrils within single-membraned lysosomes in the PLD3-TMEM106B expressing animals (L). Arrowheads depict the filaments. L: Lysosome. **(F)** Representative negative stain electron microscopy images of anti-TMEM106B^FC^ immuno-gold labeled sarkosyl-insoluble content from lysosomes isolated from PLD3-control or PLD3-TMEM106B-expressing mice. Scale bar: 100 nm. **(G)** IHC and quantification of IBA1 staining from mice expressing PLD3-TMEM106B^FC^ or PLD3-alone control. Statistics by Student’s t-test. N=7 control, 12 PLD3-TMEM106B^FC^. Scale bar: 100 µm. **(H)** IHC and quantification of GFAP staining from mice expressing PLD3-TMEM106B^FC^ or PLD3-alone control. Statistics by Student’s t-test. N=7 control, 12 PLD3-TMEM106B^FC^. Scale bar: 100 µm. **(I)** Representative IHC of pTDP-43 from frontal cortex of mice expressing PLD3-TMEM106B^FC^ or PLD3-alone control. Scale bar: 20 µm. **(J)** Representative negative stain electron microscopy image of anti-pTDP-43 immunogold-labeled fibrils isolated from the sarkosyl-insoluble fraction of lysosomes extracted from a PLD3-TMEM106B^FC^ expressing mouse. Scale bar: 50 nm.

**Figure S5:**
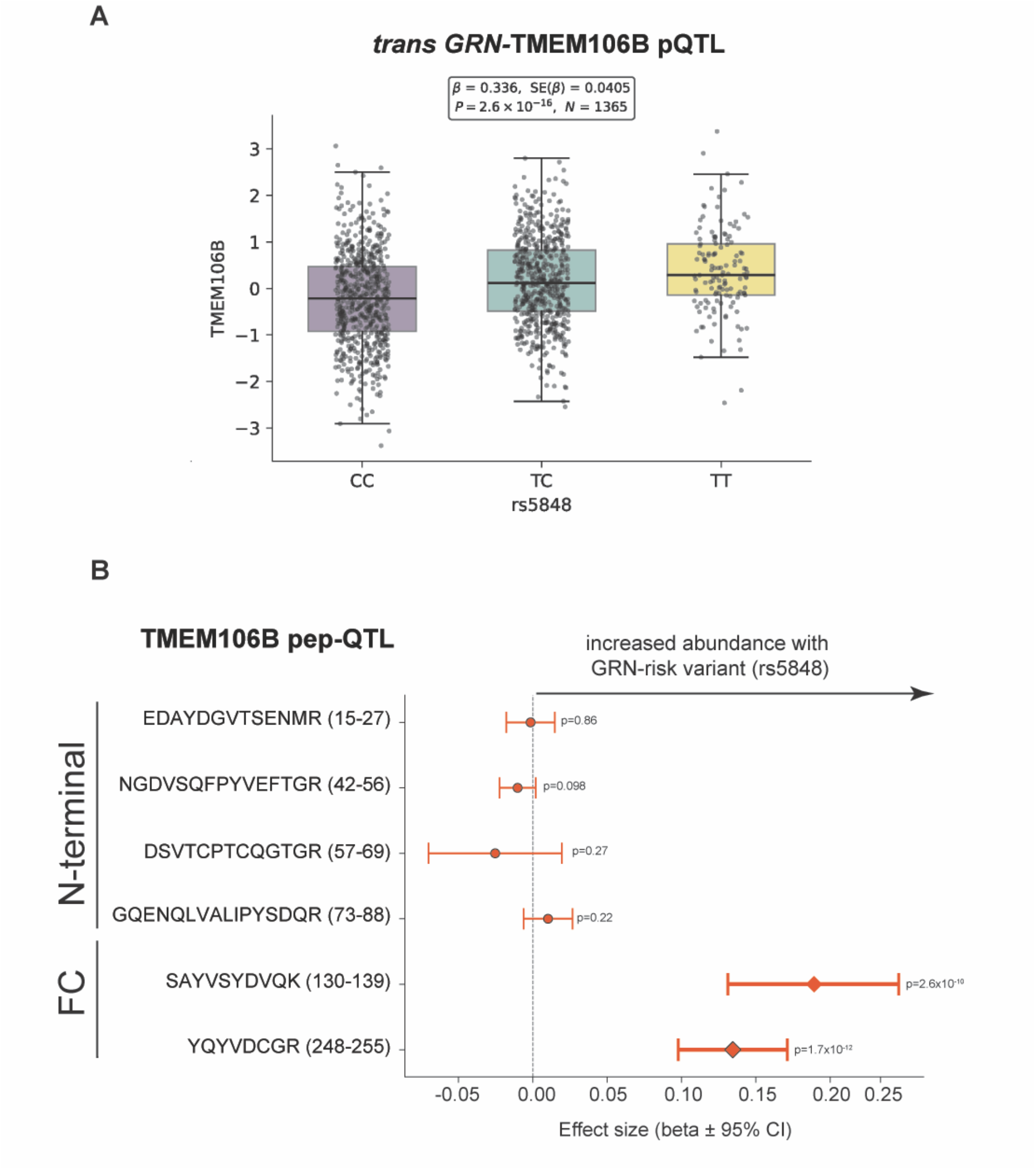
*Trans*-Association between TMEM106B peptides and *GRN* rs5848. **(A)** *Trans* pQTL analyses of *GRN* rs5848 genotype versus TMEM106B protein levels in post-mortem human brain (*35*). **(B)** *Trans*-peptide-QTL analyses for each individual TMEM106B peptide well-represented in the ROSMAP data. For each peptide, the association between peptide abundance and the *GRN* rs5848 genotype was measured. P-value indicated next to each bar; Effect size indicated on the x-axis. Positive effect size indicates increased abundance with the AD risk-associated minor allele *T*. FC: Fibril Core.

**Figure S6:**
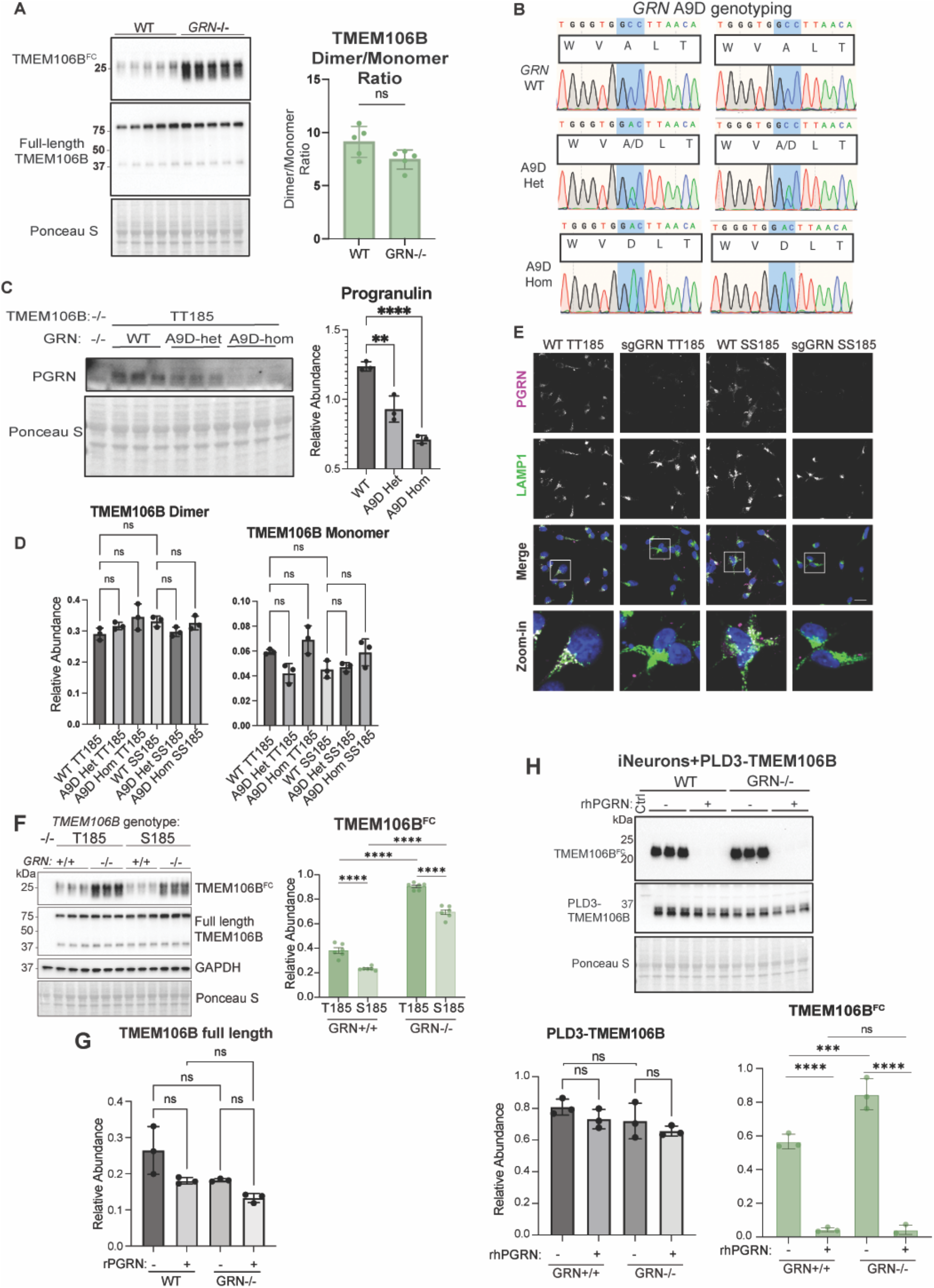
Progranulin regulates TMEM106B^FC^ accumulation in iNeurons. **(A)** Left: western blot for TMEM106B^FC^ and full-length TMEM106B of iNeuron lysates from WT or *GRN*-/-KOLF2.1J. Ponceau S as loading control. Right: Quantification on the right of TMEM106B dimer normalized to TMEM106B monomer. Student’s t-test. N = 5. Error bars = SD. **(B)** *GRN* gene Sanger sequencing data from KOLF2.1J *TMEM106B* T/T185 (left) or KOLF 2.1J *TMEM106B* S/S185 (right) cell lines engineered to be wild-type (top), heterozygous for the A9D mutation (middle), or homozygous for the A9D mutation (bottom). **(C)** Left: western blot for PGRN protein from KOLF 2.1J *TMEM106B* T/T185 cell line engineered to have wild-type, A9D/+ (heterozygous), or A9D/A9D homozygous *GRN* mutations. Right: quantification of PGRN protein normalized to Ponceau S. Student’s T-test. N=3. Error bars = SD. **(D)** Quantification of TMEM106B dimer band (left) and monomer band (right) in Western blot from Fig. 2G, normalized to Ponceau S. One-Way ANOVA with Šídák’s multiple comparisons test. N = 3 per group. Error bars = SD. **(E)** Immunofluorescence of iNeurons from KOLF2.1J *TMEM106B* T/T185 *GRN*-WT; KOLF2.1J *TMEM106B* T/T185 *GRN*-/-, KOLF2.1J *TMEM106B* S/S185 *GRN*-WT, and KOLF2.1J *TMEM106B* S/S185 *GRN*-/-cells with an anti-PGRN (magenta in merge) and anti-LAMP1 (green in merge) antibody. Nucleus in blue in merge. Scale bar 20µm. **(F)** Left: western blots from isogenic WT and *GRN*-/-iNeurons engineered to also express *TMEM106B* T/T185 or S/S185. Right: quantification of TMEM106B^FC^ normalized to Ponceau S. One-way ANOVA with Šídák’s multiple comparisons test. N = 3. Error bars = SD. **(G)** Quantification of Western blot in F. Relative abundance of full-length TMEM106B (sum of monomer and dimer) normalized to the lane’s Ponceau S signal. One-way ANOVA with Šídák’s multiple comparisons test. N=3. Error bars = SD. **(H)** Top: Western blot of lysates from either WT or *GRN*-/-iNeurons expressing PLD3-TMEM106B^FC^ (T185) that are treated with or without rhPGRN in their media throughout the differentiation. Bottom: Quantification of full-length PLD3-TMEM106B^FC^ (left) or TMEM106B^FC^ (right) normalized to the lane’s Ponceau S signal. One-way ANOVA with Šídák’s multiple comparisons test. N = 3. Error bars = SD. \**P* ≤ 0.05, \*\**P* ≤ 0.01, \*\*\**P* ≤ 0.001, \*\*\*\**P* ≤ 0.0001, ns: nonsignificant.

**Figure S7.**
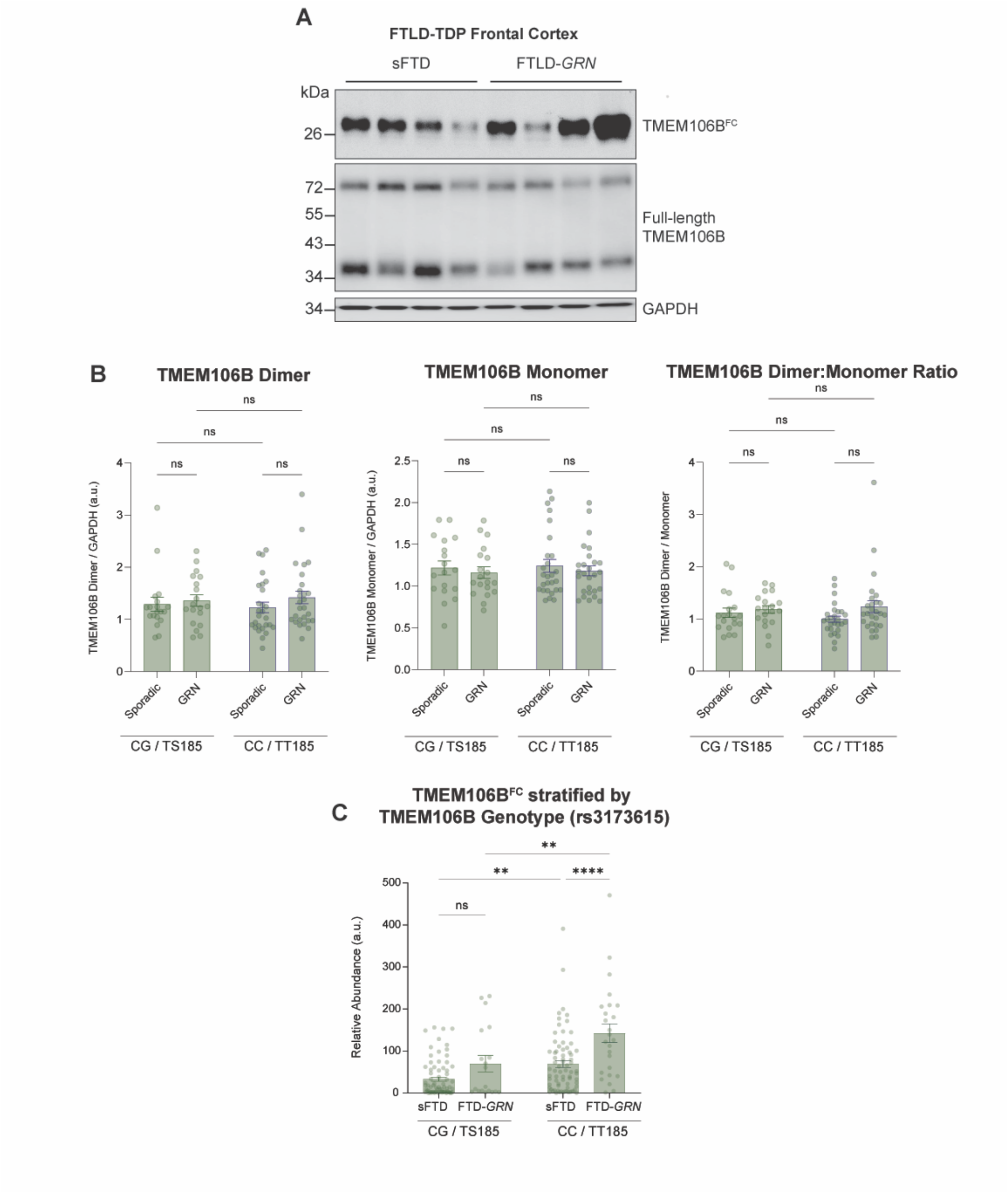
GRN and TMEM106B variants alter the accumulation of the TMEM106BFC in FTLD. **(A)** Representative Western blot of RIPA-soluble full-length TMEM106B and sarkosyl-insoluble TMEM106BFC frontal cortex from sFTLD and FTLD-*GRN* using cases initially characterized in Marks et al. (2024). **(B)** Quantification of TMEM106B dimer levels (left) and monomer levels (middle) normalized to GAPDH and dimer:monomer ratio (right) from immunoblots from FTLD-patient frontal cortex. Statistics by two-way ANOVA. **(C)** Quantification of TMEM106BFC levels stratified by both FTLD-*GRN* vs. sFTLD disease category and *TMEM106B* genotype. Statistics by two-way ANOVA with HC3 heteroscedasticity-consistent standard errors and simple-effects contrasts. *P ≤ 0.05, **P ≤ 0.01, ***P ≤ 0.001, ****P ≤ 0.0001, ns: nonsignificant.

**Figure S8.**
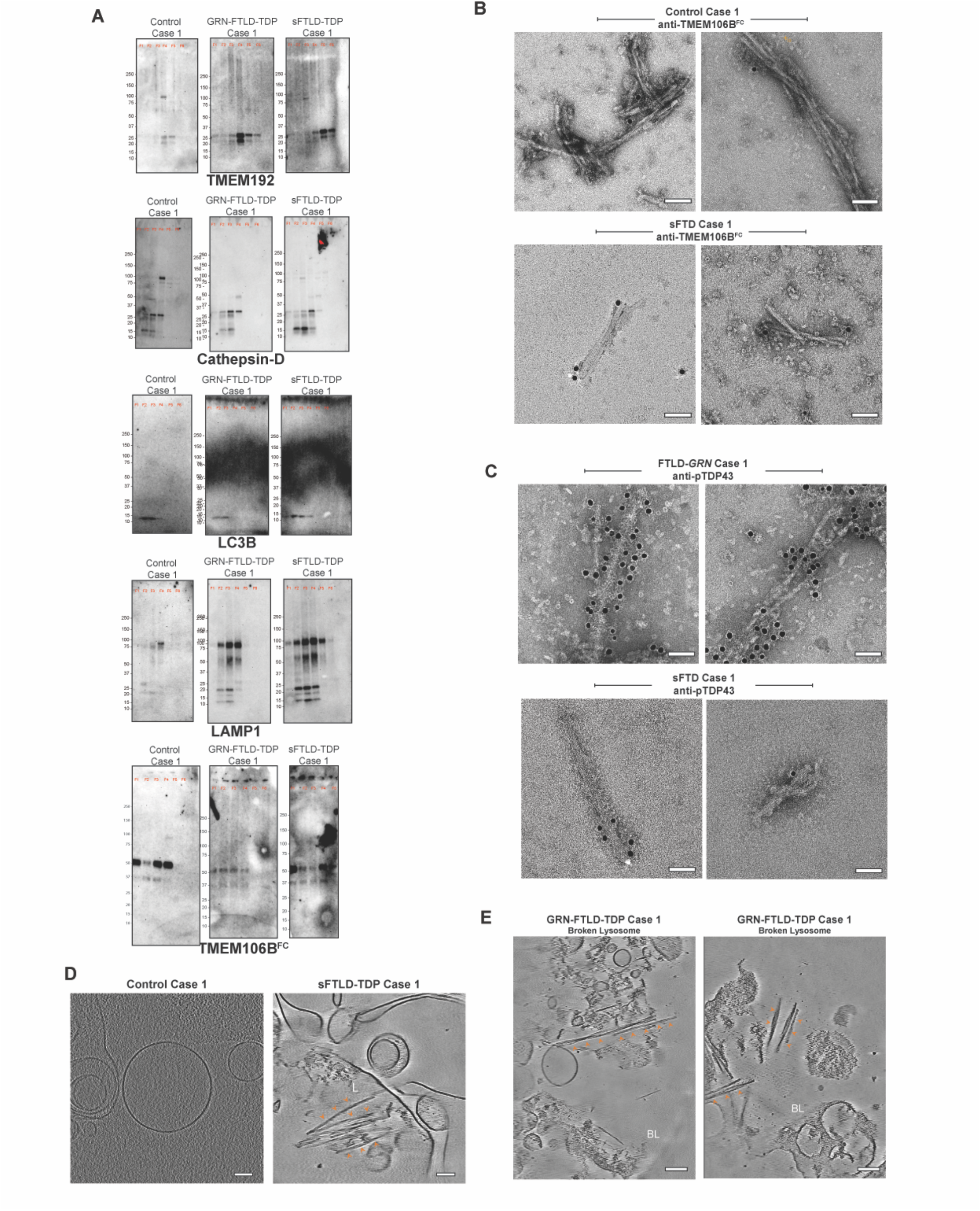
Fibrils are present within lysosomes from sFTLD and FTLD-*GRN* patient brain. **(A)** Western blots of fractions from density gradient centrifugation probed for lysosomal markers to identify lysosome-enriched fractions from human brain cases of healthy control brain, sFTLD, and FTLD-*GRN*. **(B)** Representative negative stain electron microscopy images of anti-TMEM106B^FC^ immunogold-labeled sarkosyl-insoluble lysosomal content isolated from a control (top) and a sporadic FTLD case (bottom). Scale bar: 50 nm. **(C)** Representative negative stain electron microscopy images of anti-pTDP-43 (Ser409/410) immunogold-labeled sarkosyl-insoluble lysosomal content isolated from a FTLD-*GRN* case (top) and a sporadic FTLD case (bottom). Scale bar: 50 nm. **(D)** Slices through reconstructed and denoised cryo-tomograms of fraction F4 from healthy-case (left) and sFTLD-case (right), revealing TMEM106B filaments within single-membraned lysosomes (L) in sFTLD-cases. Arrowheads depict the TMEM106B doublets with crossovers. Scale bar: 100 nm. **(D)** Slices through reconstructed and denoised cryo-tomograms of fraction F4 from FTLD-*GRN* case, revealing TMEM106B in broken lysosomes (BL). Arrowheads depict the TMEM106B doublets with crossovers. Arrowheads indicate TMEM106B doublets (orange). Scale bar: 100 nm.

**Figure S9.**
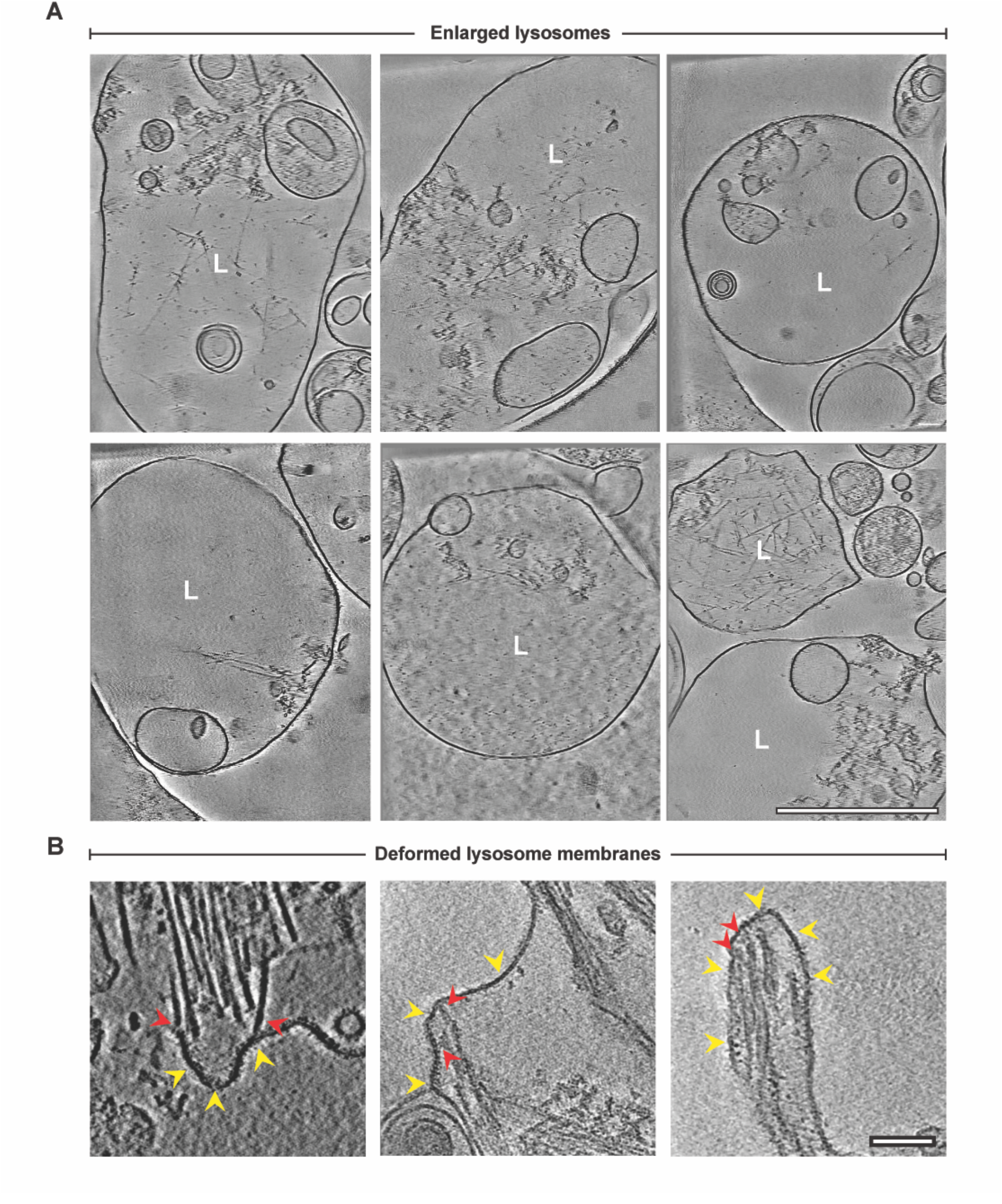
TMEM106B fibrils within large lysosomes and in contact with the lysosomal membrane. **(A)** Slices through reconstructed and denoised cryo-tomograms of lysosomes from sFTLD cases, demonstrating the very large size that these lysosomes can reach without breaking under the gradient centrifugation protocol. Scale bar: 500 nm. **(B)** Slices through reconstructed and denoised cryo-tomograms of lysosomes from sFTLD cases that show fibrils deforming lysosomal membranes. Fibrils contacting membranes are shown with red arrowheads and the lysosomal limiting membrane is highlighted in yellow. Scale bar: 100 nm.

**Figure S10.**
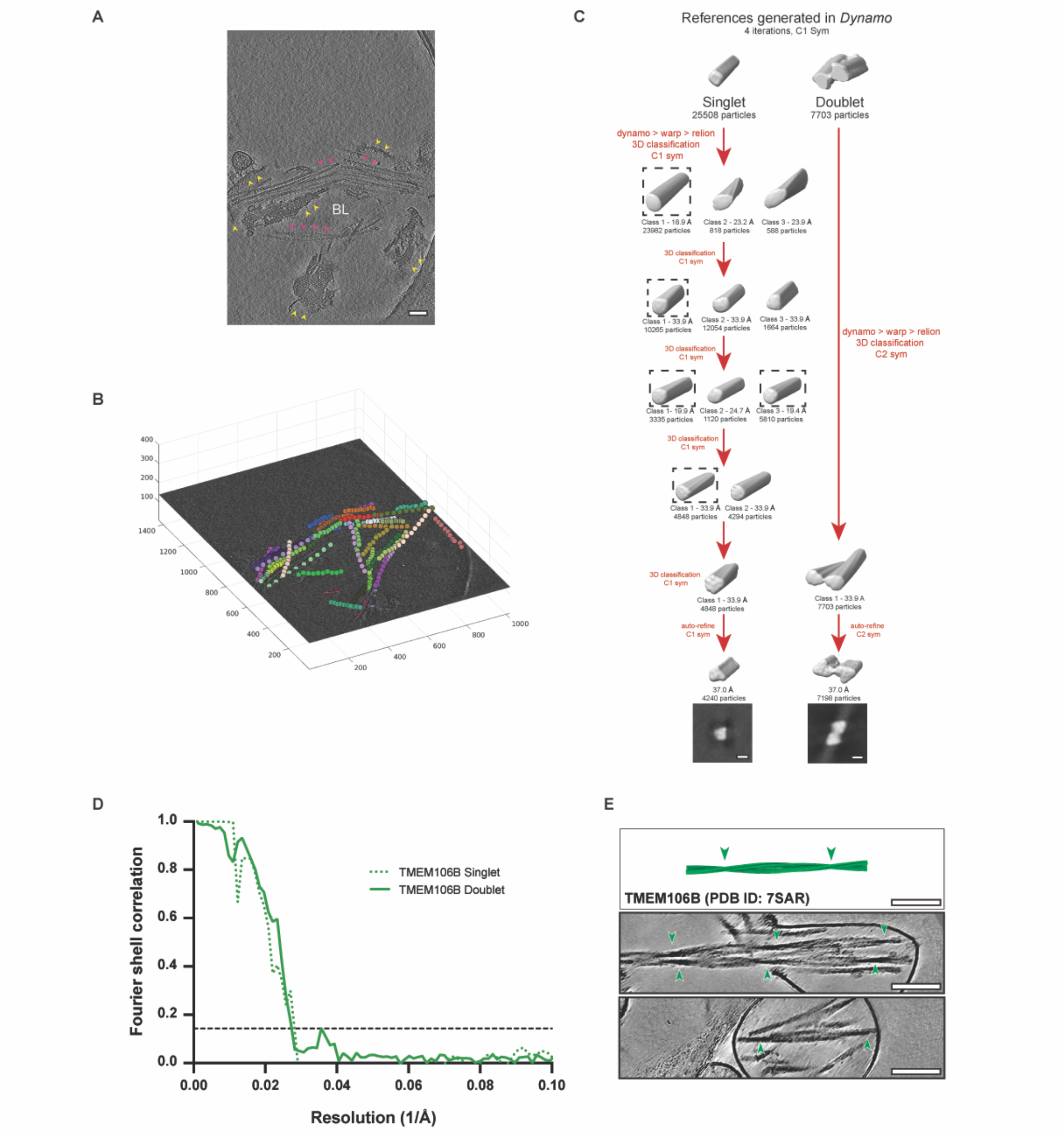
Characteristics and structure of intralysosomal fibrils demonstrate that they are composed of TMEM106B. **(A)** Slice through a representative cryo-tomogram of lysosome enriched from patients with FTLD-GRN. Red arrowheads point to the fibrils and yellow arrowheads indicate broken membranes. Scale bar: 100 nm. **(B)** 3D Representation of coordinates of the subtomograms extracted from the fibril densities in the tomogram in (A). Each filament is represented in a unique color and beads show the coordinates. **(C)** Outline of the schematic used for subtomogram averaging. Briefly, subvolumes/tomograms were extracted (from FTLD-GRN tomograms only) and aligned first in Dynamo followed by extraction of the aligned particles in Warp. The particles were imported into Relion and iteratively classified to obtain final set of good particles that were refined to obtain the density maps. **(D)** Gold-standard Fourier shell correlation (FSC) curves of the TMEM106B singlet (green dashed line), and TMEM106B doublet (green solid line) obtained from subtomogram averaging. The resolution as determined by the FSC0.143 cutoff is 37.0 Å for both TMEM106B singlet and doublet. **(E)** Top: Pseudo-TMEM106B fibril model comprised of 750 repeats generated in chimeraX (chains A, B of deposited PDB model 7SAR), known helical symmetry parameters: rise (4.78Å) and twist (-0.41°). Middle and bottom panels: Two representative examples of TMEM106B doublets within the broken or intact lysosomes isolated from patient tissue. Arrowheads depict the crossover of TMEM106B doublets. Scale bar: 100 nm.

## References

1. A. M. Cuervo, J. F. Dice, When lysosomes get old. Exp. Gerontol. 35, 119–31 (2000).

2. R. A. Nixon, The aging lysosome: An essential catalyst for late-onset neurodegenerative diseases. Biochim. Biophys. Acta Proteins Proteom. 1868, 140443 (2020).

3. R. A. Nixon, D. C. Rubinsztein, Mechanisms of autophagy-lysosome dysfunction in neurodegenerative diseases. Nat. Rev. Mol. Cell Biol. 25, 926–946 (2024).

4. L. Guerrero-Navarro, P. Jansen-Dürr, M. Cavinato, Age-Related Lysosomal Dysfunctions. Cells 11 (2022).

5. A. K. H. Stavoe, E. L. F. Holzbaur, Autophagy in Neurons. Annu. Rev. Cell Dev. Biol. 35, 477–500 (2019).

6. C. Soto, L. D. Estrada, Protein misfolding and neurodegeneration. Arch. Neurol. 65, 184–9 (2008).

7. E. Chuang, A. M. Hori, C. D. Hesketh, J. Shorter, Amyloid assembly and disassembly. J. Cell Sci. 131, jcs189928 (2018).

8. S. H. W. Scheres, B. Ryskeldi-Falcon, M. Goedert, Molecular pathology of neurodegenerative diseases by cryo-EM of amyloids. Nature 621, 701–710 (2023).

9. T. W. Todd et al., Cryo-EM structures of pathogenic fibrils and their impact on neurodegenerative disease research. Neuron 112, 2269–2288 (2024).

10. I. Riera-Tur, T. Schäfer, D. Hornburg, A. Mishra, et al., Amyloid-like aggregating proteins cause lysosomal defects in neurons via gain-of-function toxicity. Life Sci. Alliance 5 (2022).

11. N. Tesi, S. et al., Cognitively healthy centenarians are genetically protected against Alzheimer’s disease. Alzheimers Dement. 20, 3864–3875 (2024).

12. M. P. Szabo, S. Mishra, A. Knupp, J. E. Young, The role of Alzheimer’s disease risk genes in endolysosomal pathways. Neurobiol. Dis. 162 (2022).

13. J. Root et al., Lysosome dysfunction as a cause of neurodegenerative diseases: Lessons from frontotemporal dementia and amyotrophic lateral sclerosis. Neurobiol. Dis. 154, 105360 (2021).

14. J. A. Tunold et al., Lysosomal Polygenic Burden Drives Cognitive Decline in Parkinson’s Disease with Low Alzheimer Risk. Mov. Disord. 39, 596–601 (2024).

15. V. M. Van Deerlin et al., Common variants at 7p21 are associated with frontotemporal lobar degeneration with TDP-43 inclusions. Nat. Genet. 42, 234–239 (2010).

16. M. Baker et al., Mutations in progranulin cause tau-negative frontotemporal dementia linked to chromosome 17. Nature 442, 916–919 (2006).

17. M. Cruts et al., Null mutations in progranulin cause ubiquitin-positive frontotemporal dementia linked to chromosome 17q21. Nature 442, 920–924 (2006).

18. J. Gass et al., Mutations in progranulin are a major cause of ubiquitin-positive frontotemporal lobar degeneration. Hum. Mol. Genet. 15, 2988–3001 (2006).

19. Z. Li et al., The TMEM106B FTLD-protective variant, rs1990621, is also associated with increased neuronal proportion. Acta Neuropathol. 139, 45–61 (2020).

20. C. Bellenguez et al., New insights into the genetic etiology of Alzheimer’s disease and related dementias. Nat. Genet. 54, 412–436 (2022).

21. S. S. Shankaran et al., Missense mutations in the progranulin gene linked to frontotemporal lobar degeneration with ubiquitin-immunoreactive inclusions reduce progranulin production and secretion. J. Biol. Chem. 283, 1744–1753 (2008).

22. L. Currens et al., A case of familial frontotemporal dementia caused by a progranulin gene mutation. Clin. Park. Relat. Disord. 9, 100213 (2023).

23. N. Finch et al., TMEM106B regulates progranulin levels and the penetrance of FTLD in GRN mutation carriers. Neurology 76, 467–74 (2011).

24. C. Cruchaga et al., Association of TMEM106B gene polymorphism with age at onset in granulin mutation carriers and plasma granulin protein levels. Arch. Neurol. 68, 581–586 (2011).

25. H. S. Yang et al., Genetics of Gene Expression in the Aging Human Brain Reveal TDP-43 Proteinopathy Pathophysiology. Neuron 107, 496–508.e6 (2020).

26. A. Chang et al., Homotypic fibrillization of TMEM106B across diverse neurodegenerative diseases. Cell 185, 1346–1355.e15 (2022).

27. Y. X. Jiang et al., Amyloid fibrils in FTLD-TDP are composed of TMEM106B and not TDP-43. Nature 605, 304–309 (2022).

28. M. Schweighauser et al., Age-dependent formation of TMEM106B amyloid filaments in human brains. Nature 605, 310–314 (2022).

29. Y. Fan et al., Generic amyloid fibrillation of TMEM106B in patient with Parkinson’s disease dementia and normal elders. Cell Res. 32, 585–588 (2022).

30. S. Held et al., Physiological shedding and C-terminal proteolytic processing of TMEM106B. Cell Rep. 44 (2025).

31. J. D. Marks et al., TMEM106B core deposition associates with TDP-43 pathology and is increased in risk SNP carriers for frontotemporal dementia. Sci. Transl. Med. 16, eadf9735 (2024).

32. C. T. Vicente et al., C-terminal TMEM106B fragments in human brain correlate with disease-associated TMEM106B haplotypes. Brain 146, 4055–4064 (2023).

33. H. Holstege et al., Fine-mapping of the TMEM106B locus reveals four haplotypes that are differentially associated with risk for neurodegeneration. Alzheimers Dement. 21 (2025).

34. A. P. Wingo et al., Multiancestry brain pQTL fine-mapping and integration with genome-wide association studies of 21 neurologic and psychiatric conditions. Nat. Genet. 57, 2156–2165 (2025).

35. Wingo et al., In preparation (2026).

36. J. Dai et al., Effects of APOE genotype on brain proteomic network and cell type changes in Alzheimer’s disease. Front. Mol. Neurosci. 11 (2018).

37. D. A. Bennett et al., Religious Orders Study and Rush Memory and Aging Project. J. Alzheimers Dis. 64, S161–S189 (2018).

38. L. Higginbotham et al., Integrated proteomics reveals brain-based cerebrospinal fluid biomarkers in asymptomatic and symptomatic Alzheimer’s disease. Sci. Adv. 6 (2020).

39. A. M. Frankenfield et al., A Deep Quantitative Proteome Turnover Platform for Human iPSC-derived Neurons. bioRxiv, doi: 10.64898/2026.03.14.711828 (2026).

40. C. Wang et al., Scalable Production of iPSC-Derived Human Neurons to Identify Tau-Lowering Compounds by High-Content Screening. Stem Cell Reports 9, 1221–1233 (2017).

41. A. Rodney, K. Karanjeet, K. Benzow, M. D. Koob, A common Alu insertion in the 3’UTR of TMEM106B is associated with risk of dementia. Alzheimers Dement. 20, 5071–5077 (2024).

42. A. Chemparathy et al., A 3’UTR Insertion Is a Candidate Causal Variant at the TMEM106B Locus Associated With Increased Risk for FTLD-TDP. Neurol. Genet. 10, e200124 (2024).

43. H. Suzuki, M. Matsuoka, The lysosomal trafficking transmembrane protein 106B is linked to cell death. J. Biol. Chem. 291, 21448–21460 (2016).

44. J. Perneel et al., Increased TMEM106B levels lead to lysosomal dysfunction which affects synaptic signaling and neuronal health. Mol. Neurodegener. 20 (2025).

45. X. Zhou, L. Sun, O. A. Brady, K. A. Murphy, F. Hu, Elevated TMEM106B levels exaggerate lipofuscin accumulation and lysosomal dysfunction in aged mice with progranulin deficiency. Acta Neuropathol. Commun. 5, 9 (2017).

46. A. C. Gonzalez et al., Unconventional Trafficking of Mammalian Phospholipase D3 to Lysosomes. Cell Rep. 22, 1040–1053 (2018).

47. A. L. Erwin et al., Molecular Visualization of Neuronal TDP43 Pathology In Situ. bioRxiv, doi: 10.1101/2024.08.19.608477 (2024).

48. M. Bacioglu et al., Cleaved TMEM106B forms amyloid aggregates in central and peripheral nervous systems. Acta Neuropathol. Commun. 12 (2024).

49. W. Zhao et al., Tracing TMEM106B fibril deposition in aging and Parkinson’s disease with dementia brains. Life Med. 3, nae011 (2024).

50. E. R. Gallagher, E. L. F. Holzbaur, The selective autophagy adaptor p62/SQSTM1 forms phase condensates regulated by HSP27 that facilitate the clearance of damaged lysosomes via lysophagy. Cell Rep. 42 (2023).

51. R. M. Ransohoff, How neuroinflammation contributes to neurodegeneration. Science 353, 777–83 (2016).

52. A. Meneses et al., TDP-43 Pathology in Alzheimer’s Disease. Mol. Neurodegener. 16, 84 (2021).

53. P. T. Nelson et al., Limbic-predominant age-related TDP-43 encephalopathy (LATE): consensus working group report. Brain 142, 1503–1527 (2019).

54. A. M. Nicholson, R. Rademakers, What we know about TMEM106B in neurodegeneration. Acta Neuropathol. 132, 639–651 (2016).

55. M. Khalil et al., Neurofilaments as biomarkers in neurological disorders. Nat. Rev. Neurol. 14, 577–589 (2018).

56. M. Khalil et al., Neurofilaments as biomarkers in neurological disorders - towards clinical application. Nat. Rev. Neurol. 20, 269–287 (2024).

57. J. Perneel, M. Manoochehri, E. D. Huey, R. Rademakers, J. Goldman, Case report: TMEM106B haplotype alters penetrance of GRN mutation in frontotemporal dementia family. Front. Neurol. 14 (2023).

58. S. Lattante et al., Defining the association of TMEM106B variants among frontotemporal lobar degeneration patients with GRN mutations and C9orf72 repeat expansions. Neurobiol. Aging 35, 2658.e1–2658.e5 (2014).

59. S. S. Mirza et al., Disease-modifying effects of TMEM106B in genetic frontotemporal dementia: a longitudinal GENFI study. Brain 148, 2746–2762 (2025).

60. O. Mukherjee et al., Molecular characterization of novel progranulin (GRN) mutations in frontotemporal dementia. Hum. Mutat. 29, 512–21 (2008).

61. M. A. Nalls et al., Evidence for GRN connecting multiple neurodegenerative diseases. Brain Commun. 3, fcab095 (2021).

62. P. Reho et al., GRN Mutations Are Associated with Lewy Body Dementia. Mov. Disord. 37, 1943–1948 (2022).

63. Z. Ahmed et al., Accelerated lipofuscinosis and ubiquitination in granulin knockout mice suggest a role for progranulin in successful aging. Am. J. Pathol. 177, 311–324 (2010).

64. C. Fenoglio et al., Rs5848 variant influences GRN mRNA levels in brain and peripheral mononuclear cells in patients with Alzheimer’s disease. J. Alzheimers Dis. 18, 603–12 (2009).

65. B. B. Sun et al., Genomic atlas of the human plasma proteome. Nature 558, 73– 79 (2018).

66. A. P. Wingo et al., Integrating human brain proteomes with genome-wide association data implicates new proteins in Alzheimer’s disease pathogenesis. Nat. Genet. 53, 143–146 (2021).

67. T. Logan et al., Rescue of a lysosomal storage disorder caused by Grn loss of function with a brain penetrant progranulin biologic. Cell 184, 4651–4668.e25 (2021).

68. J. Root et al., Granulins rescue inflammation, lysosome dysfunction, lipofuscin, and neuropathology in a mouse model of progranulin deficiency. Cell Rep. 43 (2024).

69. T. Feng et al., Loss of TMEM106B and PGRN leads to severe lysosomal abnormalities and neurodegeneration in mice. EMBO Rep. 21 (2020).

70. J. Perneel, R. Rademakers, Identification of TMEM106B amyloid fibrils provides an updated view of TMEM106B biology in health and disease. Acta Neuropathol. 143, 527–529 (2022).

71. J. Y. Lee et al., The major TMEM106B dementia risk allele affects TMEM106B protein levels, fibril formation, and myelin lipid homeostasis in the ageing human hippocampus. Mol. Neurodegener. 18 (2023).

72. A. N. Salazar et al., An AluYb8 mobile element characterises the risk haplotype of the TMEM106B locus associated with neurodegeneration. Alzheimers Dement. 20 (2024).

73. M. D. Gallagher et al., A Dementia-Associated Risk Variant near TMEM106B Alters Chromatin Architecture and Gene Expression. Am. J. Hum. Genet. 101, 643–663 (2017).

74. S. Hasan et al., Multi-modal proteomic characterization of lysosomal function and proteostasis in progranulin-deficient neurons. Mol. Neurodegener. 18 (2023).

75. Passage Bio, Inc., A Study of PBFT02 in Participants With FTD and Mutations in the Granulin Precursor (GRN) or C9ORF72 Genes (upliFT-D). ClinicalTrials.gov (2021).

76. AviadoBio Ltd, A Study to Evaluate the Safety and Effect of AVB-101 in Subjects With FTD-GRN (ASPIRE-FTD). ClinicalTrials.gov (2023).

77. Prevail Therapeutics, Phase 1/2 Clinical Trial of LY3884963 in Patients With FTD-GRN (PROCLAIM). ClinicalTrials.gov (2020).

78. A. E. Arrant, V. C. Onyilo, D. E. Unger, E. D. Roberson, Progranulin gene therapy improves lysosomal dysfunction and microglial pathology associated with frontotemporal dementia and neuronal ceroid lipofuscinosis. J. Neurosci. 38, 2341–2358 (2018).

79. M. Kurnellas et al., Latozinemab, a novel progranulin-elevating therapy for frontotemporal dementia. J. Transl. Med.21 (2023).

80. S. S. Minami et al., Progranulin protects against amyloid β deposition and toxicity in Alzheimer’s disease mouse models. Nat. Med. 20, 1157–1164 (2014).

81. Y. Ciervo et al., Restoration of progranulin by engineered hematopoietic stem cell-derived microglia corrects phenotypes of granulin knockout mice. Sci. Transl. Med. 18, eadw9930 (2026).

82. K. Rose et al., Tau fibrils induce nanoscale membrane damage and nucleate cytosolic tau at lysosomes. bioRxiv, doi: 10.1101/2023.08.28.555157 (2023).

83. Y. X. Xie et al., Lysosomal exocytosis releases pathogenic α-synuclein species from neurons in synucleinopathy models. Nat. Commun. 13 (2022).

84. D. Li et al., Cathepsin-dependent amyloid formation drives mechanical rupture of lysosomal membranes. bioRxiv, doi: 10.64898/2026.01.17.700056 (2026).

85. R. D. Elias et al., Cathepsin C-Catalyzed Ligation Generates Intralysosomal Amyloid Fibrils from Dipeptide Esters. bioRxiv, doi: 10.64898/2025.12.23.696283 (2025).

86. G. C. Werthmann, J. Herz, Distinct lysosomal dysfunction patterns of GRN deficiency in the CNS implicate progranulin in cell type-specific protein sorting. bioRxiv, doi: 10.64898/2025.12.29.696915 (2026).

87. A. S. Kim, J. Z. Wu, R. Cai, S. A. Freeman, Mechanoresilience of lysosomes conferred by TMEM63A. J. Cell Biol. 225 (2026).

88. S. Pankiv et al., p62/SQSTM1 binds directly to Atg8/LC3 to facilitate degradation of ubiquitinated protein aggregates by autophagy. J. Biol. Chem. 282, 24131– 24145 (2007).

89. K. Kakuda et al., Lysophagy protects against propagation of α-synuclein aggregation through ruptured lysosomal vesicles. Proc. Natl. Acad. Sci. U.S.A. 121 (2024).

90. P. J. Sampognaro et al., Mutations in α-synuclein, TDP-43 and tau prolong protein half-life through diminished degradation by lysosomal proteases. Mol. Neurodegener. 18 (2023).

91. W. Zhong et al., Lysosomal escape and TMEM106B fibrillar core determine TDP-43 seeding outcomes. bioRxiv, doi: 10.64898/2025.12.19.695531 (2025).

92. M. S. Fernandopulle et al., Transcription Factor–Mediated Differentiation of Human iPSCs into Neurons. Curr. Protoc. Cell Biol. 79 (2018).

93. A. M. Frankenfield, J. Ni, M. Ahmed, L. Hao, Protein Contaminants Matter: Building Universal Protein Contaminant Libraries for DDA and DIA Proteomics. J. Proteome Res. 21, 2104–2113 (2022).

94. V. H. Ryan et al., Maintenance of neuronal TDP-43 expression requires axonal lysosome transport. eLife 14 (2025).

95. A. M. Frankenfield et al., Benchmarking SILAC Proteomics Workflows and Data Analysis Platforms. Mol. Cell. Proteomics 24 (2025).

96. T. Uenaka et al., Prevention of transgene silencing during human pluripotent stem cell differentiation. Cell Stem Cell, doi: 10.1016/j.stem.2026.01.007 (2026).

97. J. Gass et al., Progranulin regulates neuronal outgrowth independent of Sortilin. Mol. Neurodegener. 7 (2012).

98. S. Das et al., Next-generation genotype imputation service and methods. Nat. Genet. 48, 1284–1287 (2016).

99. A. Manichaikul et al., Robust relationship inference in genome-wide association studies. Bioinformatics 26, 2867–73 (2010).

100. A. L. Price et al., Principal components analysis corrects for stratification in genome-wide association studies. Nat. Genet. 38, 904–909 (2006).

101. A. T. Kong et al., MSFragger: ultrafast and comprehensive peptide identification in mass spectrometry-based proteomics. Nat. Methods 14, 513–520 (2017).

102. G. A. Khoury, R. C. Baliban, C. A. Floudas, Proteome-wide post-translational modification statistics: frequency analysis and curation of the swiss-prot database. Sci. Rep. 1 (2011).

103. L. Käll et al., Semi-supervised learning for peptide identification from shotgun proteomics datasets. Nat. Methods 4, 923–5 (2007).

104. J. T. Leek, J. D. Storey, Capturing heterogeneity in gene expression studies by surrogate variable analysis. PLoS Genet. 3, 1724–35 (2007).

105. E. B. Dammer, N. T. Seyfried, E. C. B. Johnson, Batch Correction and Harmonization of -Omics Datasets with a Tunable Median Polish of Ratio. Front. Syst. Biol. 3 (2023).

106. C. C. Chang et al., Second-generation PLINK: rising to the challenge of larger and richer datasets. Gigascience 4, 7 (2015).

107. A. P. Wingo et al., Integrating human brain proteomes with genome-wide association data implicates new proteins in Alzheimer’s disease pathogenesis. Nat. Genet. 53, 143–146 (2021).

108. L. Yu et al., Cortical Proteins Associated With Cognitive Resilience in Community-Dwelling Older Persons. JAMA Psychiatry 77, 1172–1180 (2020).

109. J. Chew et al., C9ORF72 repeat expansions in mice cause TDP-43 pathology, neuronal loss, and behavioral deficits. Science 348, 1151–1154 (2015).

110. J. Arnold et al., Site-Specific Cryo-focused Ion Beam Sample Preparation Guided by 3D Correlative Microscopy. Biophys. J. 110, 860–869 (2016).

111. G. Wolff et al., Mind the gap: Micro-expansion joints drastically decrease the bending of FIB-milled cryo-lamellae. J. Struct. Biol. 208 (2019).

112. W. J. H. Hagen, W. Wan, J. A. G. Briggs, Implementation of a cryo-electron tomography tilt-scheme optimized for high resolution subtomogram averaging. J. Struct. Biol. 197, 191–198 (2017).

113. D. N. Mastronarde, Automated electron microscope tomography using robust prediction of specimen movements. J. Struct. Biol. 152, 36–51 (2005).

114. H. Bains, R. Singh, Isolation of autophagic fractions from mouse liver for biochemical analyses. STAR Protoc. 2 (2021).

115. Y. Fan et al., Generic amyloid fibrillation of TMEM106B in patient with Parkinson’s disease dementia and normal elders. Cell Res. 32, 585–588 (2022).

116. D. Tegunov, P. Cramer, Real-time cryo-electron microscopy data preprocessing with Warp. Nat. Methods 16, 1146–1152 (2019).

117. J. R. Kremer, D. N. Mastronarde, J. R. McIntosh, Computer visualization of three-dimensional image data using IMOD. J. Struct. Biol. 116, 71–76 (1996).

118. D. Castaño-Díez, M. Kudryashev, M. Arheit, H. Stahlberg, Dynamo: A flexible, user-friendly development tool for subtomogram averaging of cryo-EM data in high-performance computing environments. J. Struct. Biol. 178, 139–151 (2012).

119. A. Burt et al., A flexible framework for multi-particle refinement in cryo-electron tomography. PLoS Biol. 19 (2021).

120. A. Burt et al., An image processing pipeline for electron cryo-tomography in RELION-5. FEBS Open Bio 14, 1788–1804 (2024).

121. X. Zhang, J. Mahamid, Protocol for subtomogram averaging of helical filaments in cryo-electron tomography. STAR Protoc. 5 (2024).

122. E. F. Pettersen et al., UCSF ChimeraX: Structure visualization for researchers, educators, and developers. Protein Sci. 30, 70–82 (2021).

123. L. Lamm et al., MemBrain v2: an end-to-end tool for the analysis of membranes in cryo-electron tomography. bioRxiv, doi: 10.1101/2024.01.05.574336 (2024).

124. Seifar et al., Large-scale deep proteomic analysis in Alzheimer’s disease brain regions across race and ethnicity. Alzheimers Dement. 2024;20(12):8878–97. Epub 20241113. doi: 10.1002/alz.14360.

125. Wojtas et al., Multi-scale integration of human brain vascular and CSF proteomes reveals biomarkers of cerebral amyloid angiopathy linked to Alzheimer’s disease risk. medRxiv 2025.10.08.25337413; 10.1101/2025.10.08.25337413

