## Supplemental Methods for "Neurodegeneration risk variants promote lysosomal TMEM106B fibril accumulation"

**Materials and Methods**

**Cell culture and differentiations**

Induced Pluripotent Stem Cells (iPSCs) were maintained in Essential 8 (E8) medium (Life Technologies A1517001). Regular splitting of iPSCs was performed using accutase (Life Technologies A1110501) or phosphate-buffered saline (PBS)+0.5 mM EDTA, and cells were plated on Matrigel-coated plates with E8+50 nM Chroman1 (MedChemExpress HY-15392). Maintenance was performed following the protocols in Fernandopulle et al., 2018 (*92*).

On Day 0 of the differentiation, WTC11-based iPSCs (cell line name: i11W-mNC; Fig. 2) were split using accutase to singularize cells and re-plated on Matrigel-coated plates in Induction Media (Knock-out DMEM/F-12 [Life Technologies 12660012] supplemented with 1× N-2 [Life Technologies 17502048], 1% GlutaMax [Life Technologies 35050061], 1% NEAAs [Life Technologies 11140050], 2 μg/mL doxycycline [Takara Bio 631311], 50 nM Chroman1) at high confluence (50–70%). On the following two days, the media was changed to Induction Media without Chroman 1. On day 3, WTC11-based iNeurons were split using accutase and re-plated on poly-L-ornithine (PLO)-coated dishes for final experiments in Neural Maturation Media (“NMM”; BrainPhys [StemCell Technologies 05790] supplemented with 1× N21-Max [R&D Systems AR008], 10 ng/mL BDNF [Life Technologies 450-02-500UG], 10 ng/mL GDNF [Life Technologies 450-10-500UG], 10 ng/mL NT-3 [Life Technologies 450-03-500UG], 1 μg/mL laminin, and 2 μg/mL doxycycline). Thereafter, iNeurons received a half media change with NMM every 2–3 days.

KOLF2.1J-based iNeurons (Fig. 1 and Fig. 4) followed a similar induction as WTC11-based iNeurons, with the following changes. On Day 3, KOLF-based iNeurons were treated with Induction Media without Chroman 1, with 1 μM FdU/uridine as a mitotic inhibitor. On Day 4, KOLF-based iNeurons were passaged using accutase and re-plated on PLO-coated dishes in “Day 4+7 NMM” (50% Knockout DMEM/F12, 50% BrainPhys supplemented with N21-Max, 10 ng/mL BDNF, 10 ng/mL GDNF, 10 ng/mL NT-3, 1 μg/mL laminin, 1 μM FdU/uridine, and 2 μg/mL doxycycline). Typically, a complete media change was performed the day after plating (Day 5). On Day 7, half of the media was removed and replaced using “Day 4+7 NMM”. On Day 10 and beyond, “Day 10 NMM” (BrainPhys supplemented with 1× N21-Max, 10 ng/mL BDNF, 10 ng/mL GDNF, 10 ng/mL NT-3, 1 μg/mL laminin, 1 μM FdU/uridine, and 2 μg/mL doxycycline) was used for half media changes every 2–3 days. Collection timepoints of cells varied slightly but most commonly was performed on Days 14 (e.g., Fig. 2B and C), 16 (e.g., Fig. 1D and 4E and G), or 18 (e.g., Fig. 2J). In all iNeuron experiments, the number of replicates indicated in figure legends consists of biological replicates defined by samples that were separated from at least day 4 onward in the differentiation.

**dSILAC labeling in iNeurons**

Human iNeurons were maintained in DMEM for SILAC media with added 0.3 mM arginine and 0.5 mM lysine. All other media components were the same as in BrainPhys media. On Day 7 or 10 of differentiations, neurons were washed with PBS and media was switch to a media containing “heavy” arginine (L-Arginine:HCL [13C6, 99%; 15N4, 99%]; Cambridge Isotope Laboratories CNLM-539-H-1) and lysine (L-lysine:2HCL [13C6, 99%; 15N2, 99%]; Cambridge Isotope Laboratories CNLM-291-H-1). Neurons were harvested at various days after media switch. Protein lysates were collected in either Lysis buffer (50 mM Tris-HCl pH 7.5, 300 mM NaCl, 1% Triton-X-100, 5 mM EDTA) or SP3 buffer (50 mM Tris-HCl, pH 8.0, 50 mM NaCl, 1% SDS, 1% Triton X-100, 1% NP-40, 1% Tween 20, 1% glycerol, 1% sodium deoxycholate [w/v], 5 mM EDTA pH 8.0, 5 mM DTT, 5 kU Benzonase, and 1X complete protease inhibitor cocktail [Sigma Aldrich 5892970001]) and then underwent SP3-cleanup and a trypsin/Lys-C digest protocol. Samples were dried by speed vac. In select experiments, peptides were then treated with PNGase F (2 µL, NEB, P0704L) for 2 hours at 37 °C. Following incubation, 4 µL of 10% formic acid (FA) was added until the pH reached ~2. Samples were then dried by speed vac, reconstituted in 2% acetonitrile (ACN) with 0.1% formic acid, and analyzed by LC-MS/MS.

**LC-MS/MS analysis**

An UltiMate 3000 nano-HPLC system (Thermo Scientific) coupled with an Orbitrap Eclipse mass spectrometer (Thermo Scientific) was used to analyze proteomic cell culture samples. After samples were normalized to 0.2 µg/µL, 1 µg of digested peptide was loaded on a trap column with the loading pump set to 5 µg/min. Peptide separation was done on an ES903A nano column (75 µm × 500 mm, 2 µm C18 particle size) using an 80-min linear gradient. Mobile phases A (5% DMSO in 0.1% FA in water) and B (5% DMSO in 0.1% FA in ACN) were used. Column flow rate was set to 300 nL/min, and the column temperature was maintained at 60 °C. Data-independent acquisition mode (DIA) was used. The MS1 scan was set at a resolution of 120,000, with a standard AGC target, and the maximum injection volume was set to auto. The MS2 scans covered a precursor mass range of 400-1000 *m/z*, using an isolation window of 8 Da with 1 Da overlap, resulting in a total of 75 windows per scan cycle. Fragmentation was performed using high-energy collisional dissociation (HCD) with a normalized collision energy of 30%. A resolution of 30,000 with a scan range of 145-1450 *m/z* and the loop control set to 3 s was used for acquiring the MS2 spectra.

**Proteomic data analysis**

For proteomics from cell culture experiments, Spectronaut (version 19, Biognosys) was used with a directDIA search strategy, using factory default settings and the UniProt human proteome reference with reviewed genes (20,384 entries) as the library FASTA file. Contaminant proteins were removed using a neuron-specific contaminant library (available to download at https://github.com/HaoGroup-ProtContLib) (*93*). Trypsin/P and Lys-C were selected as the digestion enzymes, with a specific selection criterion of at most two missed cleavages per peptide. Carbamidomethylation of cysteine residues was set as a fixed modification. Methionine oxidation, N-terminal acetylation, and glycosylation were considered for variable modifications in the analysis. False-discovery rate (FDR) thresholds for precursors, proteins, and peptides were set to 0.01. For label-free proteomics data, cross-run normalization was enabled to account for systematic variations across different runs, and no imputation was performed on the dataset. For dSILAC proteomics data, a labeled workflow was used without imputation and normalization in Spectronaut. *In silico* generation of missing channels was enabled. Lys8/Arg10 was selected as the heavy channel, and b-ions were excluded during quantification for SILAC data. Peptide-level export files were used for downstream analysis. Protein/peptide half-lives were calculated as described previously (*39*, *94*, *95*) using the heavy/light peptide abundance ratios. A representative subset of peptides is graphed in fig. S1, not every modified peptide. For all half-life data for TMEM106B peptides in fig. S1 and Fig. 2, see Tables S4 and S5.

**Immunoblots**

At maturity, cells were washed with cold PBS on ice, scraped into 1.5 mL protein lo-bind Eppendorf tubes, spun at 1,000 × *g* for 10 min at 4 °C to generate a pellet, and then pellets were snap frozen in liquid nitrogen and stored at -80 °C. Cell lysates were collected using lysate buffer (50 mM Tris-HCl pH 7.5 [Life Technologies 15567027], 300 mM NaCl [Life Technologies AM9759], 1% Triton X-100 [Sigma Aldrich 93443-100ML], 5 mM EDTA [Life Technologies 15575020], cOmplete Protease Inhibitor Tablet [Sigma Aldrich 5892970001], PhosSTOP Tablet [Sigma Aldrich 4906837001], 5KU Benzonase [Sigma Aldrich E8263-5KU]). Cells were lysed for at least 1 h at 4 °C in lysis buffer, spun at maximum speed (18,213 × *g*) for 10 min at 4 °C, and the supernatant was collected and frozen at -80 °C. Lysates were mixed with 4× Laemmli buffer containing 10% beta-mercaptoethanol and loaded onto Invitrogen Tris-Glycine 4–20% gels (XP04205BOX or WXP42026BOX), run at 100–150 V for 1–2 h on ice, and transferred to a 0.2 µm nitrocellulose membrane by semi-dry transfer using a Bio-Rad TransBlot Turbo transfer system. The membrane was then stained with Ponceau S for 15–30 min, washed with TBST (1× TBS with 0.1% Tween 20), blocked in 5% milk in TBST for at least 1 h, incubated with the indicated primary antibody (Table S8) in 5% milk in TBST overnight at 4 °C, washed with TBST, incubated with the appropriate HRP-conjugated secondary antibody (1:1000–1:10,000; ), washed with TBST, and then washed with TBS before imaging using ECL-reagent (Life Technologies 34096 or 32106) for chemiluminescence. Imaging was performed on a Bio-Rad ChemiDoc Imaging system. Immunoblot for human progranulin in Fig. 4E largely followed procedures described for RIPA-soluble human brain protein extracts below, except samples were diluted with 2– SDS gel loading buffer at a 1:1 (v/v) ratio with 10% beta-mercaptoethanol, boiled at 95 °C for 5 min, and ran on 20-well 4–20% Tris-glycine gels (Novex).

For immunoblotting of patient brain-derived lysosomes, post-enrichment, samples were thawed and resuspended in a final working concentration of 1× Laemmli buffer (Bio-Rad, 161-0747) and boiled at 95 °C for 5 min. Samples were run on a Bio-Rad Precast protein gel at 200 V and transferred onto a PVDF membrane using a Bio-Rad Trans-Blot Turbo Transfer System. Membranes were blocked in 5% milk in TBST (20 mM Tris-HCl, 150 mM NaCl, 0.1% w/v Tween 20) for 1 h at room temperature. Subsequently, the membranes were incubated in primary antibodies (Table S8) for 2 h at room temperature or overnight at 4 °C. Membranes were washed 2 times for 10 min each in TBST, then incubated with secondary antibodies for 1 h at room temperature. Following this, the membranes were washed three times for 10 min each in TBST, then developed using Clarity Western ECL substrate (Bio-Rad, 1705061) and imaged on a ChemiDoc Imaging System (Bio-Rad).

**Cell line editing and genetic engineering**

iPSCs were maintained in StemFlex (Life Technologies A3349401) or E8. To make knockout cell lines or targeted edits, Lonza P3 primary cell solution (Life Technologies V4XP-3032) was used for nucleofection of a sgRNA, ssODN, and Cas9 HiFi V3 protein (IDT #1081061). A Lonza 4D-nucleofector (AAF-1003B and AAF-1003X) was used with the “CA-137” protocol. For the A9D mutations in *GRN*, the sgRNA sequence used was: 5'-AGCTGGGTGGCCTTAACAGC/AGG and the ssODN used for templated repair was: 5'-CCAGGCAGCAGGCCACAGGGCAGAACTGACCATCTGGGCACCGCGTTCCAGCCACCAGCCCTGCTGTTAAGTCCACCCAGCTCACCAGGGTCCACATGGT. The KOLF2.1J cell line is heterozygous at the T/S185 site, so for each intended homozygous cell line, an edit was made. To introduce the T185 homozygous edit, the sgRNA sequence used was: 5'-TTAAACAACATAAGCATTAT/TGG and the ssODN used was: 5'-TCATTTAACTCTTACAAAGTATTTCATGATTGATTCTTACTTGTTTCATATCAAGTGGACCAATAATGGTTATGTTGTTTAAGCGTGCCTTTCCAATAAC. To introduce the S185 homozygous edit, the sgRNA sequence used was: 5'-TTAAACAACATAACCATTAT/TGG and the ssODN used was: 5'-TCATTTAACTCTTACAAAGTATTTCATGATTGATTCTTACTTGTTTCATATCAAGTGGACCAATAATGCTTATGTTGTTTAAGCGTGCCTTTCCAATAAC. After nucleofection, cells were grown at 32 °C for 3 days before returning to normal growth at 37°C. Alt-R HDR Enhancer V2 (IDT #10007921) was present at a 1 uM final concentration in the media for 24h post-nucleofection. Cells then underwent single cell cloning and colony picking. Colonies were screened for the desired gene edit with target site PCR and Sanger sequencing. For the GRN-knockout lines, the cells were nucleofected using two sgRNAs (5’-GCAAAGTACCAAGGAACGTC/TGG and 5’-TAAGGCCTTCCCTGTCAGAA/GGG) and an ssODN (CTGAGTGACCCTAGAATCAAGGGTGGCGTGGGCTTAAGCAGTTGCCAGA*DH*AAGGGGGTTGTGGCAAAAGCCACATTACAAGCTGCCATCCCCTCCCCGT). After single cell cloning and colony picking, the colonies with the lowest-GRN expression were selected by immunofluorescence against progranulin and LAMP1. For the TMEM106B-knockout line, KOLF2.1J iPSCs were generated using two sgRNAs (5’-CCAGGTCATAGGTGCATAGG/TGG and 5’-AGAGTACAGTTAAAAGTATG/TGG) and an ssODN (gagtttaaccgtttcaaccaattgccattaggaaatctttaaatccaccVBTGTGGACCTGCAGTTCTTGTAACTCTCCACTCTGTGTTAATGATATATT). After single cell cloning and colony picking, the colonies with TMEM106B-loss were selected by immunofluorescence staining TMEM106B protein and knockout was confirmed via immunoblot.

**PLD3-TMEM106B^FC^ plasmids**

The PLD3-TMEM106B^FC^ plasmid was cloned by Twist Biosciences using a previously published plasmid backbone to reduce gene silencing in differentiated cells (*96*) downstream of the CAG promoter. The PLD3-control plasmid encodes the PLD3(1–73)-GSGSG amino acid sequence, and the PLD3-TMEM106B plasmid encodes PLD3(1–73)-GSGSG-TMEM106B(120–274). Plasmids were transformed into competent cells. Single colonies were picked and liquid culture was grown for Maxi Prep. Whole plasmid sequencing by Quintara Bio was used to verify plasmid identity. Coding sequences for the genes are listed in Table S9. For integration into iPSCs, 2 µg of plasmid DNA was mixed with 1 µg of piggybac transposase and transfected into iPSCs using Lipofectamine Stem Reagent (Thermo Fisher STEM00015) following manufacturer protocols. Cells underwent selection with puromycin up to 10 µg/mL until culture was pure based on fluorescence from the plasmid (usually ~5–7 days of selection).

**Quantitative PCR (qPCR)**

At maturity of differentiation (typically Day 14), cells were lysed in 300 μL of TRI reagent (Zymo Research R2050-1-200). RNA was then collected using a Zymogen Direct-zol RNA kit (Cat no. R2050) following manufacturer instructions, including the optional DNase-treatment step. For mice, frozen brain tissue was homogenized in a volume (µL) of 1× Tris-EDTA buffer pH 8.0 totaling the mass of the tissue in mg × 5 (e.g., 500 µL for 100 mg of tissue); 90 µL were mixed with 270 µL Trizol LS (Invitrogen #10296010) and RNA was subsequently extracted using a Zymogen Direct-zol Miniprep Plus RNA kit (Cat no. R2073-A) following manufacturer instructions as above, including the optional DNase-treatment step. RNA concentrations were measured via Nanodrop (ThermoFisher), 500 ng of RNA per sample was then transcribed into complementary DNA (cDNA) using the High-Capacity cDNA transcription kit (Applied Biosystems), and cDNA samples were then diluted 1:20 in nuclease-free water. Quantitative real-time PCR (qRT-PCR) was subsequently performed using 2 μL of diluted cDNA per sample, with 3 μL of SYBR GreenER qPCR SuperMix (Invitrogen) added. Samples were run in triplicate on the QuantStudio™ 7 Flex Real-Time PCR System (Applied Biosystems). ΔΔCt was calculated with *GAPDH* and *RPLP0* as housekeeper gene controls. The following primers were used for both human iNeurons and mice: *PLD3-TMEM106B* forward 5’-GCGACTTGCATCTCTTTGGG; *PLD3-TMEM106B* reverse 5’-ACACCAATGTATTTCACGTCGAT. For human iNeurons, *GAPDH* forward: 5′-GTTCGACAGTCAGCCGCATC, *GAPDH* reverse: 5′-GGAATTTGCCATGGGTGGA; and *RPLP0* forward: 5′-TCTACAACCCTGAAGTGCTTGAT; *RPLP0* reverse: 5′-CAATCTGCAGACAGACACTGG. For mouse samples: *GAPDH* forward: 5’-CATGGCCTTCCGTGTTCCTA; *GAPDH* reverse 5’-CCTGCTTCACCACCTTCTTGAT; *RPLP0* forward 5’-ACTGGTCTAGGACCCGAGAAG; *RPLP0* reverse 5’-ctcccaccttgtctccagtc.

**Progranulin-rescue experiments**

Recombinant progranulin was produced as in Gass et al., 2012 (*97*). Briefly, a pCDNA3/PGRN-6His plasmid was transfected into HEK293 cells and selected for stable, high expression of recombinant PGRN secreted into the media; rhPGRN was purified from this line using standard Ni-NTA (Qiagen) affinity purification using standard manufacturer protocols.

KOLF2.1J iNeurons were differentiated following their normal differentiation protocol. Following induction (Day 4 onward for KOLF), the media on the rhPGRN-rescue conditions were supplemented with 30 ng/µL rhPGRN protein. Cells were maintained in this media until collection, with half media changes approximately 2 times per week.

**Brain protein and peptide QTL analyses**

Protein and peptide QTL mapping in postmortem human brain was performed using proteomic profiles from the dorsolateral prefrontal cortex (dPFC) on either 1,369 individuals for protein QTL mapping or 900 individuals for peptide QTL mapping (who were a subset of the 1,369 donors).

For both analyses, genotyping was either generated from array-based genotyping or whole genome sequencing (WGS), as described previously (*98*). For array-based genotyping, imputation was performed using TOPMed using recommended quality control steps, and SNPs with imputation quality R2 > 0.3 were retained (*34*). For WGS, samples were assessed for completeness, coverage, transition:transversion ratio, and silent:replacement ratio with outliers beyond 6 standard deviations (SD) being dropped. Samples were clustered with Phase 3 1000 Genomes data as reference using individual genotyping, and multidimensional scaling analysis was performed to identify group outliers beyond 3 SD. We performed further quality control in each genetic ancestry separately by excluding variants with any of the following criteria: genotype missing rate > 5%, minor allele frequency (MAF) < 5%, non-biallelic variants, and Hardy–Weinberg equilibrium *p* < 10 × 10^-07^. Related individuals were identified using KING (version 2.2.2) (*99*) and individuals who were second-degree or closer relatives were randomly removed. Finally, EIGENSTRAT (version 6.1.4) (*100*) was used to identify any remaining population outliers.

Brain proteomes were profiled using tandem mass tag mass spectrometry as described in detail in previous studies (*34*, *101*). For each round of sequencing, samples were randomized across batches to evenly distribute sex, age, diagnosis, and race/ethnicity per batch and labeled with isobaric tandem mass tag peptide labels. Each proteomic batch included a global internal standard that was created by pooling protein across samples within the batch. High-pH fractionation was performed, and samples were analyzed using liquid chromatography coupled to tandem mass spectrometry. Raw files were searched using Fragpipe (version 19.0) (*101*), MSFragger (version 3.5) (*101*), and the human proteome database Swiss-Prot (*102*) containing 20,402 sequences (downloaded 2/11/2019). Subsequently, we used Post-MSFragger (version 3.6) and Percolator (*103*) for PSM validation and Philosopher (version 4.6.0) for protein inference using ProteinProphet and filtering by false discovery rate. The database search yielded a total of 11,748 protein groups from the dorsolateral prefrontal cortex. Proteome quality control began by normalizing abundance levels using sample-specific protein counts and log2 transformation. Principal component analysis was used to identify samples > 4 SD from the mean after two iterations. Regression was used to estimate proteomic batch, postmortem interval, age, and clinical diagnosis, and those estimates were regressed out of the final protein abundance matrix. To combine proteomic profiles across rounds of sequencing (~80 batches) protein abundance was z-scale-normalized. To identify potential hidden confounding variables, surrogate variable analysis was performed (SVA v.3.20.0) (*104*), and significant surrogate variables were regressed from the proteomic profiles. Finally, the regressed proteomic profiles were inverse rank-normalized and proteins with a minimum of 200 samples per protein were retained.

Peptides were extracted from a subset of 900 individuals for TMEM106B peptide analysis, and TAMPOR (*105*) was used to perform normalization and quality control analogous to the batch quality control used for the protein quantification quality control. There were 11 TMEM106B peptides measured, with 6 of 11 passing the completeness threshold of > 50% for analysis. Notably, rs3173615 p.S185T is not covered by either C-terminal peptides studied, SAYVSYDVQK (130–139) and YQYVDCGR (248–255).

For protein and peptide QTL mapping, SNPs with MAF ≥ 0.05 were tested for their association with normalized protein expression as the outcome, adjusting for sex, 10 genetic principal components, and the source of genetic data (WGS vs. genotyping array) using PLINK2 (*106*).

**Domain QTL analyses**

Genomic and proteomic data came from the Religious Orders Study – Memory and Aging Project (ROSMAP) (*37*). All participants enrolled without known dementia and agreed to annual evaluation and brain donation. Both studies were approved by the Institutional Review Board of Rush University Medical Center, and all participants signed informed and repository consents and an Anatomic Gift Act. Genomic and proteomic data was generated as previously reported (*107*, *108*). For the domain-pQTL analysis, the ROSMAP mass spectrometry dataset was searched with a new FASTA file that removed native TMEM106B protein and added proteins based on either the TMEM106B N-terminus (TMEM106B residues 1–120) or the TMEM106B C-terminus (TMEM106B residues 119–274) to the search. The overlapping residues between the two domains (residues 119–120) were used to ensure mapping of all possible tryptic peptides in the region. With this, an abundance value was determined for each domain of TMEM106B as if they were independent proteins. Subseqeunt pQTL analyses were conducted via one-way ANOVA with Tukey post-hoc analysis using a publicly available parANOVA R script (https://github.com/edammer/parANOVA ).

**Human samples**

Postmortem tissue from individuals with FTLD-TDP was provided by the Mayo Clinic Brain Bank, and neuropathological diagnosis was determined by a single neuropathologist (D.W.D.). Written informed consent was obtained from all donors prior to entry, or from the next-of-kin if the donor was unable to provide written consent. All protocols were approved by the Mayo Clinic Institutional Review Board.

**Human brain protein extraction**

Sarkosyl-insoluble and RIPA-soluble fractions of FTLD-TDP frontal cortex were generated as described previously (*31*). Briefly, human postmortem frontal cortex tissue was homogenized in 5 volumes (w/v) of cold buffer consisting of 10 mM Tris-HCl (pH 7.4), 80 mM NaCl, 1 mM MgCl_2_, 1 mM EGTA, 0.1 mM EDTA, 1 mM PMSF, 1 mM fithiothreitol, and a protease and phosphatase inhibitor cocktail. Following homogenization, 400 µL of homogenate (corresponding to 80 mg brain tissue) was ultracentrifuged at 150,000 × *g* for 40 min at 4 °C in a TLA110 rotor. The resulting pellet was resuspended in one volume of cold buffer (containing 10 mM Tris pH 7.4, 0.85 M NaCl, 10% sucrose, and 1 mM EGTA) and subsequently centrifuged at 14,000 × *g* for 10 min at 16 °C. The supernatant was then incubated with sarkosyl at a final concentration of 1% for 1 h at room temperature with continuous shaking. Samples were then spun at 150,000 × *g* for 40 minutes at 4 °C in a TLA110 rotor. The resulting pellet (sarkosyl-insoluble P3 fraction) was resuspended in 50 µL TE buffer (10 mM Tris-HCl, 1 mM EDTA) and stored at -80 °C until use. To generate RIPA-soluble extracts, tissues were homogenized in 5 volumes (w/v) of ice-cold RIPA buffer (25 mM Tris-HCl pH 7.5, 150 mM NaCl, 1% NP-40, 1% sodium deoxycholate, 0.1% SDS, with protease and phosphatase inhibitors). Homogenates were then sonicated (1 s on, 1 s off for 10 s) and centrifuged at 100,000 × *g* for 30 min at 4 °C. The resulting supernatants were collected as the RIPA-soluble fraction.

**Immunoblotting of human brain protein extracts**

The sarkosyl-insoluble fraction (10 µL) was diluted with 2× SDS gel loading buffer at a 1:1 (v/v) ratio, then heated at 95 °C for 5 min. Samples were loaded into 20-well 4–20% Tris-glycine gels (Novex) and subsequently transferred to PVDF membranes. RIPA soluble extracts were diluted 1:1 (v/v) in 2× SDS gel loading buffer while being kept cold on ice. Samples were then run on 10% Tris-glycine gels with ice surrounding the tank to maintain cold conditions. All membranes were blocked with 5% nonfat dry milk in TBS + 0.1% Triton X (TBST) for 1 h at room temperature and then incubated with primary antibody overnight at 4 °C with gentle rocking. The next day, membranes were washed in TBST 3× for 10 min at room temperature and incubated with donkey anti-rabbit or anti-mouse IgG antibodies conjugated to horseradish peroxidase (1:5000; Jackson ImmunoResearch) for 1 h at room temperature. Protein expression was visualized using enhanced chemiluminescence and developed on an Amersham Imagequant 800. Blot quantifications were performed in ImageJ.

**Mice**

All procedures involving rodents were performed in accordance with the National Institutes of Health Guide for Care and Use of Experimental Animals and approved by the Mayo Clinic Institutional Animal Care and Use Committee (IACUC). Mice were maintained on a 12-hour light/dark cycle in standard housing at Mayo Clinic Jacksonville with access to chow and water ad libitum. C57BL/6J mice were originally obtained from The Jackson Laboratory (strain #000664).

**AAV production and neonatal mouse injections**

Recombinant adeno-associated virus (AAV) 9 was produced as previously described (*109*) and purified using standard methods. Briefly, AAV vectors encoding PLD3-GS-TMEM106B^FC^ or PLD3-control were co-transfected with the cis-plasmid pCap9 (a gift from Dr. Todd Golde, Emory University) into HEK 293T/17 cells (ATCC #CRL-11268; RRID: CVCL_1926). Cells were harvested 72 h after transfection, treated with 50 units/mL Benzonase endonuclease (Sigma-Aldrich), and lysed by freeze-thaw. The virus was isolated using a discontinuous iodixanol gradient and buffer-exchanged into PBS using an Amicon Ultra-15 centrifugation filter concentrator (Millipore #UFC910024). The genomic titer of each virus was determined by qPCR using the Quant Studio 7 Flex Real-Time PCR System (Applied Biosystems), SYBR Green PCR Master Mix (Applied Biosystems #4309155), and primers specific to the woodchuck hepatitis virus post-transcriptional regulatory element (WPRE). Samples were compared against a standard curve of supercoiled plasmid and diluted to their final concentrations in PBS. Intracerebroventricular injections of virus into post-natal day zero (P0) C57BL/6J wild-type pups were performed as previously described (*109*). Pups were cryoanesthetized on ice until they exhibited no movement, and 2 µl AAV (2.5 × 10^13^ genomes/µl) solution was injected into each lateral ventricle using a 32-gauge needle (Hamilton Company product #7803–04, 0.5 in custom length, point style 4, 12°) fitted to a 10 μL syringe (Hamilton Company). The needle was inserted at a 30° angle from the surface of the head at a point approximately two-fifths of the distance between the lambda suture and the eye and held at a depth of approximately 2 mm when injecting. Pups were then allowed to recover on a heating pad before being returned to their home cages.

**Mouse brain protein extraction**

Mouse hemibrain was harvested at 13 months of age and homogenized in 5 volumes (w/v) of cold 1x Tris-EDTA buffer pH 8 before being aliquoted for subsequent experiments. For sarkosyl extraction, 150 µL of homogenate was combined with buffer consisting of 20 mM Tris-HCl (pH 7.4), 160 mM NaCl, 2 mM MgCl_2_, 2 mM EGTA, 0.2 mM EDTA, 2 mM PMSF, 2 mM dithiothreitol, and a protease and phosphatase inhibitor cocktail. Homogenate was subsequently ultracentrifuged at 150,000 × *g* for 40 min at 4 °C in a TLA110 rotor. The resulting pellet was resuspended in one volume of cold buffer (containing 10 mM Tris [pH 7.4], 0.85 M NaCl, 10% sucrose, and 1 mM EGTA) and subsequently centrifuged at 14,000 × *g* 10 min at 16 °C. The supernatant was then incubated with sarkosyl at a final concentration of 1% for 1 h at room temperature with continuous shaking. Samples were then spun at 150,000 × *g* for 40 minutes at 4 °C in a TLA110 rotor. The resulting pellet (sarkosyl-insoluble P3 fraction) was resuspended in 50 µL TE buffer (10 mM Tris-HCl, 1 mM EDTA) and stored at -80 °C until use. To generate RIPA-soluble extracts, Tris-EDTA hemibrain homogenates were mixed with 2× ice-cold RIPA buffer (50 mM Tris-HCl pH 7.5, 300 mM NaCl, 2% NP-40, 2% sodium deoxycholate, 0.2% SDS, with protease and phosphatase inhibitors). Homogenates were then sonicated (1 s on, 1 s off for 10 sec) and centrifuged at 100,000 × *g* for 30 min at 4 °C. The resulting supernatants were collected as the RIPA-soluble fraction.

**CSF neurofilament light chain (NfL) measurement**

CSF was collected from anesthetized mice via the cisterna magna immediately prior to brain harvest and frozen on dry ice. Samples were measured in a blinded manner using the NF-Light digital immunoassay (Quanterix, Cat#103,186). Immediately prior to preparation, samples were thawed on ice and diluted 1:25 in Quanterix sample diluent (provided with assay kit) before transferring samples to 96-well plates. Samples were diluted 1:4 by the instrument and tested in duplicate. Each run included eight calibrators and two quality control samples provided with the kits. Concentrations were interpolated from the standard curve using a 4-parameter logistic curve fit (1/y2 weighted).

**Immunohistochemistry**

Paraffin-embedded sagittal mouse tissue sections were cut at 5 μm, mounted on positively charged glass slides, and dried overnight in a 60 °C oven. Prior to staining, mouse and human sections were deparaffinized in xylene three times, 5 min each, and rehydrated in progressively diluted ethanol (100%, 100%, 95% for 3 min each). Following three 5-min washes in deionized water, antigen retrieval was performed by steaming in 1× Tris-EDTA (TE) buffer (pH 9) for 20 min. Slides were allowed to cool on the bench and were subsequently incubated with Dako Peroxidase block (DAKO S2001) at room temperature for 5 min to quench endogenous peroxidase activity. IBA1 and GFAP immunostaining of mouse tissues were subsequently performed using the Thermo Scientific Lab Vision Autostainer 480S (Kalamazoo, MI) and the DAKO EnVision + HRP system. For TMEM106B, immunostaining sections were stained manually. Following the 3% hydrogen peroxide incubation, sections were washed in 1× TBST three times for 5 min and blocked at room temperature for 20 min in Dako Serum-Free Protein Block (X090930-2). Primary antibody was then diluted in Dako antibody diluent (S302283-2) and incubated on the tissues for 45 min at room temperature. Sections were then washed three times for 10 min in 1× TBST and incubated for 30 min in Dako Envision+ System HRP-Labelled Polymer (# K4000; anti-rabbit) at room temperature for 30 min. Following another round of washes in TBST, sections were incubated in Dako Envision+ Dual Link System-HRP (DAB+; #K4065) for 5 min, after which the sections were rinsed in dH_2_O before dehydration and coverslipping. For pTDP-43 immunostaining, sections were deparaffinized as above. Antigen retrieval was then performed by steaming sections in 1 mM sodium citrate buffer, pH 6.0 + 0.05% Tween 20 buffer for 30 min. Sections were then incubated with Dako Peroxidase Block (DAKO, S2001) followed by blocking in 2% normal goat serum in PBS for 1 h at room temperature. Slides were then incubated with rabbit anti-TDP-43 pS409/410 (1:500, ab 3655, developed in-house) overnight at 4 °C in a humidified chamber. After 310-min washes in 1× PBS the next morning, sections were incubated with an HRP-conjugated anti-rabbit secondary antibody (1:200, Jackson ImmunoResearch) at room temperature for 2 h. Peroxidase labeling was visualized with the chromogen solution 3,3′-diaminobenzidine (DAB-Plus, DAKO) and the VECTASTAIN Elite-ABC kit (Vector Laboratories) per manufacturer’s instructions. Following chromogen development, all slides (regardless of stain) were briefly rinsed in deionized water and subsequently counterstained with hematoxylin, dehydrated, cover-slipped, and scanned with the Leica Aperio AT2 Slide Scanner (Aperio, Vista, CA).

**Immunofluorescence and confocal microscopy**

Paraffin-embedded sagittal mouse tissue sections (5 μm) were deparaffinized, rehydrated, and washed in deionized water. Following antigen retrieval in 1× TE buffer (pH 9) or 1× citrate buffer (pH 6), slides were cooled and blocked with Dako All Purpose Blocker for 1 h. Slides were then stained in one of two ways: either with dye-conjugated primary antibodies or with primary antibodies from different species combined for overnight staining. For the former method, Biotium CF dyes were conjugated to antibodies via Biotium Mix ‘N Stain Kits (#92446) per manufacturer’s instructions. Slides were then incubated overnight with antibody in a humidified chamber at 4 °C, followed by washing in 1× TBST three times for 10 min each the next day. For routine staining, slides were also incubated with primary antibody overnight in a humidified chamber at 4 °C. The next day, slides were washed in 1× TBST three times for 10 min each, then incubated with a fluorescent dye-conjugated secondary antibody for 2 h in the dark at room temperature. After 3 10-min washes in TBST, slides were incubated for 10 min at room temperature in Hoechst dye (Invitrogen H3570) diluted in 1× PBS at 1:1000. Slides were again washed three times for10 min each in 1× TBST and were coverslipped with Fluoromount G. Images were obtained on a Zeiss LSM 980 laser scanning confocal microscope using a 40× objective.

**Image analysis**

To obtain the CD68/IBA1 ratio, IBA1+ cells were summed across 10–12 cortical fields in FIJI by researchers blinded to CD68 status. Of those IBA1+ cells, the number of CD68+ cells was counted and the ratio of CD68/IBA1 was plotted as a percentage. P62 inclusions were quantified across 6 cortical fields per animal in FIJI. Specifically, a 50-pixel rolling ball radius was applied for background subtraction of each image, and inclusions that met the intensity threshold and minimum size threshold of 0.5 µm^2^ were converted to an overlay mask. The same intensity threshold was applied to each image. The analyze particles feature was then applied to generate the total number of inclusions/objects. To calculate inclusions/mm^2^ for an individual animal, the total inclusion number was summed across all images and divided by the total area analyzed across all images. Immunohistochemistry (IHC) images (representative images in fig. S4I) were used for manual counting of pTDP-43 inclusions in Aperio Imagescope (Leica) by an investigator blinded to the experimental group. For IBA1 and GFAP IHC, a positive-pixel detection algorithm in Aperio Imagescope software was batch-run on all images to quantify the percentage of cortical area positive for the marker of interest.

***In situ* cryo-correlative light and electron microscopy**

*Grid preparation*

Gold grids, 200 mesh, with holey R1/4 SiO_2_ film (Quantifoil Micro Tools GmbH Q250AR-14S) were glow-discharged for 30 s at 5 mA using an EasiGlow system (Pelco). To achieve cell growth in the center of grid squares, grids were coated with poly-L-lysine (Sigma-Aldrich P6282) and mPEG-SVA (Laysan Bio MPEG-SVA-5000). Coated grids were then photosensitized by adding PLPP gel (Alvéole B004) and photo-micropatterned using a DMi8 microscope (Leica Microsystems) equipped with an Alvéole PRIMO 2 to generate 40 µm-wide circular patterns at the centers of grid squares. Grids were subsequently coated with 100 µg/mL PLO overnight, followed by four washes with sterile water and air-drying for at least 1 h. Next, grids were treated with 10 µg/mL laminin in a 35 mm glass-bottom dish (MatTek Life Sciences) for at least 1 h.

*Sample preparation*

Neural progenitor cells (NPCs; 1×10^5^) in TeSR-E8, doxycycline, and ROCK inhibitor were grown in 35 mm glass-bottom dishes containing micropatterned Au SiO_2_ R1/4 200 mesh grids coated with PLO and laminin as described above. NPCs were differentiated to DIV9 iNeurons before plunge-freezing in liquid ethane. After differentiation, the grids were transferred to a new 35 mm dish containing 50 nM LysoTracker Green (504/511 nm) DND-26 (Thermo Fisher Scientific L7526) in warm B27 media for 3 min at 37 °C to stain lysosomes and wash away excess/free cell debris. Blue (365/415 nm) FluoSpheres (Invitrogen F8824) were sonicated in a water bath sonicator for 10 s and then diluted 1:10 in B27 media. Grids were then incubated with 2 µL of diluted FluoSpheres, back-side blotted with calcium-free filter paper (Subangstrom LFP01), and plunge-frozen into liquid ethane at -185°C using a GP2 (Leica Microsystems) automatic grid plunger with the following specifications: *blot time*: 10 s, *temperature*: 37 °C, and *humidity*: 80%. The plunge-frozen grids were clipped with C-Clips (Thermo Fisher Scientific 1036171) into AutoGrids (Thermo Fisher Scientific 1205101) and stored in liquid nitrogen until further use.

*Cryo-fluorescence microscopy*

The quality of iNeurons and grids was assessed in widefield mode on a Stellaris-5 cryo-confocal laser-scanning microscope equipped with a cryo-stage and a 50× NA 0.9 objective (Leica Microsystems). Z-stacks of desired regions were acquired in confocal-scan mode with a 405 nm laser line, a 577 nm laser line, a 0.3 µm spacing spanning 18 µm of the focus plane, and a *x/y* pixel size of 72.11 nm. Subsequent correlations between cryo-confocal fluorescence, cryo-focused ion beam (cryo-FIB), cryo-scanning electron microscopy (SEM), and cryo-transmission electron microscopy (cryo-TEM) images were performed with 3D Correlation Toolbox (3DCT) to guide cryo-FIB milling and tomography tilt-series positioning (*110*).

*Cryo-FIB milling*

Clipped Autogrids were loaded onto the stage of an Aquilos 2 FIB-SEM (Thermo Fisher Scientific) maintained at a temperature of ≤ -185 °C. Grids were coated with a layer of organometallic platinum using a gas injection system for 30 s and then sputter-coated with inorganic platinum to form a protective/conductive surface for cryo-FIB milling. Initial correlation between the grid overview images from wide-field fluorescence and scanning electron microscopy (SEM) was performed using Maps 3.20 (Thermo Fisher Scientific) to identify neurons for milling and to provide a rough estimation of lamella positioning. Next, 3D CT was used to correlate the z-stack cryo-confocal fluorescence microscopy and cryo-FIB data, aided by FluoSpheres, and precisely define the sites of lamellae. Initial cuts with the cryo-FIB to make tension-relief trenches were milled with a beam current of 1 nA (*111*). The neurons were milled at 1 nA, 500 pA, and 300 pA sequentially to achieve a ‘rough’ lamella thickness of 5, 3, and 1 µm, respectively. The milling angle varied from 5° to 16° and was dependent on the thickness of the neuron. Milling progress and lamella integrity were monitored via SEM imaging. Upon completion of all rough lamella on a grid, each lamella was revisited and polished to a ‘fine’ lamella of ~200 nm in thickness, with the ion beam current of 50 pA.

*Tilt-series data acquisition*

Grids containing iNeuron lamellae were imaged on a 300 kV Titan Krios G4i transmission electron microscope (Thermo Fisher Scientific) equipped with a K3 direct electron detector and an imaging filter (Gatan Inc.) operated in counting mode. Dose-symmetric tilt series were collected, at correlated sites, from -60° to +60°, starting at the milling angle (5–16°) at a 3° interval, using SerialEM software (*112*, *113*). The magnification was set to a pixel size of 2.654 Å/pixel, and the total dose per tilt series was 160 e^–^/Å^2^. Data were acquired using a defocus range of -3 µm to -5 µm and with a 20 eV energy filter slit width.

**Lysosomes-enrichment from iNeurons, mouse brain and patient brain tissue**

*Pre-lysis*

At maturity of differentiation (either Day 18 or Day 42), iNeurons were scraped and pelleted at 1000 × *g* for 10 min at 4 °C and snap frozen with liquid nitrogen. The cell pellet was resuspended in ice-cold lysis buffer (0.25 M sucrose [Sigma-Aldrich 84100[, 1 mM EDTA, 10 mM HEPES pH 7.4 [Sigma-Aldrich H4034], 1 mM PMSF [Sigma-Aldrich P7626], 1× Protease Inhibitor Cocktail [APExBIO K1010]). For mouse and postmortem human brain tissue, we measured and coarsely chopped 0.6 grams of frozen prefrontal cortex tissue into 1–-5 mm^2^ pieces and added them to 7 mL of warm Hibernate A media (Gibco A1247501) containing 20 U/mL papain (Worthington LK003178) for 20 min. Papain digestion was terminated by the addition of ice-cold cocktail of protease inhibitors containing 5 μg/mL leupeptin (Sigma-Aldrich L2884), 5 μg/mL antipain dihydrochloride (Thermo Fisher Scientific AAJ63680LB0), 5 μg/mL pepstatin A (Sigma-Aldrich P4265), 1 mM PMSF, and 1 μM E64 (Tocris Bioscience 52-081-0) in 13 mL Hibernate A media. Digested tissue was spun at 500 × *g* for 10 min at 4˚C and resuspended in ice-cold lysis buffer.

*Cell lysis and fractionation*

All steps of gradient fractionation were performed with ice-cold buffers and centrifuges pre-cooled to 4 °C, using a slightly modified autophagic isolation protocol previously reported (*114*). The resuspended cell/tissue pellet was lysed using a pre-chilled Dounce homogenizer and a tight-fitting pestle (Kimble “B”) for 20 strokes on ice before centrifugation on a benchtop fixed-angle centrifuge. The lysate was centrifuged at 1000 × *g* for 5 min to pellet nuclei and unbroken cells. The remaining supernatant was collected for centrifugation, and the remaining pellet fraction was resuspended and dounced again to ensure robust cell lysis. After nuclei were cleared a second time, the supernatant fractions were combined and centrifuged at 5000 × *g* for 10 min to separate the heavy mitochondrial fraction. The resulting supernatant was collected and spun at 21,000 × *g* for 20 min to pellet the membrane fraction (which contains autophagosomes and lysosomes). The supernatant was discarded, and the pellet fraction was resuspended in 750 μL of 51% Nycodenz (Accurate Chemical, AN1002423) in gradient dilution buffer (0.125 M sucrose, 1 mM EDTA, 10 mM HEPES pH 7.4, with fresh protease inhibitors) on ice. The resuspended fraction was overlaid with equal volumes of 40%, 30%, 25%, 20%, 15%, 10%, 5%, and 0% Nycodenz (all prepared in gradient dilution buffer on ice; 10% and 5% Nycodenz layers were only included for mice and human brain tissue) in an ultracentrifuge tube (Beckman Coulter). This was centrifuged for 3 h at 105,000 × *g* in a Beckman SW 40 rotor at 4 °C. Following centrifugation, the fractions 1–-6 were collected. iNeuron fraction 2 (at the 0% and 15% Nycodenz interfaces) and mouse/patient tissue material fraction 4 (at the 10% and 15% Nycodenz interfaces) were used for downstream cryo-ET and immunogold labeling analyses. All gradient fractions, either from iNeurons, mouse, or patient brain tissue, were stored at -80 °C for western blot analysis.

**Preparation of sarkosyl-insoluble fractions**

For sarkosyl-insoluble material from iNeurons (Fig. 2G), iNeurons were grown on 10-cm dishes until Day 42. On Day 42, cells were washed with PBS and collected in protein lo-bind eppendorf tubes by spinning at 1000 × *g* for 10 min at 4 °C. The cell pellet was flash frozen in liquid nitrogen and kept at –80 °C until further processing. Sarkosyl extraction was performed as described below.

Lysosome fractions derived from iNeurons (fraction 2), mouse brain (fraction 4), and patient brain tissue (fraction 4) were used for sarkosyl-extraction. Briefly, samples were resuspended in 1 mL of extraction buffer (10 mM Tris-HCl, 0.8 M NaCl, 10% sucrose, 1 mM EGTA at pH 7.5) and then sonicated twice (Virsonic 100 probe sonicator) at a power setting of 2 for 10 s on, 50 s off. A 25% sarkosyl solution was added to the homogenates to create a final concentration of 2% sarkosyl. Homogenates were incubated for 1 h at 37 °C with orbital shaking (INCU-SHAKER™ 10LR) at 250 rpm. Next, the homogenates were transferred to thick-walled centrifuge tubes (Beckman Coulter, NC9495303) and spun at 27,000 × *g* for 10 min using the TLA120.2 Rotor (Beckman). The supernatant was transferred to a new centrifuge tube and spun at 166,000 × *g* for 20 min with a TLA120.2 rotor (Beckman). Finally, the resulting pellets were resuspended in 30–50 μL of 20 mM Tris-HCl at pH 7.4 and 150 mM NaCl by sonication (Virsonic 100 probe sonicator) for 30 s at 100% amplitude. Additional sonication, if necessary, was performed with a water bath sonicator (Branson 2800) for 30 s. The sarkosyl-insoluble fraction was then analyzed with immunogold labeling.

**Immunogold negative-stain electron microscopy**

Carbon-coated 300-mesh copper grids (Electron Microscopy Sciences CF300-CU-50) were glow-discharged for 30 s at 5 mA (Pelco). Three µL of the sarkosyl-insoluble fraction was applied to the grids, incubated for 1 min, and then blotted using Whatman 1 filter paper (Cytiva, 1001090). Blocking buffer (PBS, pH 7.4, 0.1% w/v cold water fish skin gelatin [Aurion, 900.033]) was applied to the grids and incubated for 10 min, then blotted with filter paper. Subsequently, the primary antibody (Table S8), diluted 1:20 in blocking buffer, was applied to the grids and incubated for 1 h. Following blotting, grids were washed five times with a blocking buffer, and excess solution was blotted off between each wash. Twelve nm Colloidal Gold AffiniPure Goat Anti-Rabbit IgG (Jackson ImmunoResearch Laboratories 111-205-144) diluted 1:4 in blocking buffer was applied to the grids and incubated for 1 h. Grids were washed five times with Milli-Q water, and excess liquid was blotted off between washes. Finally, grids were stained with 3 µL of 0.75% uranyl formate (Electron Microscopy Sciences 16984-59-1) for 1 min and washed twice with Milli-Q water; the excess solution was removed. The grids were imaged using a Morgagni transmission electron microscope (Thermo Fisher Scientific) at an acceleration voltage of 100 kV. The images were acquired using a Gatan Orius SC200 CCD camera at a resolution of 2.1 Å/pixel with Digital Micrograph software (Gatan Inc).

**Cryo-ET of iNeurons, mouse brain, and patient brain derived lysosomes**

Copper R2/1 300 mesh grids (Quantifoil GmBH Q310CR1) were glow-discharged at 5mA for 30 s (Pelco), and 3µL of the lysosome-enriched fraction was added before plunge-freezing into liquid ethane using the Vitrobot Mark IV (Thermo Fisher Scientific) with the following specifications: *blot time: 3 s, blot force: 0, temperature: 4˚C, humidity: 90%*. The plunge-frozen grids were clipped with C-clips into AutoGrids (Thermo Fisher Scientific 1036173) and stored in liquid nitrogen until further use. Tilt series data was collected on a 300 kV Titan Krios G4i transmission electron microscope (Thermo Fisher Scientific) equipped with a K3 direct electron detector and an imaging filter (Gatan Inc.) operated in counting mode. Dose-symmetric tilt-series were collected from -51° to +51°, starting at 0° at a 3° interval as described above. The magnification was set to a pixel size of 2.12 Å/pixel, and the total dose per tilt series was 120 e^–^/Å^2^. Data were acquired using a defocus range of -3 µm to -5 µm and with a 20 eV energy filter slit width.

**Tilt series alignment and processing**

A custom script implementing Warp (2.0.0), IMOD, and Isonet2 was used to automatically align tilt series, and to reconstruct, deconvolve, and denoise tomograms (*115*–*117*). Briefly, preprocessing steps, including frame gain normalization, motion correction, and contrast transfer function (CTF) estimation, were carried out using WarpTools. Tilt series alignment was performed using patch tracking in IMOD, discarding poorly aligned patches until satisfactory alignment scores (usually a residue error mean < 0.65) were obtained. Tomograms were reconstructed in Fourier space at a pixel size of 8.48 (4× binning) and filtered using Wiener-like deconvolution in WarpTools. Even-odd tomograms were generated separately by splitting frames and their respective IMOD alignment transformations from the full tomogram metadata. Tomograms were denoised with Isonet2, trained on even-odd tomograms, with a B-factor set to 400. Denoised and deconvolved tomograms were inspected with 3dmod in the IMOD package to annotate filament-containing tomograms, but all measurements reported (including filament crossover, widths, and lengths) were done on the unfiltered tomograms.

**Subtomogram averaging**

An integrated workflow combining WarpTools, Dynamo, and RELION-5 was employed for helical reconstruction, following published protocols (*118*–*121*). Only FTLD-*GRN* patient-derived lysosome tomograms with a pixel size of 8.48 Å were imported into Dynamo (version 1.1.532) to trace filaments using a torsion model, picking the extreme ends of relatively straight filaments and grouping them based on morphology. Filament traces were resampled at 16.96 Å intervals for particle extraction (box size of 64 pixels), resulting in 25,508 and 7,703 total subtomograms being picked for singlet and doublet filaments, respectively. An initial template was created by averaging a random subset of 500 particles. One round of alignment was conducted with 4 iterations, a limited search cone of 60°, allowed azimuthal rotation of up to 180º, and limited particle shift (4 pixels in any direction). Coordinates and Euler angles of aligned particles were converted into warp Self-defining Text Archiving and Retrieval (STAR) format using dynamo2warp functionality from the dynamo2m package (<https://github.com/alisterburt/dynamo2m>). The filament number (_rlnHelicalTubeID) was added using the Starparser package (<https://github.com/sami-chaaban/starparser>). Dynamo-aligned subtomogram volumes were then re-extracted as 2D particle image series in WarpTools, either at 8.48 Å and a box size of 96 pixels or at 4.24 Å and a box size of 192 pixels, for helical reconstruction in RELION-5. Initial reference models were either imported from Dynamo using *relion_image_handler* or generated as a featureless Gaussian tube using *relion_helix_toolbox*.

Subsequently, for the TMEM106B singlet, particles were refined and classified to remove junk, with C1 symmetry with a helical twist of -0.41°, and a rise of 4.8 Å. A set of 4,848 particles, as judged by the rough 3D shape, was used for final refinement. For TMEM106B doublets, refinements were performed with C2 symmetry, a helical twist of -0.41°, and a rise of 4.8 Å; the final refinement contained 7,198 particles.

The final reconstructions were sharpened using the standard post-processing procedures in RELION-5, and resolution was estimated by Gold-standard Fourier shell correlation (cutoff at 0.143) between the two independently refined half-maps, using phase randomization to correct for convolution effects introduced by a generous soft-edged solvent mask.

TMEM106B structures of the doublet and singlet determined from sarkosyl-insoluble patient-derived brain material (PDB: 7SAR and 7SAQ, respectively) were fit as rigid bodies in the density maps obtained from subtomogram averaging in ChimeraX v1.11 (*122*).

**Segmentation**

The v10_alpha and v_10_beta_FAaug pre-trained models of MemBrain v2 were used to help detect organelle membranes in reconstructed tomograms (*123*). Subsequently, the tomograms were manually segmented using Amira (Thermo Fisher Scientific). Thick and thin filaments in iNeuron and mice lysosome tomograms were automatically detected by the convolutional neural network–based segmentation implemented within Amira. A pseudo-TMEM106B fibril model comprised of repeats generated in chimeraX (chains A, B of deposited PDB model 7SAR) with known helical symmetry parameters (rise [4.78 Å], twist [-0.41°], and resolution filtered to 8 Å), were manually positioned into human lysosome tomograms within Amira (Thermo Fisher Scientific) to occupy assigned TMEM106B density within tomograms. Segmented features were exported as Wavefront .obj files for visualization in UCSF ChimeraX. Figures and movies were produced using UCSF ChimeraX.
