## Supplemental Tables S1-S10 for "Neurodegeneration risk variants promote lysosomal TMEM106B fibril accumulation"

| SNP | Protein | N | Beta | SE | CI_lo | CI_hi | P |
| --- | --- | --- | --- | --- | --- | --- | --- |
| rs5848 | GRN | 1178 | -0.2886 | 0.0439 | -0.3746 | -0.2026 | 7.33E-11 |
| rs5848 | TMEM106B | 1365 | 0.336 | 0.0405 | 0.2566 | 0.4154 | 2.58E-16 |
| rs3173615 | GRN | 1182 | -0.0895 | 0.0417 | -0.1712 | -0.0078 | 0.032 |
| rs3173615 | TMEM106B | 1369 | -0.4944 | 0.0363 | -0.5655 | -0.4233 | 9.15E-40 |

**Table S1: TMEM106B and GRN pQTL statistics on TMEM106B and GRN protein.**

| **Variable** | **Overall** | **C/C** | **C/G** | **G/G** | **P value** |
| --- | --- | --- | --- | --- | --- |
| N | 723 | 218 | 359 | 146 |  |
| Age at death (years) | 88.89 (6.70) | 89.25 (6.63) | 88.82 (6.86) | 88.52 (6.43) | 0.343 |
| Post-mortem interval (hours) | 8.22 (6.12) | 7.97 (4.90) | 7.82 (6.12) | 9.59 (7.47) | 0.000871 |
| Education (years) | 15.65 (3.52) | 15.70 (3.25) | 15.81 (3.67) | 15.20 (3.53) | 0.195 |
| Braak stage | 3.69 (1.23) | 3.64 (1.20) | 3.62 (1.27) | 3.93 (1.15) | 0.0241 |
| CERAD (standard) | 1.87 (1.11) | 1.82 (1.16) | 1.85 (1.09) | 2.01 (1.08) | 0.208 |
| Amyloid | 4.99 (4.55) | 4.73 (4.53) | 4.90 (4.46) | 5.65 (4.78) | 0.128 |
| NFT | 0.72 (0.84) | 0.69 (0.85) | 0.69 (0.79) | 0.87 (0.91) | 0.0569 |
| Tangles | 7.10 (8.27) | 6.53 (7.66) | 7.00 (8.31) | 8.29 (9.00) | 0.0827 |
| APOE4 dose | 0.30 (0.50) | 0.28 (0.49) | 0.28 (0.48) | 0.39 (0.56) | 0.0743 |
| TDP stage (tdp_st4) | 1.00 (1.20) | 1.14 (1.25) | 0.98 (1.20) | 0.85 (1.10) | 0.128 |
| TDP score (6-region) | 0.60 (0.93) | 0.63 (0.91) | 0.63 (0.98) | 0.48 (0.83) | 0.324 |
| Female (%) | 491 (67.9%) | 154 (70.6%) | 240 (66.9%) | 97 (66.4%) | 0.584 |
| Clinical group: AD | 266 (36.8%) | 87 (39.9%) | 133 (37.0%) | 46 (31.5%) | 0.106 |
| Clinical group: AsymAD | 292 (40.4%) | 76 (34.9%) | 144 (40.1%) | 72 (49.3%) |  |
| Clinical group: Control | 165 (22.8%) | 55 (25.2%) | 82 (22.8%) | 28 (19.2%) |  |
| Ancestry: AA | 77 (10.7%) | 11 (5.0%) | 39 (10.9%) | 27 (18.5%) | 0.00181 |
| Ancestry: EUR | 643 (88.9%) | 206 (94.5%) | 319 (88.9%) | 118 (80.8%) |  |
| Ancestry: NA | 3 (0.4%) | 1 (0.5%) | 1 (0.3%) | 1 (0.7%) |  |
| APOE ε4 carrier: ≥1 ε4 | 202 (27.9%) | 56 (25.7%) | 95 (26.5%) | 51 (34.9%) | 0.0878 |

**Table S2: Cohort for QTL analyses in Figure 1 stratified by TMEM106B rs3173615 genotype.** Comparison of continuous variables: mean (SD) with Kruskal–Wallis P value across groups. Comparison of categorical variables: n (% within column) with chi-square P value across groups. APOE shown as ε4 carrier status (≥1 ε4 vs 0 ε4).

| SNP | Peptide | Position | N | Beta | SE | CI_lo | CI_hi | P |
| --- | --- | --- | --- | --- | --- | --- | --- | --- |
| rs3173615 | EDAYDGVTSENMR | 15-27 | 636 | 0.0104 | 0.0076 | -0.0045 | 0.0253 | 0.1708 |
| rs3173615 | NGDVSQFPYVEFTGR | 42-56 | 603 | 0.0063 | 0.0058 | -0.0051 | 0.0176 | 0.279 |
| rs3173615 | DSVTCPTCQGTGR | 57-69 | 474 | 0.0131 | 0.0213 | -0.0286 | 0.0547 | 0.5392 |
| rs3173615 | GQENQLVALIPYSDQR | 73-88 | 723 | -0.0051 | 0.0076 | -0.02 | 0.0099 | 0.508 |
| rs3173615 | SAYVSYDVQK | 130-139 | 772 | -0.2577 | 0.026 | -0.3086 | -0.2068 | 6.91E-22 |
| rs3173615 | YQYVDCGR | 248-255 | 769 | -0.0608 | 0.0175 | -0.095 | -0.0266 | 5.29E-04 |
| rs5848 | EDAYDGVTSENMR | 15-27 | 633 | -0.0015 | 0.0083 | -0.0178 | 0.0148 | 0.857 |
| rs5848 | NGDVSQFPYVEFTGR | 42-56 | 600 | -0.0103 | 0.0062 | -0.0224 | 0.0019 | 0.0983 |
| rs5848 | DSVTCPTCQGTGR | 57-69 | 471 | -0.0253 | 0.0229 | -0.0702 | 0.0195 | 0.2693 |
| rs5848 | GQENQLVALIPYSDQR | 73-88 | 719 | 0.0102 | 0.0084 | -0.0062 | 0.0266 | 0.2224 |
| rs5848 | SAYVSYDVQK | 130-139 | 768 | 0.1891 | 0.0295 | 0.1313 | 0.247 | 2.63E-10 |
| rs5848 | YQYVDCGR | 248-255 | 765 | 0.1345 | 0.0187 | 0.0978 | 0.1712 | 1.68E-12 |

**Table S3: TMEM106B and GRN pQTL on TMEM106B peptides.**

| **Variable** | **Overall** | **CC** | **CT** | **TT** | **P value** |
| --- | --- | --- | --- | --- | --- |
| N | 719 | 333 | 296 | 90 |  |
| Age at death (years) | 88.89 (6.70) | 89.35 (6.05) | 88.78 (7.15) | 87.57 (7.28) | 0.113 |
| Post-mortem interval (hours) | 8.21 (6.12) | 8.01 (6.51) | 8.28 (5.13) | 8.69 (7.54) | 0.0703 |
| Braak stage | 3.69 (1.22) | 3.74 (1.19) | 3.59 (1.26) | 3.84 (1.20) | 0.183 |
| CERAD (standard) | 1.88 (1.11) | 1.90 (1.09) | 1.82 (1.14) | 1.97 (1.06) | 0.612 |
| Amyloid | 5.01 (4.56) | 5.11 (4.69) | 4.98 (4.57) | 4.66 (3.91) | 0.894 |
| NFT | 0.72 (0.84) | 0.74 (0.85) | 0.71 (0.83) | 0.68 (0.80) | 0.407 |
| Tangles | 7.09 (8.25) | 7.14 (8.68) | 6.91 (7.75) | 7.52 (8.24) | 0.651 |
| APOE4 dose | 0.30 (0.50) | 0.28 (0.49) | 0.33 (0.51) | 0.31 (0.51) | 0.373 |
| TDP stage (tdp_st4) | 1.00 (1.20) | 0.92 (1.13) | 1.05 (1.25) | 1.14 (1.27) | 0.412 |
| TDP score (6-region) | 0.60 (0.93) | 0.55 (0.88) | 0.60 (0.93) | 0.79 (1.11) | 0.403 |
| Female (%) | 489 (68.0%) | 217 (65.2%) | 214 (72.3%) | 58 (64.4%) | 0.119 |
| Clinical group: AD | 265 (36.9%) | 132 (39.6%) | 96 (32.4%) | 37 (41.1%) | 0.245 |
| Clinical group: AsymAD | 291 (40.5%) | 130 (39.0%) | 124 (41.9%) | 37 (41.1%) |  |
| Clinical group: Control | 163 (22.7%) | 71 (21.3%) | 76 (25.7%) | 16 (17.8%) |  |
| Ancestry: AA | 77 (10.7%) | 11 (3.3%) | 36 (12.2%) | 30 (33.3%) | 5.05e-14 |
| Ancestry: EUR | 639 (88.9%) | 320 (96.1%) | 259 (87.5%) | 60 (66.7%) |  |
| Ancestry: NA | 3 (0.4%) | 2 (0.6%) | 1 (0.3%) | 0 (0.0%) |  |
| APOE ε4 carrier: ≥1 ε4 | 201 (28.0%) | 86 (25.8%) | 90 (30.4%) | 25 (27.8%) | 0.367 |

**Table S4: Cohort for QTL analyses in Figure 4C stratified by GRN rs5848 genotype.** Comparison of continuous variables: mean (SD) with Kruskal–Wallis P value across groups. Comparison of categorical variables: n (% within column) with chi-square P value across groups. APOE shown as ε4 carrier status (≥1 ε4 vs 0 ε4).

| **Protein** | **Disease / tissue source** | **Fibril width (nm)** | **Crossover distance (nm)** | **Primary cryo-EM reference** | **PMID** |
| --- | --- | --- | --- | --- | --- |
| Tau (PHF/SF) | Alzheimer’s disease brain | ~10–20 | ~65–80 | Fitzpatrick et al., *Nature* (2017) | 28678775 |
| Tau | Pick’s disease brain | ~10–15 | ~70–100 | Falcon et al., *Nature* (2018) | 30158706 |
| Tau | CTE brain | ~15–20 | ~70 | Falcon et al., *Nature* (2019) | 30894745 |
| Tau | Corticobasal degeneration brain | ~10–20 | ~80–150 | Zhang et al., *Nature* (2020) | 32050258 |
| Amyloid-β | Alzheimer’s plaques | ~6–12 | ~120–160 | Kollmer et al., *Nat Commun* (2019) | 31664019 |
| Amyloid-β (Aβ42) | Alzheimer’s brain | ~7–12 | ~100–160 | Yang et al., *Science* (2022) | 35025654 |
| α-Synuclein | Multiple system atrophy brain | ~10–18 | ~100–150 | Schweighauser et al., *Nature* (2020) | 32461689 |
| TDP-43 | ALS/FTLD brain inclusions | ~10–20 filament | long/irregular | Arseni et al., *Nature* (2022) | 34880495 |
|  |  | (~5 nm for the fibril core) |  |  |  |
| TMEM106B (singlet) | FTLD-TDP brain | ~12–13 | ~200–220 | Jiang et al., *Nature* (2022) | 35344984 |
| TMEM106B (doublet) | Neurodegenerative brains | ~24–26 | ~210 | Chang et al., *Cell* (2022) | 35247328 |
| PrPSc | Prion disease brain | ~10–20 | ~50–120 | Kraus et al., *Mol Cell.* (2021) | 34433091 |
| TAF15 | FTLD-FET brain | ~8 | ~40 | Tetter et al., *Nature* (2024) | 38057661 |
| SOD1* | Recombinantly produced | ~7–12 | variable | Wang et al., *Nat Commun* (2020) | 35715417 |
| FUS* | Recombinant disease-relevant fibrils | ~8–12 | variable | Sun et al., *iScience* (2021) | 35036880 |

**Table S5: Known widths and crossover distances of published fibril structures.**

| PG Protein Groups | PG Genes | PG Protein Descriptions | PEP Stripped Sequence | EG Protein PTM Locations | T1/2 R16 Rep 1 | T1/2 R16 Rep 2 | T1/2 R16 Rep 3 |
| --- | --- | --- | --- | --- | --- | --- | --- |
| Q00000 | TMEM106B_Svariant | Transmembrane protein 106B S-variant | LNNISIIGPLDMK | (N182,N183) |  |  |  |
| Q00000 | TMEM106B_Svariant | Transmembrane protein 106B S-variant | LNNISIIGPLDMK | (N183,M192) | 12.7 | 13.4 | 13.7 |
| Q00000 | TMEM106B_Svariant | Transmembrane protein 106B S-variant | LNNISIIGPLDMK | (N183) | 8.6 | 8.8 | 8.6 |
| Q00000 | TMEM106B_Svariant | Transmembrane protein 106B S-variant | LNNISIIGPLDMK |  | 8.9 | 9 | 8.3 |
| Q9NUM4 | TMEM106B | Transmembrane protein 106B | LNNITIIGPLDMK | (N183,M192) |  |  |  |
| Q9NUM4 | TMEM106B | Transmembrane protein 106B | EDAYDGVTSENMR | (M26) | 1.3 | 1.6 | 1.7 |
| Q9NUM4 | TMEM106B | Transmembrane protein 106B | LNNITIIGPLDMK | (N182,N183) |  |  |  |
| Q9NUM4 | TMEM106B | Transmembrane protein 106B | LNNITIIGPLDMK | (N183) |  |  |  |
| Q9NUM4 | TMEM106B | Transmembrane protein 106B | LNNITIIGPLDMK |  |  |  |  |
| Q9NUM4 | TMEM106B | Transmembrane protein 106B | YQYVDCGR | (Q249,C253) |  |  |  |
| Q9NUM4 | TMEM106B | Transmembrane protein 106B | GQENQLVALIPYSDQR | (Q77) | 1.8 | 1.8 | 1.7 |
| Q9NUM4 | TMEM106B | Transmembrane protein 106B | NGLVNSEVHNEDGR | (N28) | 1.2 | 1.1 | 1.1 |
| Q9NUM4 | TMEM106B | Transmembrane protein 106B | SAYVSYDVQKR |  | 7.7 | 7.6 | 6.2 |
| Q9NUM4 | TMEM106B | Transmembrane protein 106B | NGDVSQFPYVEFTGR | (N42) | 1.3 | 1.4 | 1.3 |
| Q9NUM4 | TMEM106B | Transmembrane protein 106B | NGLVNSEVHNEDGR |  | 1 | 1 | 1 |
| Q9NUM4 | TMEM106B | Transmembrane protein 106B | EDAYDGVTSENMR |  | 1.2 | 1.3 | 1.1 |
| Q9NUM4 | TMEM106B | Transmembrane protein 106B | GQENQLVALIPYSDQR |  | 1.8 | 1.8 | 1.7 |
| Q9NUM4 | TMEM106B | Transmembrane protein 106B | NGDVSQFPYVEFTGR |  | 1.3 | 1.3 | 1.3 |
| Q9NUM4 | TMEM106B | Transmembrane protein 106B | SAYVSYDVQK |  | 8.9 | 7.5 | 8.1 |
| Q9NUM4 | TMEM106B | Transmembrane protein 106B | SLSHLPLHSSK |  | 1.4 | 1.4 | 1.3 |
| Q9NUM4 | TMEM106B | Transmembrane protein 106B | YQYVDCGR | (C253) | 6.7 | 7.6 | 7.3 |

**Table S6: Half-Life by peptide from TMEM106B overexpressing iNeurons.**

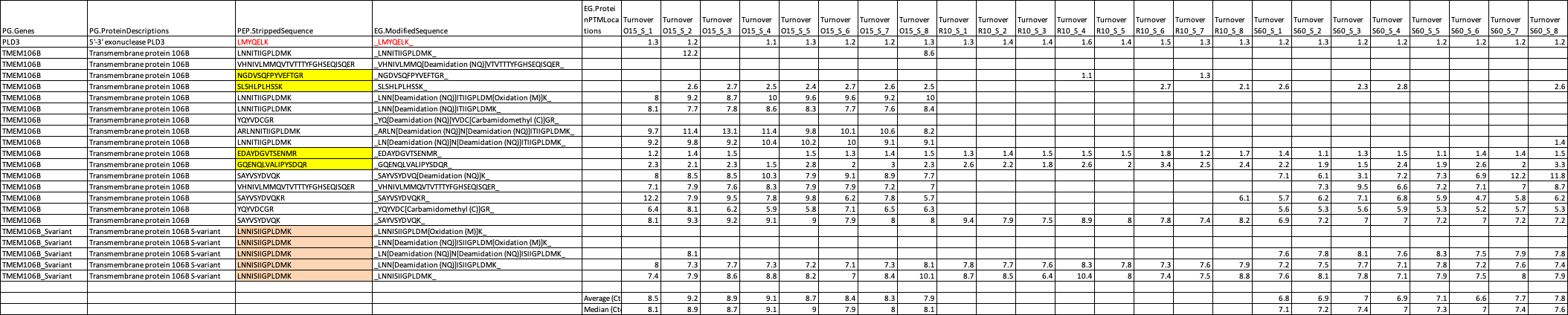

**Table S7: PLD3-TMEM106BFC half-life data by peptide (O15 - T185 variant; S60 - S185 variant; R10 - PLD3-control containing peptides from endogenous protein).** Yellow-highlighted peptides are from the N-terminus of native TMEM106B in the cells, not PLD3-TMEM106B. Orange-highlighted peptides are from the S-variant of TMEM106B (thus, from endogenous TMEM106B) not of 'O15'- the T variant of PLD3-TMEM106B.

| Target | Source | Catalog # or ID | Species | Figs and Panels | Western blot conc. | IHC/IF Conc. | Citation |
| --- | --- | --- | --- | --- | --- | --- | --- |
| N-terminal TMEM106B | CST | E7H7Z #93334 | Rabbit | fig. 1, 4, s6 | 1:1000 |  |  |
| Cleaved TMEM106B(FC) | CST | F9W60 #87145 | Rabbit | fig. 1, s1, 4, s6 | 1:1000 | 1:100 |  |
| C-terminus of TMEM106B | Petrucelli Lab | LP26.2.3 | Rabbit | 2b, s2a, s2b, s2d, s6 | 1:1000 | 1:1000 | PMID: 38232138 |
| C-terminus of TMEM106B | SySy | 506 017 | Rat | fig. 2c | 1:1000 |  | PMID: 39709600 |
| C-terminal Tail TMEM106B (254-274) | gift from Anja Capell |  | Rat | fig. s2c | 1:1000 |  |  |
| N-terminus of PLD3 | gift from Markus Damme | | Rabbit | fig. 2, s6 | 1:2000 |  | PMID: 29386126 |
| PGRN | R&D Systems | AF2420 | Goat | 4, s6 | 1:200-250 | 1:1000 |  |
| GAPDH | Genetex | GT239 | Mouse |  | 1:5000 |  |  |
| LAMP1 | DSHB | H4A3 | Mouse | fig. s2 |  | 1:1000 |  |
| pTDP-43 (Ser409/410) | Petrucelli Lab | 3655 | Rabbit | Fig 4J |  | 1:50 | PMIDs: 25977373; 30767771 |
| p62 | CST | 23214 | Rabbit | Fig 4F |  | 1:50 |  |
| CD68 | abcam | ab125212 | Rabbit | Fig 4H |  | 1:100 |  |
| Iba1 | MilliporeSigma | MABN92 | Mouse | fig s5c |  | 1:50 |  |
| GFAP | Santa Cruz | sc-58766 | Mouse | fig s5c |  | 1:200 |  |
| MAP2 | ThermoFisher | PA1-10005 | Chicken | fig s5c |  | 1:2000 |  |
| GAPDH | Meridian Life Science | H86504M | Mouse | fig s5b |  | 1:5000 |  |
| Cathepsin D | R&D Systems | AF1029 | Goat | fig s5c |  | 1:200 |  |
| Cathepsin D | R&D Systems | AF1014 | Goat | Fig. 2A |  | 1:100 |  |
| Iba1 | Wako/FujiFilm | 019-19741 | Rat | fig. s5d |  | 1:3000 |  |
| GFAP | BioGenex | PU020-UP | Rabbit | fig. s5e |  | 1:2500 |  |

**Table S8: Antibodies and Concentrations.**

| Table S9 | coding sequence |
| --- | --- |
| PLD3- Control | atgaagcctaaactgatgtaccaggagctgaaggtgcctgcagaggagcccgccaatgagctgcccatgaatgagattgaggcgtggaaggctgcggaaaagaaagcccgctgggtcctgctggtcctcattctggcggttgtgggcttcggagccctgatgactcagctgtttctatgggaatacggcgacttgcatctctttgggcccaaccagcgcggcagcggcagcggctaa |
| PLD3-TMEM106B (T185) | atgaagcctaaactgatgtaccaggagctgaaggtgcctgcagaggagcccgccaatgagctgcccatgaatgagattgaggcgtggaaggctgcggaaaagaaagcccgctgggtcctgctggtcctcattctggcggttgtgggcttcggagccctgatgactcagctgtttctatgggaatacggcgacttgcatctctttgggcccaaccagcgcggcagcggcagcggctctatcgacgtgaaatacattggtgtaaaatcagcctatgtcagttatgatgttcagaagcgtacaatttatttaaatatcacaaacacactaaatataacaaacaataactattactctgtcgaagttgaaaacatcactgcccaagttcaattttcaaaaacagttattggaaaggcacgcttaaacaacataaccattattggtccacttgatatgaaacaaattgattacacagtacctaccgttatagcagaggaaatgagttatatgtatgatttctgtactctgatatccatcaaagtgcataacatagtactcatgatgcaagttactgtgacaacaacatactttggccactctgaacagatatcccaggagaggtatcagtatgtcgactgtggaagaaacacaacttatcagttggggcagtctgaatatttaaatgtacttcagccacaacagtaa |
| PLD3-TMEM106B (S185) | atgaagcctaaactgatgtaccaggagctgaaggtgcctgcagaggagcccgccaatgagctgcccatgaatgagattgaggcgtggaaggctgcggaaaagaaagcccgctgggtcctgctggtcctcattctggcggttgtgggcttcggagccctgatgactcagctgtttctatgggaatacggcgacttgcatctctttgggcccaaccagcgcggcagcggcagcggctctatcgacgtgaaatacattggtgtaaaatcagcctatgtcagttatgatgttcagaagcgtacaatttatttaaatatcacaaacacactaaatataacaaacaataactattactctgtcgaagttgaaaacatcactgcccaagttcaattttcaaaaacagttattggaaaggcacgcttaaacaacataagcattattggtccacttgatatgaaacaaattgattacacagtacctaccgttatagcagaggaaatgagttatatgtatgatttctgtactctgatatccatcaaagtgcataacatagtactcatgatgcaagttactgtgacaacaacatactttggccactctgaacagatatcccaggagaggtatcagtatgtcgactgtggaagaaacacaacttatcagttggggcagtctgaatatttaaatgtacttcagccacaacagtaa |

**Table S9: DNA coding sequences of PLD3-control and PLD3-TMEM106B variants.**

| **Case** | ***TMEM106B* rs3173615** | **Age at death (yrs)** | **Sex** | **Braak** | **Thal** | **PMI (hr)** |
| --- | --- | --- | --- | --- | --- | --- |
| Sporadic FTLD-TDP Type A | C/C (T185T) | 77.37 | M | 1 | 0 | - |
| *GRN* FTLD-TDP Type A | C/C (T185T) | 72.18 | M | 0 | 2 | 4 |
| Control | C/C (T185T) | 80 | M |  |  | - |

**Table S10: Information of cases used for cryo-ET.**
